# Multidimensional single-cell benchmarking of inducible promoters for precise dynamic control in budding yeast

**DOI:** 10.1101/2020.08.16.253310

**Authors:** Vojislav Gligorovski, Ahmad Sadeghi, Sahand Jamal Rahi

## Abstract

For quantitative systems biology, simultaneous readout of multiple cellular processes as well as precise, independent control over different genes’ activities are essential. In contrast to readout systems such as fluorescent proteins, control systems such as inducible transcription-factor-promoter systems have only been characterized in an *ad hoc* fashion, impeding precise system-level manipulations of biological systems and reliable modeling.

We designed and performed systematic benchmarks involving easy-to-communicate units to characterize and compare inducible transcriptional systems. We built a comprehensive single-copy library of inducible systems controlling standardized fluorescent protein expression in budding yeast, including *GAL1pr*, *GALL*, *MET3pr*, *CUP1pr*, *PHO5pr*, *tetOpr*, *terminator*-*tetOpr*, Z_3_EV system, the blue-light optogenetic systems El222*-LIP*, El222*-GLIP* and the red-light inducible PhyB-PIF3 system. To analyze these systems’ dynamic properties, we performed high-throughput time_-_lapse microscopy. The analysis of >100 000 cell images was made possible by the recently developed convolutional neural network YeaZ. We report key kinetic parameters, scaling of noise levels, impacts on growth, and, crucially, the fundamental leakiness of each system. Our multidimensional benchmarking additionally uncovers unexpected disadvantages of widely used tools, e.g., nonmonotonic activity of the *MET3* and *GALL* promoters, slow off kinetics of the doxycycline and estradiol-inducible systems *tetOpr* and Z_3_EV, and high variability of *PHO5pr* and red-light activated PhyB-PIF3 system. We introduce two new tools for controlling gene expression: strongLOV, a more light-sensitive El222 mutant, and *ARG3pr* that functions as an OR gate induced by the lack of arginine or presence of methionine. To demonstrate the ability to finely control genetic circuits, we experimentally tuned the time between cell cycle Start and mitotic entry in budding yeast, artificially simulating near-wild-type timing.

The characterizations presented here define the compromises that need to be made for quantitative experiments in systems and synthetic biology. To calibrate perturbations across laboratories and to allow new inducible systems to be benchmarked, we deposited single-copy reporter yeast strains, plasmids, and computer analysis code in public repositories. Furthermore, this resource can be accessed and expanded through the website https://promoter-benchmark.epfl.ch/.

## Introduction

Control over the level and timing of gene activity does not only offer advantages over more traditional genetics approaches such as gene knockouts or constitutive overexpression but is indispensable for many applications. In particular, understanding system-level properties and constructing artificial cellular behaviors frequently require the independent, temporally precise, and reversible manipulation of different nodes in a genetic network. As a result, inducible expression systems and their characterization are critical for advances in systems and synthetic biology.

Inducible systems are widely used in systems biology for studying the dynamics, topology, and stochasticity of genetic networks.^1–3^ For example, 1 sec long pulses of light were used to recruit the proteins that control the site of budding^4^; 5 min galactose induction was used to express double-strand DNA break-inducing endonucleases.^5^ In metabolic engineering, inducible systems are employed for the reversible activation of biosynthetic pathways at specific stages of growth or for fine-tuning activation levels.^6–8^ Reversible activation of gene activity is also needed in synthetic biology for the construction of switchable logic circuits^9,10^ or to reduce the toxic effects of specific gene products^11^.

Exogenous regulation of gene expression in eukaryotes can in principle be introduced at different stages, the transcriptional or translational level as well as at the posttranslational level by controlling protein-protein interactions or protein degradation.^12,13^ In *Saccharomyces cerevisiae*, a widely used organism in research and industry, the most common way of tuning the level of gene expression is by regulating transcription.^14^ Moreover, the majority of the tools for manipulating gene expression have been engineered for yeasts.^15^ This is why we focus on inducible transcriptional systems here.

Many commonly used inducible transcriptional systems in budding yeast are regulated by small metabolites such as galactose, methionine, or copper^16^. Using nutrients to control gene expression has the advantage that the relevant transcription factors are already present in cells and have been fine-tuned over the course of the evolution. On the other hand, the drawback is that changes in nutrient levels generally also affect metabolism. To avoid this, synthetic systems have been created which respond to compounds not naturally present in the host. In addition to tetracycline-regulated transcription factors^17^, several systems that are estradiol-inducible have been constructed for budding yeast^18–20^, such as the Z_3_EV system. While synthetic systems are usually orthogonal to cell physiology, they can nevertheless have an effect on cellular growth, for example, due to the toxicity of the inducer. More recently, light sensors from bacteria and plants have been adapted for use as transcriptional control systems in budding yeast^21^. In contrast to the other systems for manipulating cellular processes, light provides a rapid, noninvasive, and convenient means of control^22^.

For precise control of gene activity, inducible systems should ideally have fast kinetics, high dynamic range, low basal activity (leakiness), and low noise. Leakiness is a poorly characterized but crucial property since for many applications it is essential to be able to turn expression truly ‘off’. Leakiness is particularly important when controlling genes that are toxic or cause changes to the genome such as Cas9^23^, Cre-loxP^11^, or Ho^24^ endonuclease. However, for inducible systems, most of these properties have either not been assessed precisely, not in a manner that would allow their direct comparison, or have not been determined at all. Although new inducible systems are being developed,^18,19,25–28^ a standard benchmark for rigorous evaluation of their properties does not exist. Due to the absence of standardized quantitative descriptions, the choice of inducible systems is usually guided by intuition or time-consuming trial and error. The lack of such benchmarks for controlling cellular behavior stands in contrast to existing thorough characterizations of readout systems such as fluorescent proteins.^30–33^

There are multiple technical challenges for characterizing inducible systems quantitatively:

1. Single-cell time courses need to be recorded by fluorescence microscopy and analyzed. For this to be feasible with sufficient numbers of cells, a highly efficient and accurate segmentation method such as the newly developed convolutional neural network YeaZ^34^ was needed, which we used to analyze >100000 yeast cell images. Population snapshots by flow cytometry do not suffice for reconstructing single-cell time courses unambiguously. Moreover, flow cytometry has substantially higher levels of measurement noise and thus overestimates the true expression stochasticity^35^ compared to fluorescence microscopy (Fig. 1).
2. To allow comparisons, all reporters for the inducible systems must be designed uniformly, e.g., introduced at the same genomic locus and in the same number of copies. Here, we ensure that each reporter is introduced as a single copy at the same locus (*URA3*).
3. Additionally, to measure absolute characteristics, the absolute copy numbers of the reporter systems in each cell must be fixed. Our single-copy reporters allow us to measure the fundamental characteristics of the inducible transcriptional system such as their minimal leakiness.
4. Fluorescence levels are often reported in ‘arbitrary units’ (*AU*), which differ among fluorescent proteins and microscopes, making measurements difficult to compare between different laboratories. To overcome this limitation, we calibrated all fluorescence units to an easy-to-communicate reference unit, “peak *GAL1pr* expression” (maxGAL1), whose intuitiveness makes it appealing as a practical unit for measuring gene induction levels. Thus, we avoid the difficulty of quantifying expression in terms of absolute protein numbers but instead normalize all levels to a very well-known expression system, which could therefore serve as a universal expression unit.

**Figure 1.**
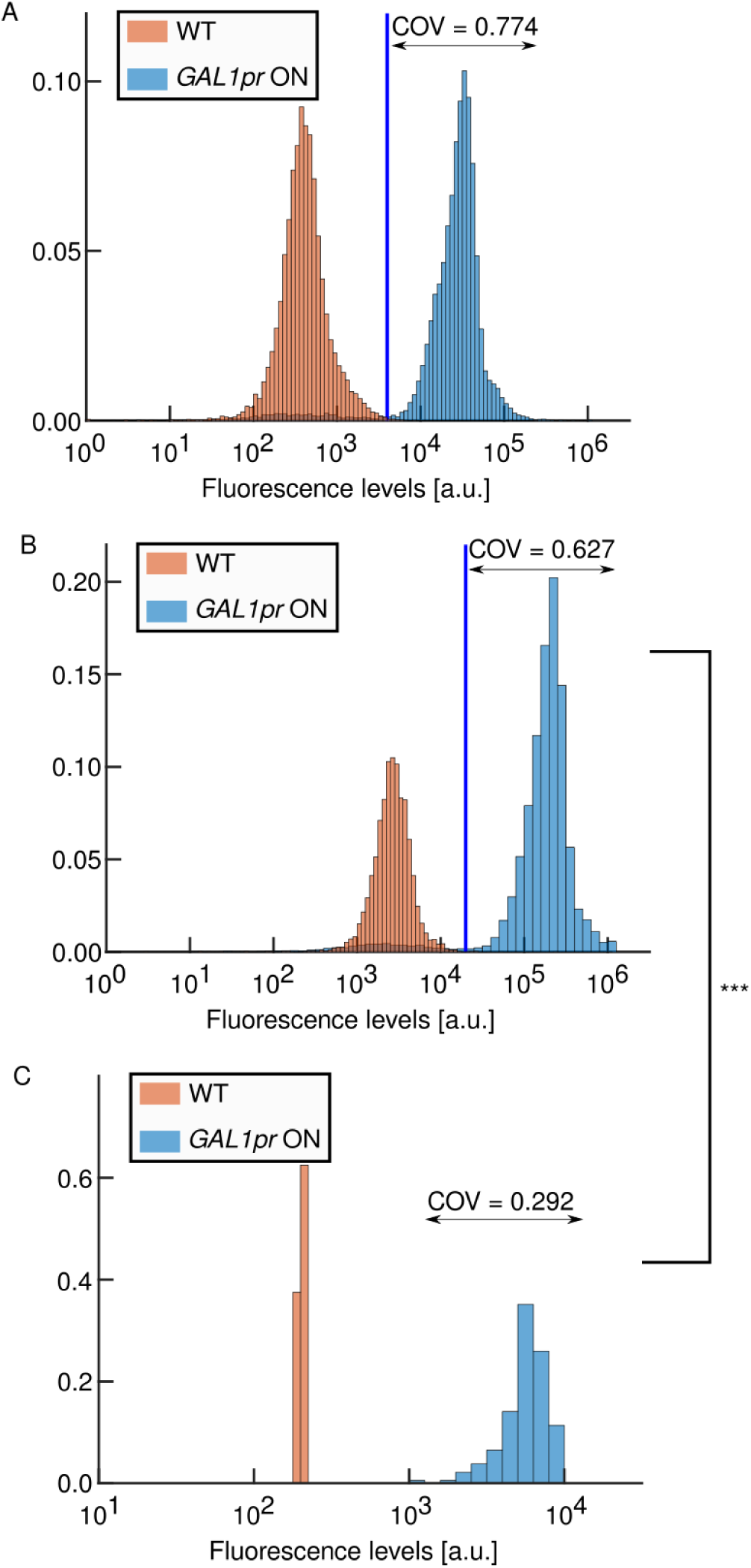
Measurement noise is substantially higher in flow cytometry compared to fluorescence microscopy. A: Flow cytometry measurements of cells with a *GAL1pr-yEVenus-PEST* construct and wild-type control cells in inducing galactose medium (COV: coefficient of variation). B: Increasing the excitation laser intensity even up to the saturation point of the sensor (2^20^ ≈ 1.05·10^6^) does not substantially reduce the COV. C: Fluorescence microscopy measurements of the same cells. COV is calculated for all cells with the *GAL1pr-yEVenus-PEST* construct (no gating was applied). A, B: COV is calculated for the induced *GAL1pr-yEVenus-PEST* population, which is defined by fluorescence values higher than the threshold indicated by the blue vertical line. A, B, C: Data is shown on a logarithmic scale but the COV is computed based on non-transformed values. B, C: p = 7.8e-15 (one-tailed z-test for significance of COV differences between induced populations in B and C panels).

Here, we present:

1. a single-cell based characterization of new as well as widely used inducible systems, identifying several noteworthy features of these systems,
2. the extraction of key parameters from the time courses: on time lag, off time lag, induction speed and strength, noise levels, and leakiness,
3. their population-level averages and cell-to-cell variability,
4. strongLOV, a new mutant of the light-inducible transcription factor El222 with higher light sensitivity,
5. an analysis of the *ARG3* promoter for potential use as an inducible system,
6. a demonstration of how these data enable fine experimental tuning by timing successive cell-cycle transitions with close-to-wild-type timing,
7. computer code, budding yeast strains, and plasmids to allow the benchmark to be applied to future inducible systems and to calibrate measurements across laboratories, and
8. the website promoter-benchmark.epfl.ch to make the benchmark extendable and all data easily accessible.

## Results

### Selection of inducible transcriptional systems

The galactose regulon has been utilized for many decades to control gene expression with *GAL1pr* potentially being the most widely used inducible promoter in budding yeast^36,37^. *GAL1pr* is tightly repressed in the presence of glucose by the Gal80 and Mig1 repressors^38^. In the presence of galactose, *GAL1pr* is induced more than 1000-fold^39^. Glucose repression induces transcriptional downregulation of *GAL* regulatory genes. To avoid delays when switching from glucose to galactose, cells are typically grown in non-inducing and non-repressing raffinose medium. Because *GAL1pr* is too strong for many applications when induced, a weakened version, *GALL*, was developed.^40^

*MET3* was discovered through a screen for methionine auxotrophy in yeast^41^. Met3 is an ATP-sulfurylase which catalyzes the first step in the sulfur assimilation pathway.^42^ Its transcription is strongly repressed in methionine-rich media^42^. It is commonly used to control the expression of budding yeast genes whose transcription levels are lower than *GAL1pr*. For example, the continuous expression of G1/S budding yeast cyclin *CLN2*, whose promoter is comparable in strength to *MET3pr*^43^, from a *MET3pr-CLN2* construct in a *cln1,2,3Δ* background causes almost no discernable effects in cell-cycle timing and cellular morphology^44,45^. In contrast, overexpression of *CLN2* from *GAL1pr* in the same genetic background slows down the cell cycle^46^ and in the wild-type background produces cells with elongated buds^47^.

*CUP1* is part of a feedback loop that mediates resistance to copper toxicity. Its transcription increases when cells are exposed to copper (II) ions.^48^ Although *CUP1pr* has been used as a tool for dynamic gene expression control, it has the fundamental drawback of being regulated by copper ions, which are both essential and, if supplied at high concentrations, toxic.^49^

*PHO5pr* is a member of the *PHO* regulon, which has been researched intensively as a model for studying the relationship between chromatin structure and gene expression dynamics^50^. *PHO5pr* is upregulated in response to a lack of inorganic phosphate^51^, which is required for energy and nucleotide metabolism. The *PHO5* promoter becomes fully active after phosphate, including any stored in the vacuole, is used up^52^.

Tetracycline-responsive systems are widely used for controlling transcription. The core of the system, the *tetO* sequence, is controlled by the tTA regulator, which has been identified as a tetracycline-responsive element in bacteria. In its original form, tTA is part of the Tet-Off system that is inhibited by the antibiotic tetracycline or the closely related molecule doxycycline. The mutations that reverse tTA activity with respect to the inducer have been identified. However, this system, called the Tet-On system, exhibits high basal activity in the absence of the inducer^53^.

Several other synthetic systems, which are estradiol-inducible, have also been constructed for budding yeast^18–20^, such as the Z_3_EV system. Z_3_EV is a transcription factor in which the estradiol receptor is fused to the DNA binding domain of the mouse transcription factor Zif268 and the transcriptional activation domain VP16. One of the main advantages of using artificial transcription factors is that they can be designed to recognize comparatively long DNA motifs, thereby reducing off-target binding^20^. While synthetic systems are usually orthogonal to cellular physiology, they can nevertheless have an effect on cellular growth due to off-target effects or the toxicity of the inducer, for example.

The first light-regulated transcriptional system used in budding yeast was derived from a plant phytochrome and consists of the protein PhyB (<u>Phy</u>tochrome <u>B</u>) and its interaction partner, PIF3 (<u>P</u>hytochrome-Interacting <u>F</u>actor <u>3</u>) protein.^54^ While it paved the way toward optogenetic control of cellular processes, PhyB-PIF3 has the disadvantage that it requires the exogenous addition of the chromophore phycocyanobilin (PCB), which is not produced by most eukaryotes other than plants. For transcriptional control, PhyB and PIF3 are fused to the Gal4 transcriptional activation domain and the Gal4 DNA binding domain, respectively^54,55^. When bound to PCB and activated by red light (≈ 650 nm) PhyB binds PIF3, thereby bringing the transcriptional activation and DNA binding domains close and leading to the expression of the *GAL* family of genes such as *GAL1pr*. In the presence of far-red light (≈ 740 nm), PhyB changes conformation again and dissociates from PIF3. Since the spectra of activating and deactivating light overlap, PhyB is maintained in a dynamic equilibrium between the two states whose ratio depends on the wavelengths of the light.^56^ In addition to the disadvantage of requiring exogenous PCB, the system also affects galactose metabolism in budding yeast when used for transcriptional induction with the split Gal4 transcription factor.

A popular system that overcomes some of the limitations of the PhyB-PIF3 system is the blue-light inducible El222 transcription factor. This prokaryotic LOV-domain photosensor has been adapted for use in many organisms such as yeast, zebrafish, and mammalian cell lines^6,57,58^ by fusing it to the transcriptional activation domain VP16 and a nuclear localization sequence. When exposed to blue light (≈ 465 nm), El222 dimerizes and recognizes its binding sites. These binding sites are typically placed upstream of a minimal promoter^6,59^. We refer to the whole promoter, introduced in ref.^57^ as *LIP* (“light-inducible <u>p</u>romoter”). Unlike PhyB, El222 incorporates flavin-mononucleotide as chromophore, which is naturally occurring in budding yeast. A recent version of *LIP* has been built using the *GAL1* promoter with the Gal4 activator binding sites deleted^29^ instead of the minimal promoter, which we refer to as *GLIP* (“<u>G</u>AL1pr-based light-inducible <u>p</u>romoter”).

### Construction of the *promoter-yEVenus-PEST* library

In order to characterize the inducible systems in a systematic and comprehensive manner, we constructed a library of promoters driving the expression of yEVenus^60^, a bright and fast-folding^30^ yellow fluorescent protein optimized for expression in budding yeast. For a fast-reacting transcriptional reporter (Fig. 2 A), we fused the fluorescent protein to a constitutive degron (PEST) from the *CLN2* gene, which leads to fast degradation of the protein.^61^ The *yEVenus-PEST* construct has been extensively used in the past, including as a transcriptional reporter in budding yeast.^43,62^ In the library, we included *GAL1pr*, *GALL*, *MET3pr*, *CUP1pr*, *PHO5pr*, the synthetic *tetOpr*/Tet-On, and Z_3_EV systems, and the optogenetic systems PhyB-PIF3 and El222 controlling two different promoters, *LIP* and *GLIP*. In addition, we created a new El222 mutant, strongLOV controlling *LIP*, which is introduced in greater detail below.

**Figure 2.**
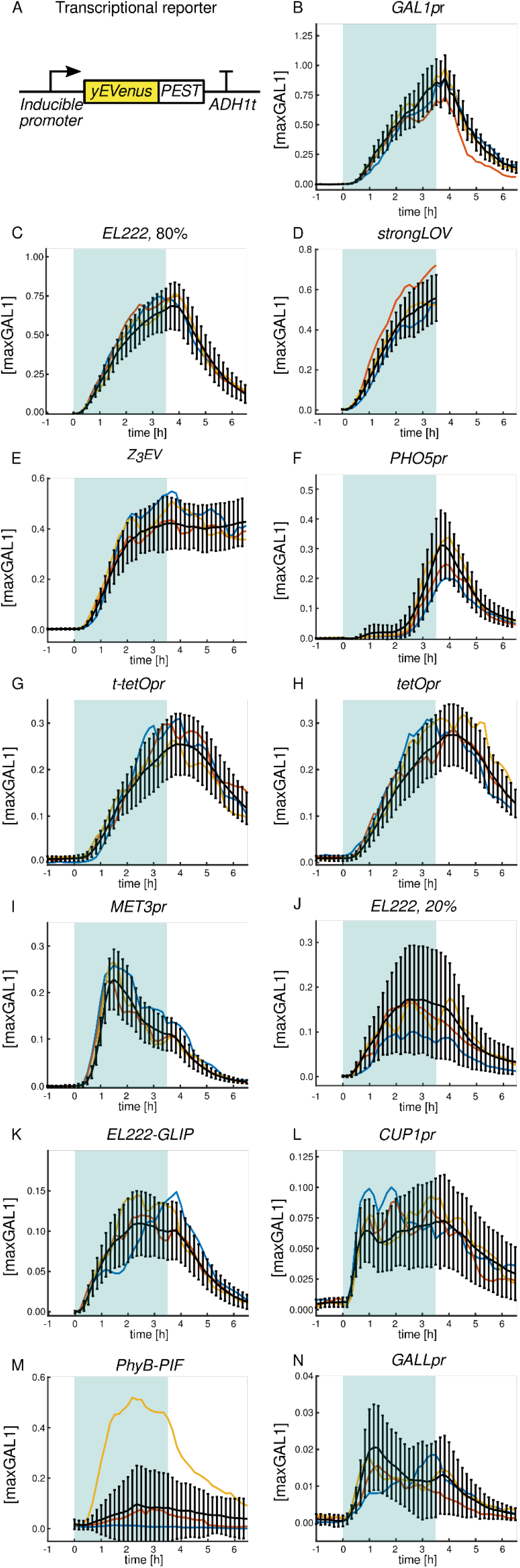
On and off dynamics of inducible systems. A: The reporter for transcriptional activity consists of an inducible promoter and the fast-folding yellow fluorescent protein *yEVenus* gene fused to a constitutive degron (*PEST*) and the *ADH1* terminator. B-N: Time courses of activation and deactivation for different inducible systems sorted in descending order by peak average strength. Induction starts at t = 0 h and finishes at t = 3.5 h. The blue background represents the induction period. Expression is quantified in maxGAL1 units, where 1 maxGAL1 corresponds to steady-state expression of *GAL1pr*. Black lines show the average of the mean cellular expression and standard deviation. Colored lines show different representative single-cell time courses. For the light-inducible systems, fluorescence was not measured prior to induction in order to avoid possible activation by the light source used for fluorescent protein excitation. *EL222* refers to the WT-El222 transcription factor inducing *LIP* under 20% or 80% light intensity, as indicated. *strongLOV* refers to the Glu84Asp El222 mutant introduced in this article inducing *LIP* under 20% light. *GLIP* is induced by El222 under 80% light. Due to the high sensitivity of strongLOV to the excitation light used for the fluorescence measurements, quantification of the off dynamics by microscopy was not possible with our system; turn-off experiments were thus done with samples taken from liquid culture (Fig. 13). Numbers of analyzed cells for each plot are given in Supplementary Table 14.

Several factors such as the genomic integration site^63,64^, the sequence between the promoter and the gene used for cloning^65^, and the terminator sequence^66^ are thought to potentially influence expression in budding yeast. In addition, genetic constructs can in principle be integrated in different copy numbers in the genome, resulting in different levels of expression and noise (Supplementary Fig. 1).^67,68^ To allow direct comparisons between the inducible systems, we built the *promoter-yEVenus-PEST* circuits using the same plasmid backbone sequence and the same cloning strategy and we integrated them as single copies in the same locus (*URA3*) in the genome (Methods).

To prevent transcriptional read-through, some researchers have placed a terminator upstream of the genetic circuit of interest.^1,69–71^ It has been suggested that in yeast, terminators themselves can function as promoters due to the presence of a hexamer motif which resembles the *TATA* box sequence, required for transcriptional initiation.^72^ However, the effect of an upstream terminator on gene expression has not been determined. To test whether an upstream terminator modulates the activity of the downstream expression cassette, we also tested the doxycycline-inducible promoter (*tetOpr*) with the *ADH1* terminator placed upstream of the promoter, which we refer to as *t-tetOpr*.

### Measurement process

We measured the induction dynamics by tracking single cells using time-lapse microscopy. Cells were grown in non-inducing medium overnight (>12 h), diluted to remain in log phase. Then, the *promoter-yEVenus-PEST* circuit was induced for 3.5 hrs, then shut off, and monitored for another 3 hrs. The period of induction corresponded to roughly 2.5 budding yeast cell cycles in glucose medium, a sufficiently long time for many applications. A summary of the inducing and non-inducing/repressing conditions is given in Table 1. Detailed descriptions of the conditions are given in Supplementary Note 1. For tuning the induction level of the blue light-inducible systems, it was convenient to use the diascopic LED source.

**Table 1.** Inducing and non-inducing/repressing conditions used for controlling the activity of inducible systems.

| Transcriptional system | Inducing condition | Non-inducing or repressing condition |
| --- | --- | --- |
| *GAL1pr*, *GALL* | Galactose | Raffinose, Glucose |
| *MET3pr* | Absence of methionine | Methionine |
| *CUP1pr* | Cu<sup>2+</sup> ions | Absence of Cu<sup>2+</sup> |
| *PHO5pr* | Absence of inorganic phosphate | Inorganic phosphate |
| *tetOpr*, *t-tetOpr* | Doxycycline | Absence of doxycycline |
| Z3EV | Estradiol | Absence of estradiol |
| El222-*LIP*, strongLOV-*LIP*,<br>El222-*GLIP* | Blue light | Absence of blue light |
| PhyB-PIF3 | Red light ($\approx 650$ nm) and PCB | Far-red light ($\approx 750$ nm) |

To quantify the inducible systems’ characteristics, intuitive and transferable units are needed. Given that *GAL1pr* is plausibly the most widely used inducible system in yeast, the strongest one among the ones tested by us, and has been adapted for other model systems such as *Drosophila sp.*^73^ and mammalian cell lines^74^, we introduce a unit for promoter activity which we denote maxGAL1. The value of 1 maxGAL1 corresponds to the stationary level of expression from a single *GAL1* promoter (Fig. 5 D). Introducing a unit allows easy comparison of promoter strengths from different sources assuming that the promoter construction is standardized within each set of experiments and that one includes *GAL1pr* as a reference. For example, the activity of frequently used constitutive promoters can be expressed in terms of maxGAL1, with *PGK1pr* and *TEF1pr* in glucose having ≈0.40 maxGAL1 and *TDH3pr* ≈0.70 maxGAL1 transcriptional activity^75^.

**Figure 3.**
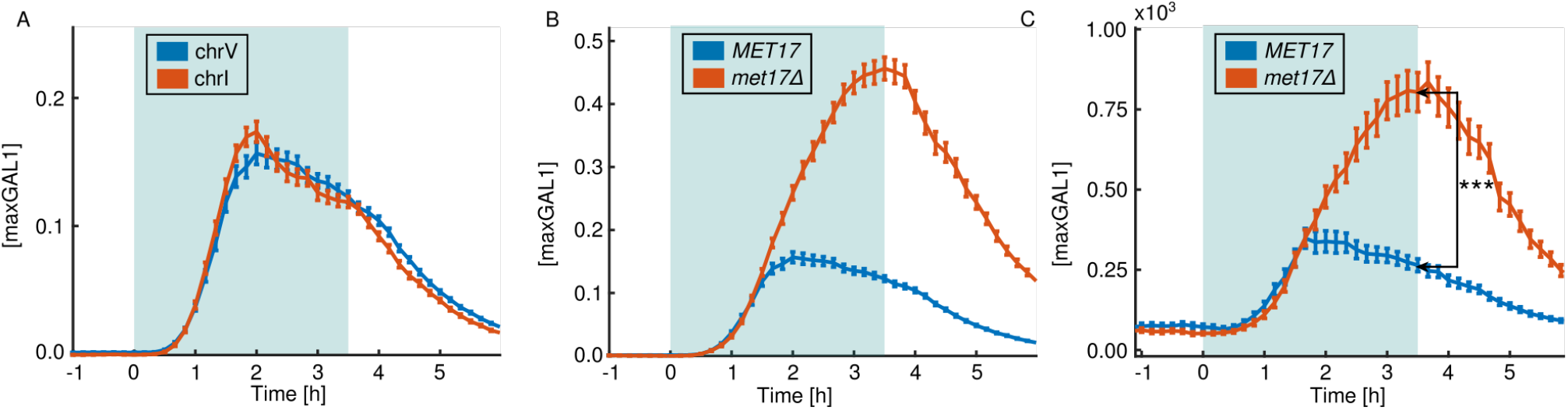
Feedback-mediated cellular production of methionine contributes to an overshoot and decline of *MET3pr* activity. A: Changing the integration site does not alleviate the overshoot as the amplitude and timing remain unaffected. Induction of two constructs is shown, one integrated at the *URA3* locus on chromosome V and one integrated on chromosome I between the *SSA1* and *EFB1* open reading frames. B: Induction of the *MET3pr-yEVenus-PEST* construct in the *met17Δ* background and wild type. C: Total levels of yEVenus fluorescence per cell (instead of the average fluorescence as in panels A and B) suggest that dilution of the fluorescent protein due to growth does not explain the observed differences in *MET3pr* activity. p = 3.7e-11, one-tailed t-test. A, B, C: The blue background represents the induction period, i.e., lack of methionine. Bars around the points show the standard error of the mean (SEM). Numbers of analyzed cells are given in Supplementary Table 15.

**Figure 4.**
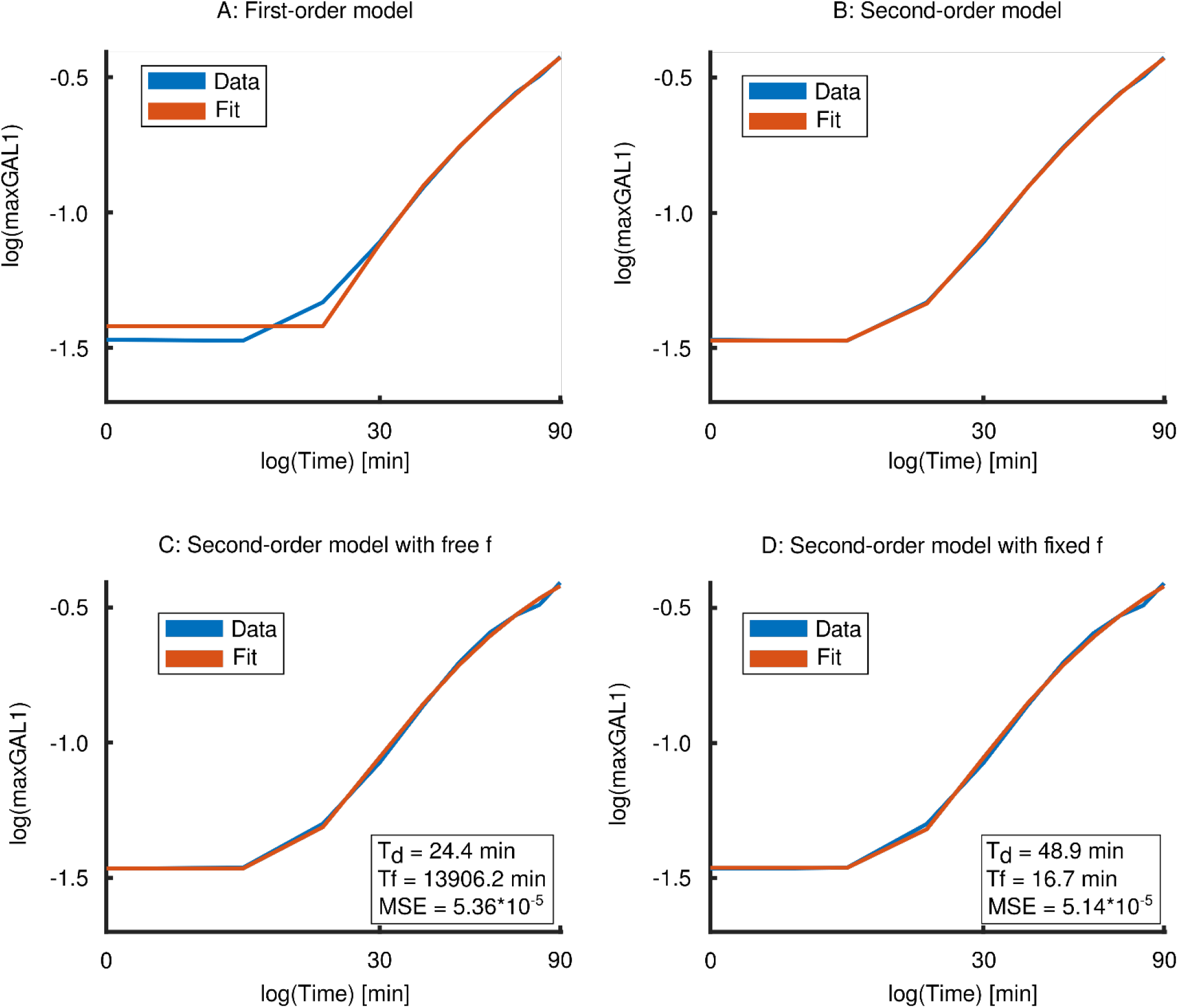
A second-order differential equation fits the time courses well. A: A first-order differential equation model (red) does not fit the time course of El222-driven *LIP-yEVenus-PEST* expression with 80% light induction (blue) well. B: The expression dynamics is fit well by a second-order differential equation model. C, D: However, the second-order differential equation model is not constrained sufficiently since the data can be fit well using very different parameters for *d* and *f*. To avoid this, we will fix the maturation rate *f* in the second-order model by measuring it in an independent experiment (see Supplementary Note 2). Fits are shown for single-cell data expression of *LIP-yEVenus-PEST*. MSE: mean squared error of the fit. T_d_ and T_f_ are ln(2)/*d*, and ln(2)/*f*, respectively.

**Figure 5.**
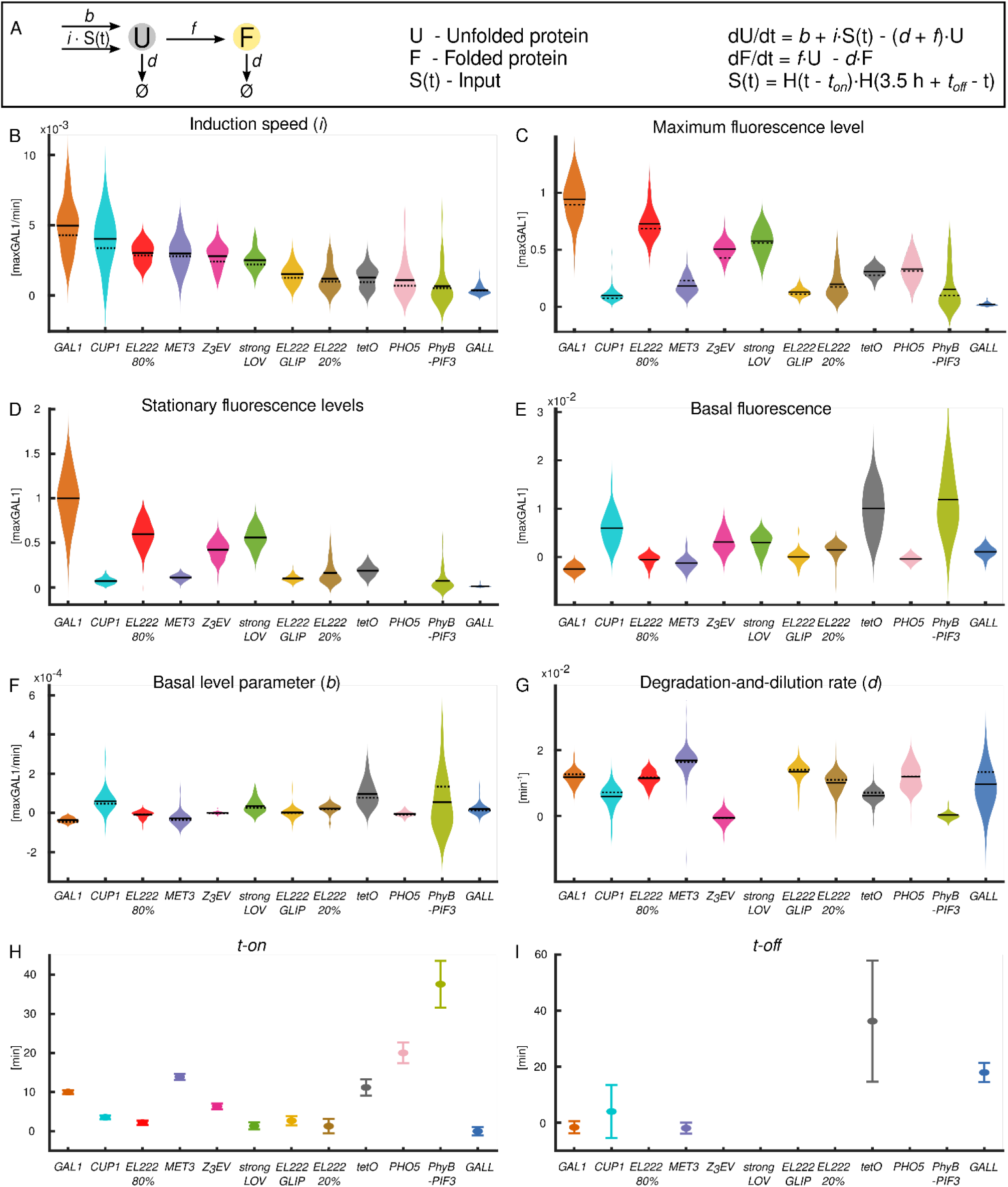
Single-cell-level characteristics of the inducible transcriptional systems. Violin plots show distributions of parameters estimated by fitting fluorescence levels from single cells. Black solid lines show the mean of the distribution. Black dashed lines in panels B-G represent the extracted parameters after averaging over all cells first. *EL222 20%, EL222 80%, strongLOV,* and *GLIP* are defined in the caption of Fig. 2. A: Model of gene expression used to extract the quantitative parameters describing the inducible systems. H(t) is the Heaviside step function, 0 for t < 0 and 1 for t >= 0. B: Speed of induction *i*. C: Maximum fluorescence levels. D: Steady-state level of induction was defined for *CUP1pr*, *MET3pr*, Z_3_EV, El222 with 20% induction light*, GLIP*, PhyB-PIF3, strongLOV and *GALL* as the level of induction at t = 3.5 h. For *GAL1pr*, El222 with 80% induction light, and *tetOpr*, we defined the steady-state levels after overnight (>16 h) induction since these systems did not reach steady-state levels during the first 3.5 hours of induction. The steady-state level of *PHO5pr* is not shown given that the prolonged lack of inorganic phosphate causes cell-cycle arrest. E: Basal fluorescence levels. F: Basal activity parameter *b*. G: Degradation-and-dilution rate *d*. Degradation-and-dilution rate not shown for strongLOV-*LIP* because it could not be measured by fluorescence microscopy and was determined by sampling cells from liquid culture (Fig. 13 B). H: Time delay upon activation *t-on*. I: Time delay upon deactivation *t-off*. H, I: To estimate the time delay upon deactivation reliably, we fitted the model only to the average expression values, not single-cell data. Standard errors of the mean shown in panels H and I were estimated by bootstrapping single-cell expression values and fitting 100 averaged time courses to the model. H, I: For all systems except the light-inducible ones, we removed the time it takes the medium to reach the microfluidics chamber (160 s) from the estimated *t-on* and *t-off* values. For light-inducible systems and *PHO5pr*, *t-off* could not be determined precisely (see main text). “pr” in the promoter names was omitted for brevity. For determining the PhyB-PIF3 leakiness and parameter b, we noticed that a few cells (n = 3) out of 33 showed substantially higher values than the rest of the population. Since this causes an increase of the bandwidth for the violin plot and prevents clear visualization of the other systems’ leakiness, we excluded these cells from the main figure but provide the panels E and F without the removal of the cells in Supplementary Fig. 6. The p-values for differences between different systems are shown in Supplementary Tables 3-10. Numbers of analyzed cells are given in Supplementary Table 14.

### Single-cell time courses

The strength of the systems varied more than 50-fold, from ≈0.02 maxGAL1 for *GALL* to ≈1 maxGAL1 for *GAL1pr* (Fig. 2). (The standard deviation (SD) reflecting noise is discussed in Section ‘Noise’. The data is replotted with the standard error of the mean (SEM) in Supplementary Fig. 2 showing that the underlying cell numbers sufficed for determining the mean.) Interestingly, several systems showed complex dynamics upon induction. *MET3pr* and *GALL* exhibited a decline in activity for t > 1.5 h. The initially weak activation of *PHO5pr* was followed by substantially stronger induction starting at around t = 2 h. In addition, *CUP1pr* and *GALL* showed strong temporal fluctuations (single-cell trajectories in Fig. 2 L, N). The *tetO* promoters showed a substantial delay in shut-off compared to other systems. We found that a terminator placed upstream of the *tetO* expression cassette did not have a substantial effect on the expression dynamics. Given that the expression pattern of *tetOpr* was hardly distinguishable from the one of *t-tetOpr*, we focused on characterizing *tetOpr* only in the subsequent analyses. The red-light inducible optogenetic system showed high stochasticity, with only 25% of cells being substantially activated by the red-light pulse (t = 3.5 h). The comparison between the Gal4-based transcriptional PhyB-PIF3 system and the PhyB-PIF3 system used for subcellular localization suggests that the high stochasticity comes from the DNA binding functionality (Supplementary Fig. 3). Expression of *yEVenus* in cells with the Z_3_EV system stayed high even 3 hrs after estradiol was depleted from the medium. We wondered whether this sustained activity could be due to the hormone sticking to the surfaces of our microfluidic chips, continuously activating the system. To test this, we monitored the transcriptional activity after a thorough washout in liquid culture. The results showed that the system needs several hours to begin to turn off (Supplementary Fig. 4) independently of any potential adhesion of estradiol to the microfluidic chamber walls.

Intriguingly, *MET3pr* showed an overshoot and partial adaptation after induction. To investigate the mechanism for this, we changed the site of integration of the construct, which had no apparent influence on *MET3pr* expression dynamics (Fig. 3 A). To test whether the partial adaptation could be attributed to upregulation of methionine biosynthesis upon removal of methionine from the medium, we deleted the *MET17* gene (also known as *MET15, MET25*), which is responsible for most of the synthesis of homocysteine, the precursor of methionine.^76,77^ Inducing the single-copy *MET3pr-yEVenus-PEST* construct in the *met17Δ* background generated a stronger response to methionine depletion and without the distinctive overshoot (Fig. 3 B). Since methionine depletion in *met17Δ* cells causes growth defects that might kick in during the 3.5 hrs of the *MET3pr* induction, we tested whether the lack of overshoot in the *met17Δ* mutant can be simply explained by a lower protein dilution rate. To exclude the effect of cell growth, we compared the *total* yEVenus levels accumulated during the 3.5 hrs. Even after accounting for the differences in growth, the final level of yEVenus was higher in the *met17Δ* than in the WT background (Fig. 3 C), suggesting that feedback from methionine biosynthesis contributes to the partial adaptation of *MET3pr* activity.

The *PHO5* promoter presented another intriguing time course. The observed two-step induction pattern could be due to phosphate depletion beginning to block growth at about 2-2.5 hrs after induction, thus, preventing dilution of yEVenus. The activation pattern could also be due to fluctuations in cytosolic phosphate levels during induction; for example, phosphate released from the vacuole could be depleted at 2-2.5 hrs. We decided to test whether a growth block makes the fluorescence from the *PHO5pr-yEVenus-PEST* reporter, averaged over the cell area, appear to shoot up. Thus, we analyzed the growth rate of cells during the last hour of induction. Cells showed a healthy growth rate comparable to cells grown in synthetic complete media (Fig. 7). Thus, changes in growth rate are not responsible for the second jump in *PHO5pr* activity. On the other hand, the reported timing of polyphosphate exhaustion from the vacuole^52^ matched the time of the second jump in *PHO5pr* activity. Thus, internal phosphate stores are more likely to be responsible for the two-step transcriptional *PHO5pr* activation pattern than effects on growth and dilution.

**Figure 6.**
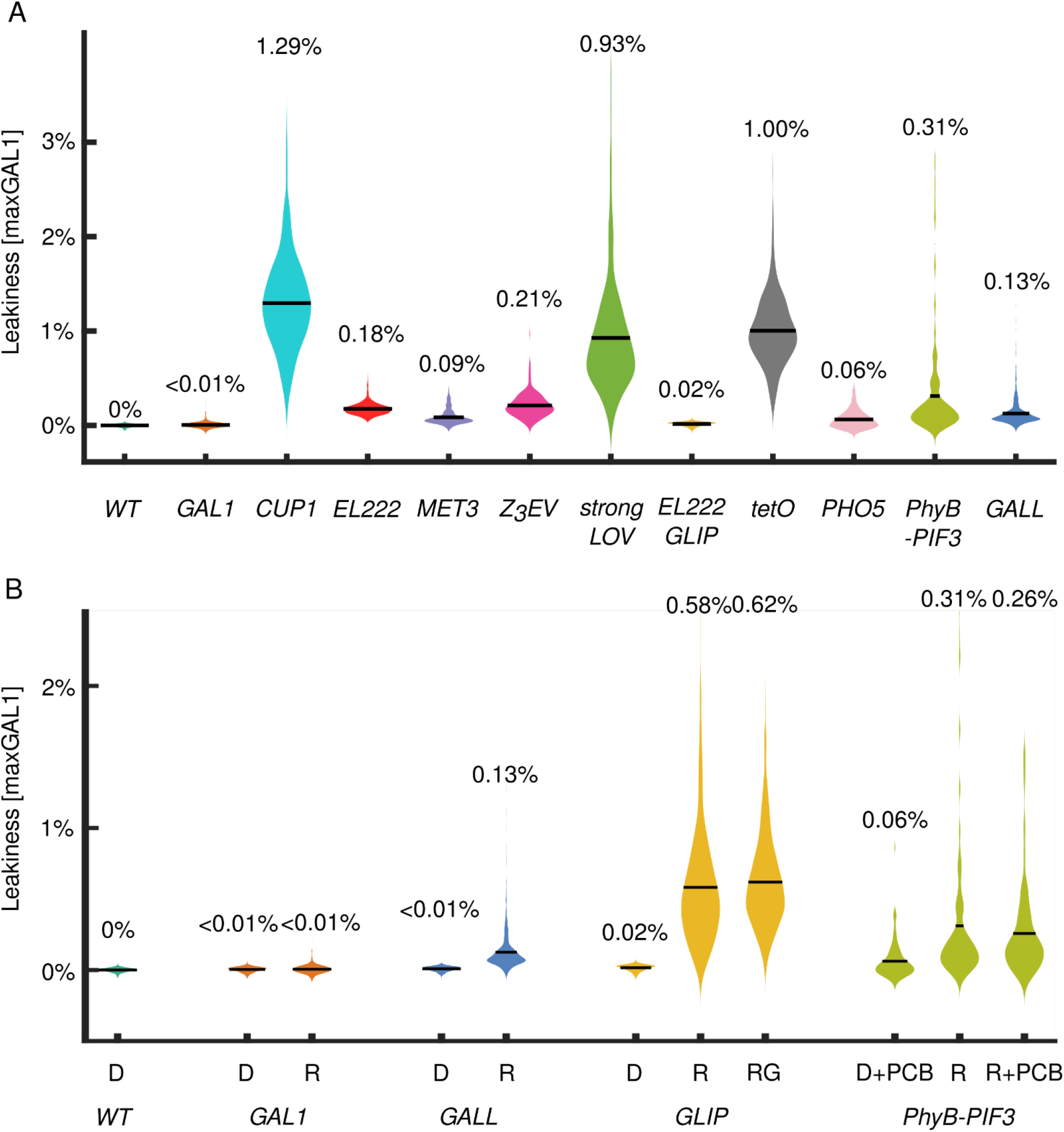
Minimal leakiness measurements using *promoter-yEVenus* reporters (without the PEST degron). A: Removal of the *PEST* sequence from the transcriptional reporters uncovers the leakiness of each system. *GALL, GAL1pr,* and PhyB-PIF3 leakiness was measured in raffinose. *EL222* refers to leakiness of the El222*-LIP* system. *strongLOV* refers to the leakiness of strongLOV*-LIP*. B: Basal activities of *GAL1pr, GALL, GLIP,* and PhyB-PIF3 depend on the carbon source, D - glucose, R - raffinose, G - galactose. A, B: “pr” in the promoter names was omitted for brevity. The measurements were calibrated with respect to the previous figures (where the PEST degron was present) using the leakiness of *tetOpr*. Thus, all expression levels are comparable across different figures and are always normalized to peak *GAL1pr* expression levels, i.e., shown in maxGAL1 units. Average values for each measurement are shown above the corresponding violin plots. p-values for statistical significance of differences are given in Supplementary Tables 11-12. Numbers of analyzed cells are given in Supplementary Table 16.

**Figure 7.**
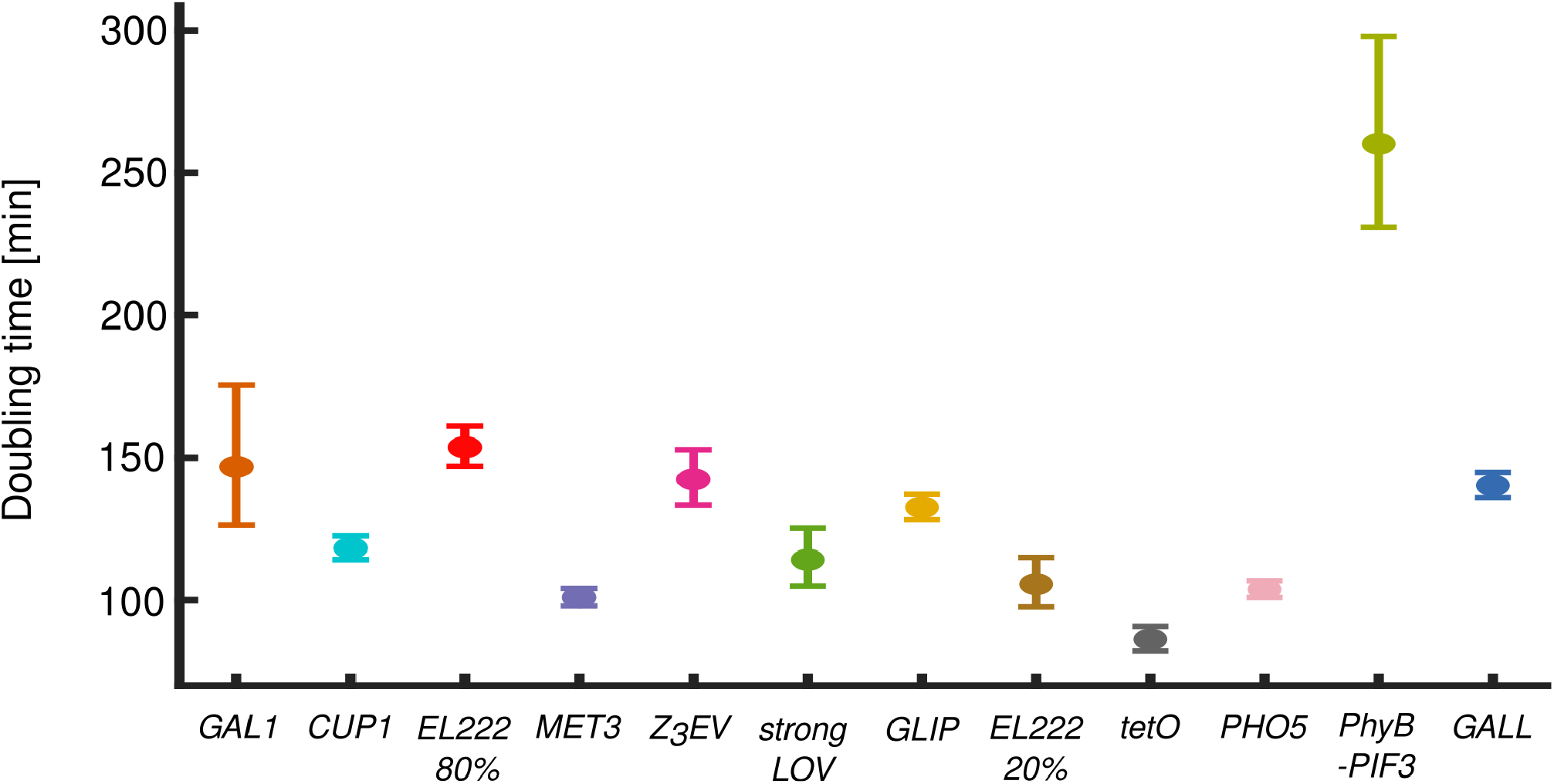
Area doubling time of cells harboring different inducible constructs in the induced state. Error bars represent 90% confidence intervals. *EL222 20%, EL222 80%, strongLOV,* and *GLIP* are defined in the caption of Fig. 2. “pr” in the promoter names was omitted for brevity. p-values for the differences between the pairs of parameters are supplied in Supplementary Table 13. Numbers of analyzed cells are given in Supplementary Table 14. For *GAL1pr*, the colony that was not fully present in the field of view and which would bias the estimation of the growth rate, was excluded, reducing the number of analyzed cells shown in Supplementary Table 14 to 79 (t = 3.5 h).

### Mathematical model of inducible transcriptional system dynamics

We wished to distill the time courses for each inducible system (Fig. 2 B-N) into intuitive parameters. Quantitative descriptions could not simply be extracted from the time courses ‘by eye’. This is because the time courses did not, for example, consist of piece-wise linear functions, which allow one to read off parameters directly. Instead, the time courses were smooth (see Fig. 4 A for a magnified plot of the initial rise of the fluorescence). There are two well-known reasons for this smoothing, the maturation and the degradation-and-dilution times of the destabilized fluorescent protein reporter, with previously reported timescales of ≈20 min and ≈40 min, respectively^43^. A sudden increase in fluorescent protein expression manifests as a smooth increase with these two timescales determining how fast the fluorescence follows the underlying transcriptional dynamics. Thus, to extract parameters from the time courses, a mathematical model needed to be fit.

A minimal model would have parameters with obvious meanings and would prevent overfitting. To identify the minimal model complexity that was needed, we analyzed the initial rise in fluorescence (Fig. 4). This part of the time course fit a quadratic function well (slope of 1.85, 99% confidence interval: 1.67-2.03, on a log-log scale for the time points from t = 20 min to t = 70 min). Thus, a second order differential equation, in which the activation of the promoter is a step function (Fig. 5 A), was called for. Such a model has been used previously^29,32,43,67^. The first equation in this model describes the expression dynamics of the unfolded fluorescent protein. The maturation of the fluorescent protein, a slow step during gene expression, is modeled by the second equation. Both steps are affected by protein degradation- and-dilution equally. In the model, the basal (non-induced) expression is controlled by *b*. Promoter activity upon inducer addition is determined by an initial lag *t-on* between the start of the induction signal and the start of gene expression. The initial slope of the unfolded protein rise is denoted by *i*. The time between the inducer removal and the start of decline in promoter activity is characterized by the lag *t-off*. The rate of the fluorescence decay after promoter turn-off is characterized by degradation-and-dilution rate *d*.

Approximating the initial rise of fluorescence using a simpler, first-order model yielded poor fits (Fig. 4 A). Therefore, using simple methods such as tresholding also fails to extract parameters accurately. On the other hand, increasing the order of the model requires more parameters to be extracted from the data. Already with the second-order model, we observed that the data does not constrain the parameters enough since, for example, very different kinetics of promoter activation *f* and *d* fit the data equally well (Fig. 4 C and D). To prevent this, we had to measure the yEVenus maturation rate in our experiments directly and used this value as a fixed parameter *f* when fitting the model to the data (see Supplementary Note 2). Thus, with the model we chose, all remaining unknown parameters (*b, i, d* and the time delays) could be uniquely identified based on the fluorescent protein level measurements only (see Methods section for mode details). No parameter could be removed without the fit clearly becoming worse, and adding more parameters led to poorly constrained parameters and overfitting. Extending the model to characterize the gene expression process in greater detail would be possible by using more experimentally measured parameters^18,29^ but was not necessary nor desirable for the purpose of extracting intuitive quantitative characteristics and benchmarking.

Note that *GALL*, *MET3pr*, *CUP1pr*, and *PHO5pr* show more complicated time courses. To be able to compare the different systems using quantitative parameters nevertheless, we used the model only for the rise (from -50 min to 50 min for extracting *b*, *i* and *t-on*) and the fall (from 210 min to 270 min for *t-off* and 270 min to 390 min for *d*) of the time courses (see Methods sections for more details on fitting procedure). While interesting and potentially important for certain applications, the rest of the dynamics is not comparable between all of the different inducible systems. Thus, we only distill the dynamics around the on and off switches into coarse-grained parameters.

### Inference of intuitive parameters

By fitting the model in Fig. 5 A to the observed fluorescence values (Methods), we extracted the values for the initial speed (*i*), basal activity (*b*), degradation rate (*d*), and lag upon activation and deactivation (*t-on* and *t-off*, respectively). In cases where the systems did not reach their maximal activity during the 3.5 h induction period, we measured the steady-state expression levels after an overnight growth in inducing media with dilutions to keep cells in log phase throughout. Single-cells fits are shown in Supplementary Fig. 7.

#### Initial speed (i)

The initial speed *i* spanned a 10-fold range, with *GAL1pr* being the fastest and *GALL* the slowest system (Fig. 5 B).

The initial slopes of induction were as follows:

*GAL1pr > CUP1pr >* El222-*LIP, 80%> MET3pr >* Z_3_EV *>* strongLOV *>* El222*-GLIP >* El222-*LIP, 20% > tetOpr > PHO5pr >* PhyB-PIF3 *> GALL*

#### Maximum level and steady-state ‘on’ levels

It is interesting that the maximum induction levels (Fig. 5 C) did not necessarily reflect the initial speed of the induction. Because some inducible transcriptional systems showed transient dynamics, e.g., an overshoot, which was not followed by a long-term, steady-state behavior of the system, we also measured the steady-state induction levels (Fig. 5 D). For systems that reach a stationary expression level during the 3.5 h long induction experiment (Fig. 2), the steady-state level was defined as the level at the last timepoint of induction (Fig. 5 D). Given that *GAL1pr*, *LIP*, and *tetOpr* did not reach steady-state levels during the 3.5 h induction period, we measured the expression levels for these systems after an overnight (> 16 h long) induction (Fig. 5 D) during which the cultures were diluted to keep them in log phase. *PHO5pr* did not reach a steady-state level after 3.5 h but given that the prolonged absence of inorganic phosphate causes cell cycle arrest^78^, we did not perform an overnight induction for this system. Hence, steady-state levels and *t-off* for *PHO5pr* are not shown.

Steady-state induction levels were as follows:

*GAL1pr >* El222-*LIP, 80% >* strongLOV *>* Z_3_EV *>* El222-*LIP, 20% > tetOpr > MET3pr >* El222*-GLIP >* PhyB-PIF3 *> CUP1pr > GALL*

#### Basal activity (b) / leakiness

With three exceptions, the *promoter-yEVenus-PEST* reporters showed no activity in the off state, that is, no leakiness at the level of sensitivity of fluorescence microscopy (Fig. 5 E). Only the *CUP1pr*, *tetOpr*, and the PhyB-PIF3 system showed considerable levels of expression (approx. 1% maxGAL1) in the absence of the inducing signal. Therefore, we boosted the sensitivity of our system by removing the *PEST* sequence, the results of which are presented in the Section ‘Leakiness’ (Fig. 6). Note that in Fig. 5 E, negative values arise in part because auto-fluorescence can vary across a population and is systematically lower in raffinose versus glucose media.

#### Degradation-and-dilution rate rate (d)

The rate *d* includes two components: active degradation of the reporter protein which is destabilized by the PEST degron and degradation through dilution due to cellular growth, which is non-negligible in fast-growing cells such as budding yeast. Interestingly, we measured large differences in degradation-and-dilution rates for the different systems. We hypothesized that this is due to differences in growth rates under different inducing conditions (see Section ‘Effect of induction conditions on cellular growth’, Fig. 7, 8 for more details). Indeed, for most of the inducible systems, the overall degradation-and-dilution rate changed linearly as a function of the growth rate with slope equal to one. *GALL* showed substantial variations in degradation-and-dilution rates due to the large temporal fluctuations during the induction period introducing large variability in the estimated parameters (single-cell trajectories of cells shown in Fig. 2). For the four inducible systems Z_3_EV, PhyB-PIF3, *CUP1pr*, and *tetOpr*, the very small degradation- and-dilution rates could not be explained by slow growth alone since they fell far from the linear regression line, indicating particularly slow turn-off of these systems after the induction signal was turned off. (We discussed the slow turn-off for Z_3_EV above.)

**Figure 8.**
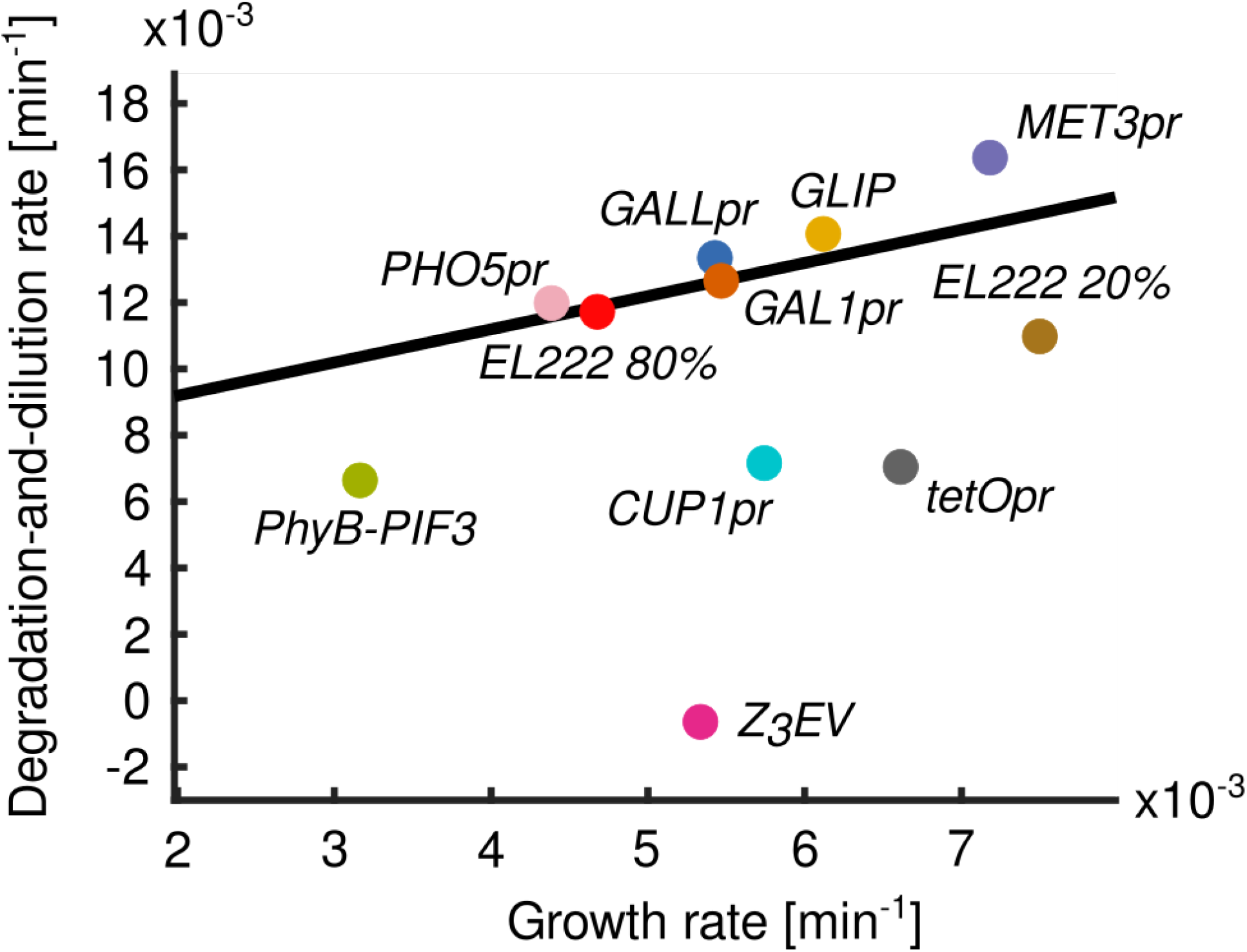
The differences in fitted degradation-and-dilution rates *d* in the different systems can be largely explained by differences in growth rates. Since the overall degradation-and-dilution rate *d* is the sum of the rate of dilution due to growth and the rate of degradation by the protein degradation machinery, we performed a linear fit with slope fixed to one. The fit shown in black is obtained by excluding *CUP1pr*, *tetOpr*, PhyB-PIF3, and Z_3_EV, which deviate from the general trend. The most prominent outlier is Z_3_EV. Cells with the *Z_3_EV* system continue to grow but do not turn the construct off, resulting in a degradation-and-dilution rate close to zero. Similarly, for PhyB-PIF3, *CUP1pr*, and *tetOpr*, the overall degradation-and-dilution rate *d* is smaller than expected given the growth rate. This can be due to residual transcription in the absence of the inducer. Incidentally, among the inducible systems in the plot, these are also the systems for which the fundamental leakiness was the highest. *EL222 20%, EL222 80%, strongLOV,* and *GLIP* are defined in the caption of Fig. 2. Degradation-and-dilution rates shown here are extracted from averaged fluorescence values, and are the same as the ones shown in Figure 5. Growth rates shown in the plot are calculated using the same timepoints as for the degradation-and-dilution rate (for exact values see Materials and methods section). Note that these growth rates can be different from the ones measured during the last hour of the induction period, which are shown in Fig. 7. The only exception from this is *PHO5pr*, for which we neglected the timepoints after which cells abruptly stopped growing presumably due to a lack of inorganic phosphate in the induction medium.

#### Lag times (t-on, t-off)

The lag times turned out to be particularly sensitive to temporal fluctuations in the single-cell time courses. Therefore, we extracted the delay upon activation (*t-on*) and upon deactivation (*t-off*) of the inducible systems after averaging the time courses over the population, resulting in smoother time courses (Fig. 2). To estimate *t-off* precisely for the systems that did not reach steady-state levels during the 3.5 h induction period (*GAL1pr*, *tetOpr*), we performed an experiment in which we switched off the system after an overnight induction, keeping cells in log phase throughout. *In vitro* measurements of *t-off* for the El222 protein have shown that this parameter is on the order of 1 min.^79^ Because this quantity has already been measured accurately and because the fluorescence measurements themselves induced *LIP* and *GLIP*, we do not report *t-off* for these two systems. In contrast, we could measure *t-on* accurately for *LIP* and *GLIP* by just not taking any images before induction.

Time delays upon activation and deactivation of the constructs (Fig. 5 H and I) are summarized below:

*t-on*: *GALL<* El222-*LIP, 20% <* strongLOV *<* El222-*LIP, 80% <* El222*-GLIP < CUP1pr <* Z_3_EV *< tetOpr < GAL1pr < MET3pr < PHO5pr <* PhyB-PIF3

*t-off*: *MET3pr < GAL1pr < CUP1pr < GALL< tetOpr*

### Leakiness

For many applications, e.g., expression of toxic genes, the basal activity of the inducible systems is critical and needs to be known. Yet, it has not been measured systematically or quantitatively. To determine leakiness rigorously, we boosted the reporter levels by removing the *PEST* sequence, and measured activities in non-inducing conditions. While the degron was important for quantifying the expression dynamics, it was not needed to measure leakiness, which is a steady-state property. Crucially, since the strains only had one copy of the *promoter-yEVenus* constructs, we were able to measure the minimal, fundamental leakiness of each system. For the synthetic inducible systems, the transcription factor levels allow further tuning of the strength and leakiness of the systems; however, our measurements showed stark differences between the different systems, making this an inauspicious avenue for substantially changing the ranking of the different systems with respect to this characteristic.

In glucose, all systems except *GAL1pr*, *GALL*, and *GLIP*, showed leakiness greater than 0.05% maxGAL1 (Fig. 6 A). As expected from the previous measurements with the PEST degron (Fig. 5 E), the *tetOpr*, *CUP1pr*, strongLOV driving *LIP*, and PhyB-PIF3 systems showed the highest levels of leakiness.

The tight nature of the *GAL1* and *GAL1*-based promoters might come from a glucose-repression system that is independent of the Gal4/Gal80 activator/repressor system^80^ and is mediated by the Mig1 repressor. To investigate this, we measured the basal activity of *GAL1pr*, *GALL*, and *GLIP* in media with different sugars (Fig. 6 B). *GAL1* showed no detectable leakiness in glucose or raffinose, in which the Gal4 activator is repressed by Gal80^81^. However, *GALL* showed detectable basal expression in raffinose. For complete repression of *GAL* genes by Gal80, two adjacent Gal4-binding sites are needed as in *GAL1pr*.^82^ In contrast, *GALL* contains only one of the two sites from *GAL1pr*, which may explain its increased level of basal activity in raffinose compared to glucose (for visual representation see Supplementary Fig. 7 B). Similarly, *GLIP* showed significantly higher basal levels of expression in raffinose and galactose, compared to glucose. Given that *GLIP* inherited the Mig1 binding sites from *GAL1*, this difference is presumably due to basal activity of the El222 transcription factor that becomes detectable once the inhibition by the glucose-repression system is alleviated (Supplementary Fig. 7 C). However, although the endogenous *GAL80* repression machinery was present, the Gal4-based PhyB-PIF3 system caused substantial leakiness of *GAL1pr* in raffinose (Fig. 6 B), presumably because the split Gal4 protein in this system is no longer sufficiently repressed by Gal80^81^.

The doxycycline-inducible system, used widely in many different organisms, showed remarkably high levels of basal expression (≈1% maxGAL1), comparable to the induced state of *GALL*. To address the leakiness problem, mutant doxycycline-responsible transcription factors were developed in ref. ^83^. Testing the tightest of those systems, the rtTA system, we observed under a variety of doxycycline concentrations and induction times that the induction was highly unreliable and generated substantial cell-to-cell variability (Supplementary Figure 9). Thus, the leakiness of the *tetOpr* system remains an important concern for applications.

The basal activities of the systems shown in Fig. 6 A are summarized below:

*CUP1pr >* strongLOV *> tetOpr >* PhyB-PIF3 *>* Z_3_EV > El222-*LIP > GALL > MET3pr > PHO5pr >* El222- *GLIP > GAL1pr*

### Effect of induction conditions on cellular growth

Expression systems may interfere with growth due to less favorable nutrient conditions needed for induction, toxicity of the inducers, or metabolic burden^18^. To benchmark the systems with respect to cell growth, we measured the doubling times of the areas of the cell colonies during the last hour of induction (2.5 h < t < 3.5 h) (Fig. 7).

The diascopic light used to induce the expression of LIP had an effect on growth when applied at 80% of the maximal strength (Fig. 7 *EL222* 80% and *GLIP*). Cells exposed to light at 20% of maximal strength had a more healthy area doubling time of around 100 min (Fig. 7 *EL222* 20% and strongLOV).

Cell size doubling times during the last hour of induction are summarized below:

*tetOpr < MET3pr < PHO5pr <* El222-*LIP*, 20% *<* strongLOV *< CUP1pr <* El222*-GLIP < GALL <* Z_3_EV *< GAL1pr <* El222*-LIP*, 80% *<* PhyB-PIF3

Since growth dilutes cellular contents, we wished to analyze how active degradation due to the PEST degron and dilution due to cell growth contribute to the overall degradation-and-dilution rate *d* from the model. By plotting *d* versus the growth rate, we found that the relationship was explained well by a line with slope 1 with a few prominent exceptions (Fig. 8). This indicates that the differences in degradation-and-dilution rates are mostly due to differences in the growth rates. The intercept of the optimal fit is 0.0072 min^-1^, from which the half-life of yEVenus-PEST can be calculated: ln(2)/0.0072 min = 96.3 min. This agrees with the yEVenus-PEST degradation half-life which we also measured directly by blocking protein translation with cycloheximide (Supplementary Note 3).

Furthermore, Z_3_EV, PhyB-PIF3*, CUP1pr,* and *tetOpr* fell far below the linear regression line (Fig. 8), indicating that the slow degradation-and-dilution rates cannot be explained by slower growth. Instead, these systems are turning off slowly.

### Noise

Within a population of genetically identical cells, the responsiveness of a genetic circuit can vary. The relationship between mean and standard deviation can be complex.^84,85^ To investigate this for inducible transcriptional systems, we calculated the coefficient of variation for the last timepoint (t = 3.5 hrs) of induction in the time course experiments (Fig. 9).

**Figure 9.**
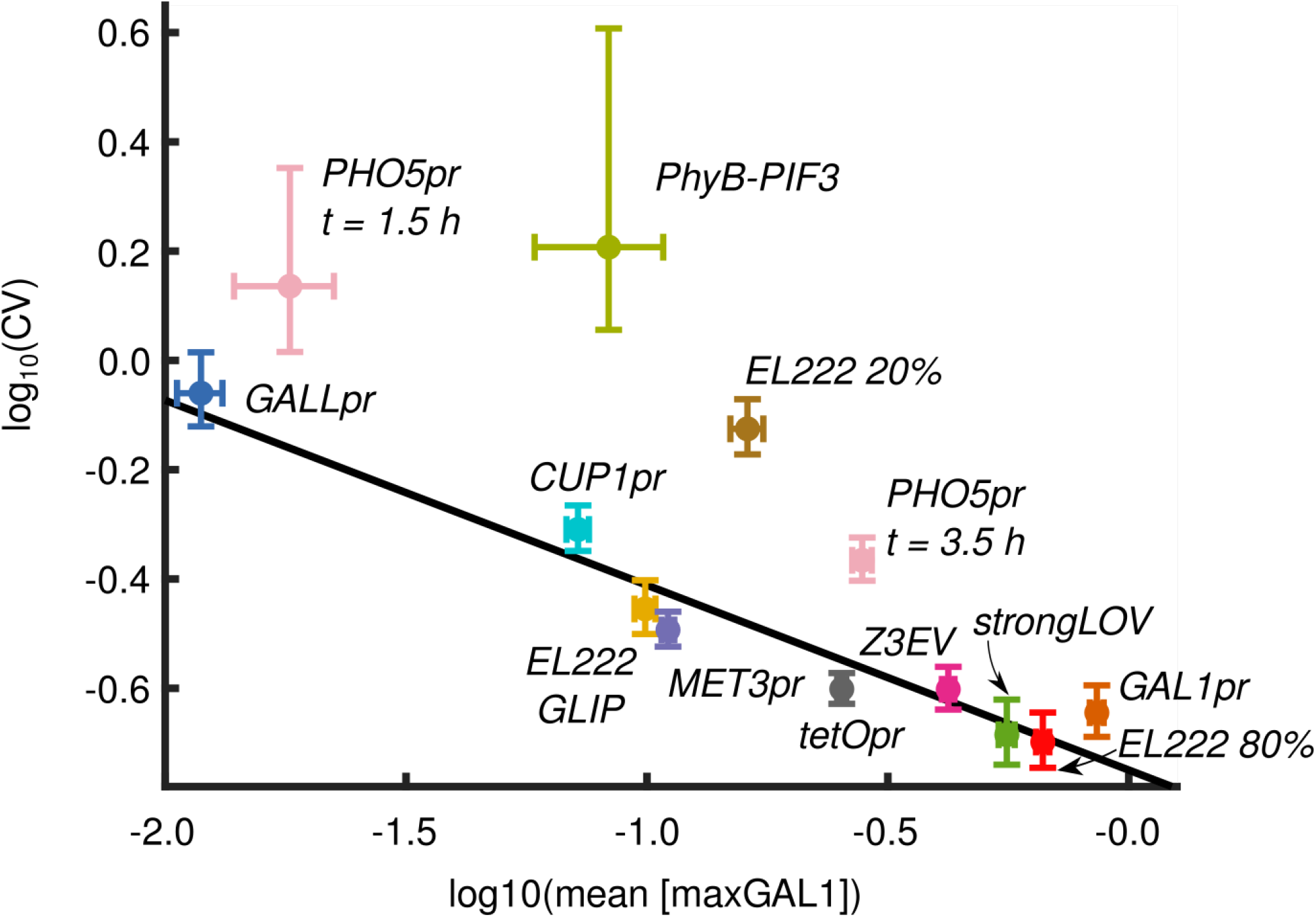
Log of noise (CV) versus mean expression levels are inversely correlated. Noise is calculated as the coefficient of variation for the population of cells at the last timepoint of induction, t = 3.5 h, unless stated otherwise. Fluorescence values are in maxGAL1 units. Vertical and horizontal bars around the values show 90% confidence intervals. *EL222 20%, EL222 80%, strongLOV,* and *GLIP* are defined in the caption of Fig. 2. The least squares regression was computed after excluding PhyB-PIF3, El222 induced with 20% light intensity, and *PHO5pr*, slope = -0.34, R^2^ = 0.93, 95% confidence interval: [-0.42, -0.26]. Numbers of analyzed cells are given in the Supplementary Table 14.

As expected^86,87^, noise levels decreased with the increase in the mean expression level, meaning that the strongest inducible systems were also the least noisy ones. The coefficient of variation scaled linearly with the mean level of expression on a log-log scale (Fig. 9). PhyB-PIF3 showed a high level of noise for its mean expression level compared to other systems. To test whether this observation can be associated with noise in the PCB internalization by cells, we constructed a plot similar to Fig. 9 but under non-induced conditions and in the absence of the PEST degron (Supplementary Figure 8). We observed a similar level of leakiness noise in the PhyB-PIF3 system with and without PCB, indicating that high noise in this system is not due to PCB internalization. Interestingly, the El222-*LIP* system induced under low light conditions (20% of maximal intensity) showed a comparatively high level of noise. *PHO5pr* was also noisy relative to its mean compared to other systems (Fig. 9). We wondered whether the additional slow step, in which the internal storage of inorganic phosphate has to be used up before *PHO5pr* is fully activated^52^, introduces additional noise. However, the level of noise for the *PHO5* promoter at t = 1.5 h after induction, before the second activation of *PHO5*, was also substantially higher than expected from the linear regression line (Fig. 9). Given the relatively low noise of the non-induced *PHO5* promoter (Supplementary Fig. 8), these results point to other mechanisms that might be contributing to the particular pattern of *PHO5pr* noise such as chromatin remodeling.^88,89^

### Characterization of the arginine-responsive promoter *ARG3pr*

We decided to expand our analysis by an additional promoter, *ARG3pr*, which is part of the arginine-synthesis pathway in budding yeast and has not previously been characterized for use as an inducible system. *ARG3* is essential for arginine biosynthesis, coding for ornithine carbamoyltransferase, which converts ornithine to citrulline, a precursor of arginine^90^. At the transcriptional level, *ARG3* is controlled by arginine availability through transcription factors Arg80, Arg81, and Arg82, which form the repressive ArgR complex^91^ as well as by general amino acid control mechanisms through the Gcn4 activator^92^.

We chose to characterize *ARG3pr* since the transcriptomic analysis of Gasch et al.^93^ showed that ARG3 is the 7^th^ most upregulated transcript upon amino-acid starvation longer than 30 min. For comparison, *MET3* is the 8^th^ most upregulated gene under the same conditions. The motivation to pursue *ARG3pr* came from our observation that many of the synthetic systems we benchmarked have important shortcomings and endogenous inducible systems such as *GAL1pr* and *MET3pr* are some of the overall best inducible promoters at least with respect to strength, speed, and reversibility. Furthermore, no additional transcription factors have to be introduced for endogenous systems, making them convenient in various situations where more cell or molecular biology work would be needed to introduce the synthetic transcription factor. For many applications, finding a third, good inducible trancriptional system in addition to the *GAL* promoters and *MET3pr* would be very useful.

As a first test, we transferred cells containing a single copy of an *ARG3pr-yEVenus-PEST* construct from synthetic complete medium to medium lacking all amino acids and measured the expression level of the reporter after 1.5 h (Fig. 10). Unexpectedly, *ARG3pr* activity decreased in response to amino acid depletion; supplying only the essential nutrients did not change this result (Fig. 10). Since *ARG3* is known to be also post-transcriptionally regulated^94^, we hypothesized that in synthetic minimal medium, the overall transcript levels might still increase if degradation of ARG3 mRNA decreased. To test this, we measured fluorescence levels in a strain with an *ARG3pr-ARG3-mNeonGreen* gene fusion^95^. Under the same starvation conditions, we observed an Arg3-mNeonGreen protein trend similar to *ARG3pr-yEVenus-PEST* (Supplementary Fig. 10). Thus, neither transcription from *ARG3pr* nor Arg3 protein levels reflect the strong upregulation of ARG3 mRNA reported by Gasch et al.^93^. The following results indicate that this could be due to minor but difficult-to-replicate or -control differences in media.

**Figure 10.**
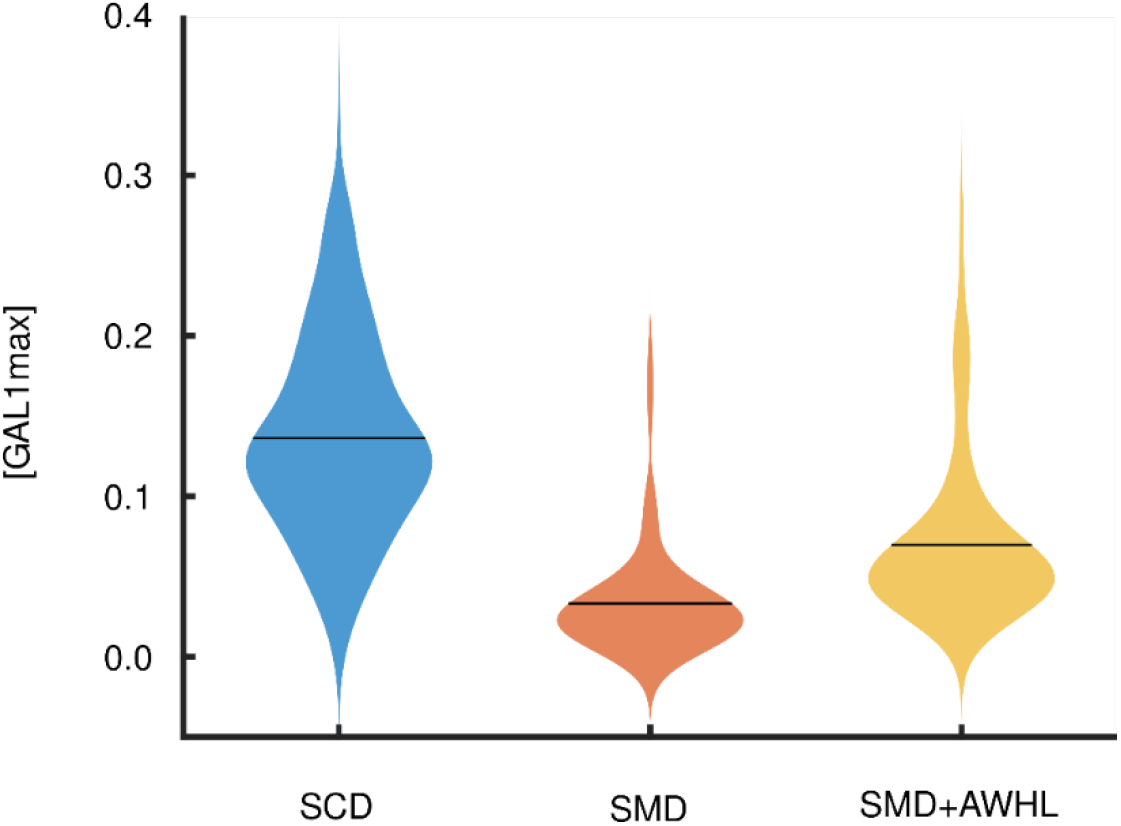
*ARG3* promoter is induced less in synthetic minimal medium than synthetic complete medium. SCM - Synthetic complete medium. SMM – Synthetic minimal medium. SMM+AWHL – Synthetic minimal medium with adenine, tryptophan, histidine, and leucine, for which our strain was auxotrophic. Horizontal bars denote the mean of the population. For details about media composition, see Supplementary Note 1. Numbers of analyzed cells are given in Supplementary Table 17.

Since using media without amino acids has the drawback that it slows down growth and blocks growth completely when cells are auxotrophic for the amino acids not present in the medium, we moved on to characterize *ARG3pr* when only certain amino acids were removed. Cells grown in synthetic complete medium did not show a substantial difference in *ARG3pr* induction in response to arginine removal (Fig. 11, the two violin plots on the right). We assumed that this behavior could be explained by the combinatorial regulation of *ARG3pr* with other nutrients present in synthetic complete medium which mask arginine regulation^96^. Thus, we analyzed the effect of arginine in combination with methionine, one of the nutrients that strongly upregulates *ARG3*^97^ and that would be used in combination with the *MET3pr* system. We found that methionine indeed activates *ARG3pr*. Interestingly, *ARG3pr* is turned on to a similar extent by either the absence of arginine, the presence of methionine, or both, resembling an OR logic function (-A OR +M) (Fig. 11 A). However, the basal level of activity in the presence of arginine and absence of methionine was relatively high, favoring the use of *ARG3pr* as a sensor in bulk culture.

**Figure 11.**
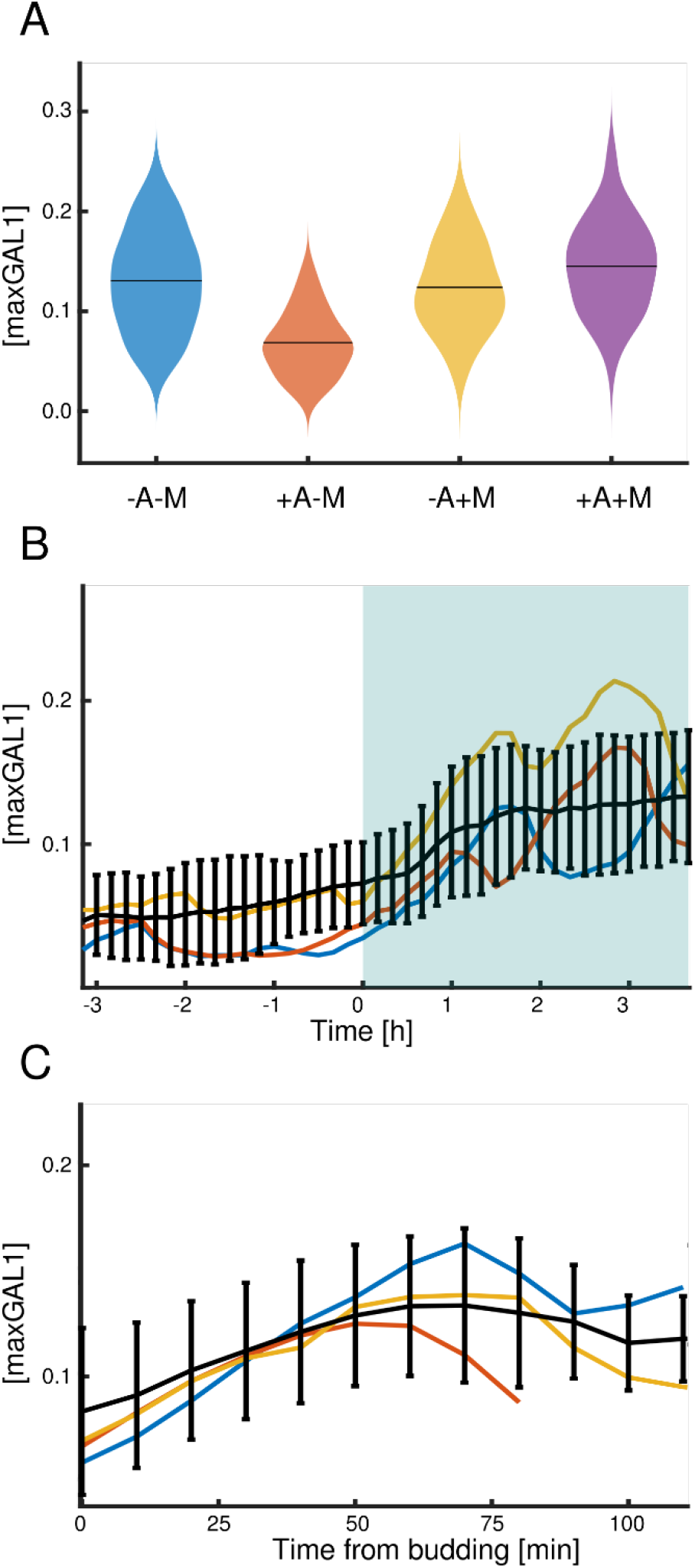
Dynamic properties of the *ARG3* promoter. A: Mean activity of *ARG3pr* in different media. +A or +M denote 10x concentrations of arginine or methionine, respectively, and -A and -M denote the lack of arginine or methionine in the medium. Numbers of analyzed cells are n = 131 (-A-M), n = 79 (+A-M), n = 83 (-A+M), n = 110 (+A+M). B: Time courses of ARG3-yEVenus-PEST activity in medium lacking methionine. The switch from +A to -A occurred at 0 h. Black line represents the average of the cells’ fluorescence levels and colored lines represent examples of single-cell fluorescence time courses. Number of cells present at t = 0 h is n = 51. C: Alignment of the single-cell trajectories (n = 17) using the time of budding shows that *ARG3pr* is likely cell-cycle regulated. In all panels, fluorescence is normalized with respect to steady-state levels of *GAL1pr* induction. Error bars indicate the standard deviation (SD).

Negative auto-regulation such as the repression of *ARG3* transcription by arginine is present in many other anabolic processes. Examples include the control of *LEU2*^98^, *URA3*^99^, *LYS20*^100^, and *MET3*^42^. On the other hand, the induction of *ARG3pr* by methionine was more puzzling since the biosynthesis of methionine and arginine are not obviously linked. We speculate that this is due to methionine serving as a global anabolic activation signal^101,102^. Gcn4, one of the *ARG3pr* regulators^92^, is essential for arginine biosynthesis and is induced in the presence of methionine^102^. It is unclear, however, what the functional role of the global regulation of metabolism by methionine is.

To characterize the dynamics of arginine-controlled switching between the off state (in -M+A medium) to the on state (in -M-A medium), we analyzed the *ARG3pr-yEVenus-PEST* expression time courses. *ARG3pr* responds quickly to the removal of arginine in medium lacking methionine (Fig. 11 B). Although at the population level, the *ARG3* promoter showed stable changes in activity in the presence of inducing medium, single-cell trajectories showed strong oscillations with a period close to the cell-cycle period, which was not detected previously^103^. The transcriptional regulation of *ARG3* involves the transcription factor Mcm1,^104^ which controls the expression of several cell-cycle periodic genes.^105,106^ When analyzing the cell-cycle-dependent trajectories of *ARG3pr* expression (Fig. 11 C), we observed that its expression peaked roughly after the middle of the cell cycle, potentially coinciding with peaks in other Mcm1-regulated genes such as *CLB2*.

Given that *ARG3pr* is activated by methionine, while *MET3pr* suppressed, they can be used jointly when inverted control of two circuits by a single input is needed.

### strongLOV: a more light-sensitive El222 mutant

We sought to broaden the repertoire of optogenetics systems used for control of cellular processes by creating and characterizing a variant of the El222-*LIP* transcription-factor-promoter system that is more sensitive to light.

We focused on identifying mutations that increase the light sensitivity of El222. The output of El222 is thought to depend on the time the protein spends in the active state, bound to the promoter.^57^ By comparing the dark-reversion kinetics and the amino acid sequences of El222 and other LOV-based photoswitches, we found several residues that are not present in El222 but are shared among other proteins with slower turn-off kinetics: Val71Leu, Ala79Gln, and Glu84Asp (amino acid identities given with respect to El222) (Fig. 12 A)^107^. Our hypothesis was that introducing a residue from the slow-cycling proteins (YtvA, AsLOV2 and VVD) into El222 would stabilize the light-activated state. A similar approach has been used to develop the AQTrip El222 mutant^59^, which incorporates the Ala79Gln^108,109^ mutation, among others. However, this mutant has an active state with an *in vitro* half-life of around 30 min, which may impede its applications in experiments where faster off switching is needed. Thus, we considered the other two candidates for mutations (Val71Leu and Glu84Asp). Given the proximity of Glu84Asp to the chromophore in the tertiary structure of the protein (Fig. 12 B) and the milder nature of the residue exchange (aspartic for glutamic acid), we decided to characterize the Glu84Asp mutant, which we named strongLOV.

**Figure 12.**
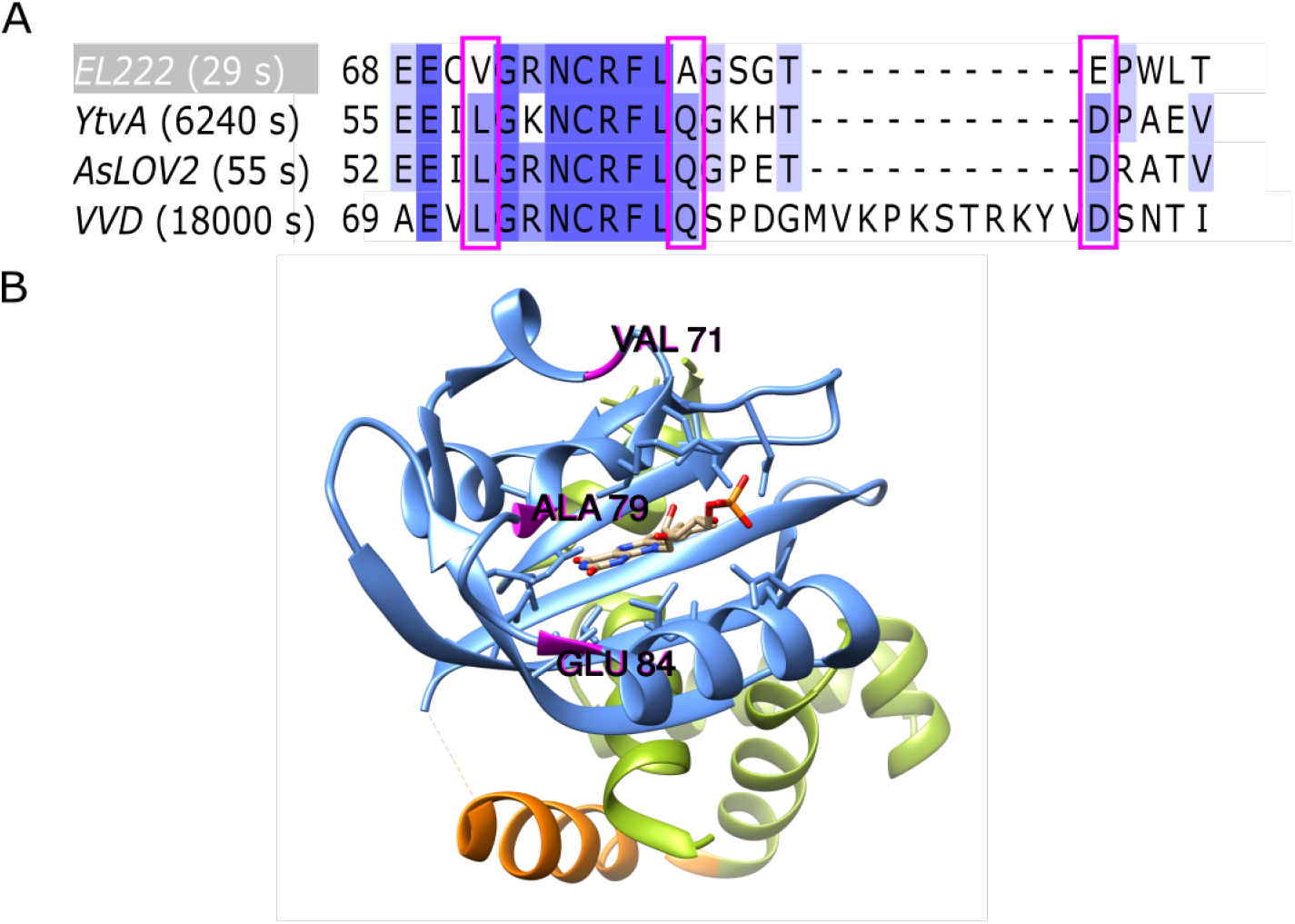
A comparison of LOV-domain sequences suggests candidates for mutations that stabilize the active state of El222. A: Multiple-sequence alignment of LOV-domain proteins with characterized dark-reversion kinetics. Amino acids are colored based on their similarity to the consensus sequence. The numbers next to the protein names indicate the half-life of the active state^107^. The residues that are conserved between YtvA, AsLOV2, and VVD but not present in El222 are marked by the pink boxes. (There are no more such residues outside of the subsequence of El222 shown.) B: The position of the identified residues (pink) in the El222 structure. The LOV domain is shown in blue, while the Jα helix and the HTH domain are shown in orange and green, respectively. The light-absorption center, flavin-mononucleotide chromophore, is shown in the middle of the structure.

To compare the *in vivo* performance of strongLOV to wild-type El222, we introduced both transcription factors in single copies into the yeast genome harboring a single copy of *LIP-yEVenus-PEST* as a transcriptional reporter. We first measured the induction of both strains under low light conditions (20% of maximal light intensity). strongLOV indeed responded more strongly to light activation, with an increased maximal intensity of around 5.5x (Fig. 13 A). When activated by high-intensity light (80% of maximal intensity), strongLOV showed induction levels comparable to wild-type El222 (Supplementary Fig. 11), suggesting saturation under strong light induction.

**Figure 13.**
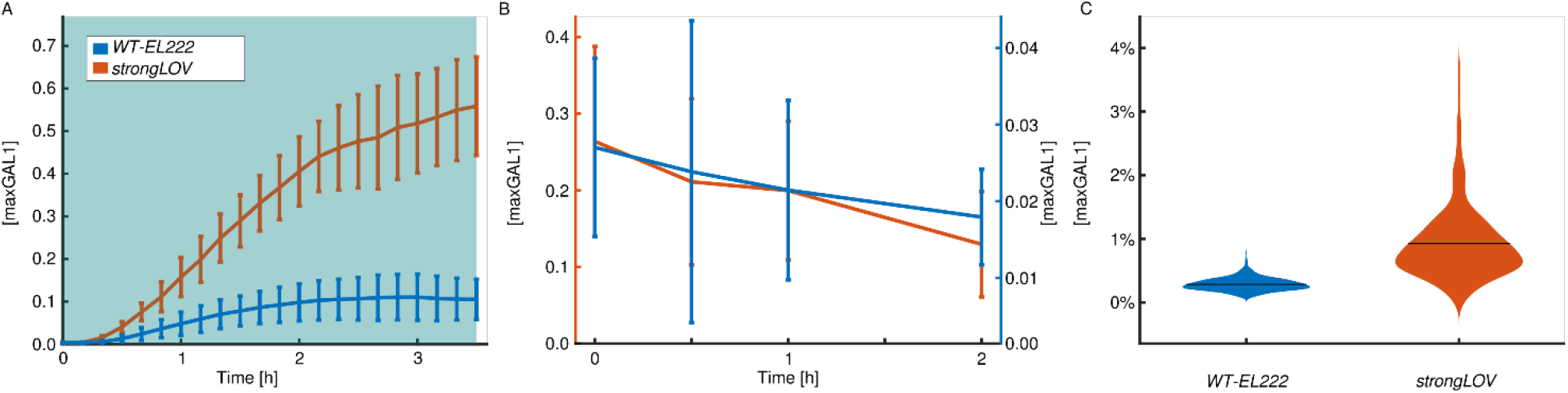
The strongLOV variant responds more strongly compared to WT-El222 under low light conditions. A: Induction of wild-type and mutant El222 using light with 20% of maximal intensity (standard deviation shown around each timepoint). The blue background denotes the presence of continuous light. B: Turn-off dynamics obtained by sampling cells with *strongLOV* and *EL222* in bulk culture, which were previously pulsed with blue light for 1 min every 15 min, which is much weaker than 20% light induction in panel A. The light is turned off at timepoint 0, standard deviations around each timepoint shown. Note the different y-axes for strongLOV (orange, left) and WT-El222 (blue, right). C: Basal activity measured with the *LIP-yEVenus* (no *PEST*) reporter strain. Horizontal bars denote mean values. Numbers of analyzed cells are given in Supplementary table 14.

To determine the turn-off dynamics of strongLOV, we could not use fluorescence microscopy, which continuously excited the system during measurements (Supplementary Fig. 11). We thus performed experiments in liquid culture where after long (> 12 h) log phase growth under low duty-cycle pulsing light (1 min of blue light every 15 min), we turned off the blue light source and monitored the dynamics of the fluorescent reporter by sampling the population of cells at different timepoints (Fig. 13 B). We observed a decline of the strongLOV activity with kinetics similar to WT-El222.

To measure the leakiness of strongLOV we introduced it in a strain harboring the transcriptional reporter without the *PEST* sequence, as before. We observed a mean increase in the leakiness of the mutated protein of 3.2x compared to El222 (Fig. 13 C).

Taken together, these results show that the newly described Glu84Asp mutation effectively increases the sensitivity of El222 but also increases its leakiness.

### Multidimensional trade-offs

Different experiments might require systems with different maximal levels of induction, or may tolerate different levels of leakiness or growth burden. To show how the multidimensional characterization presented here highlights the drawbacks of the different inducible systems for budding yeast, we plotted the relationship between maximal levels of induction, leakiness, delay upon induction, and growth data (Fig. 14). Strong induction systems such as El222-*LIP* induced at 80% of maximal light strength and *GAL1pr* are associated with slow cellular growth likely due to phototoxicity and a suboptimal carbon source, respectively. The weaker promoters *tetOpr*, *MET3pr*, *GALL*, and *CUP1pr*, either show substantial levels of leakiness (*tetOpr*) or show fluctuations (unstable expression) in time (*MET3pr*, *GALL*, and *CUP1pr*). The new strongLOV system is induced by less intense light; thus, it resolves the trade-off between phototoxicity and strength of induction – but has more leakiness in the dark.

**Figure 14.**
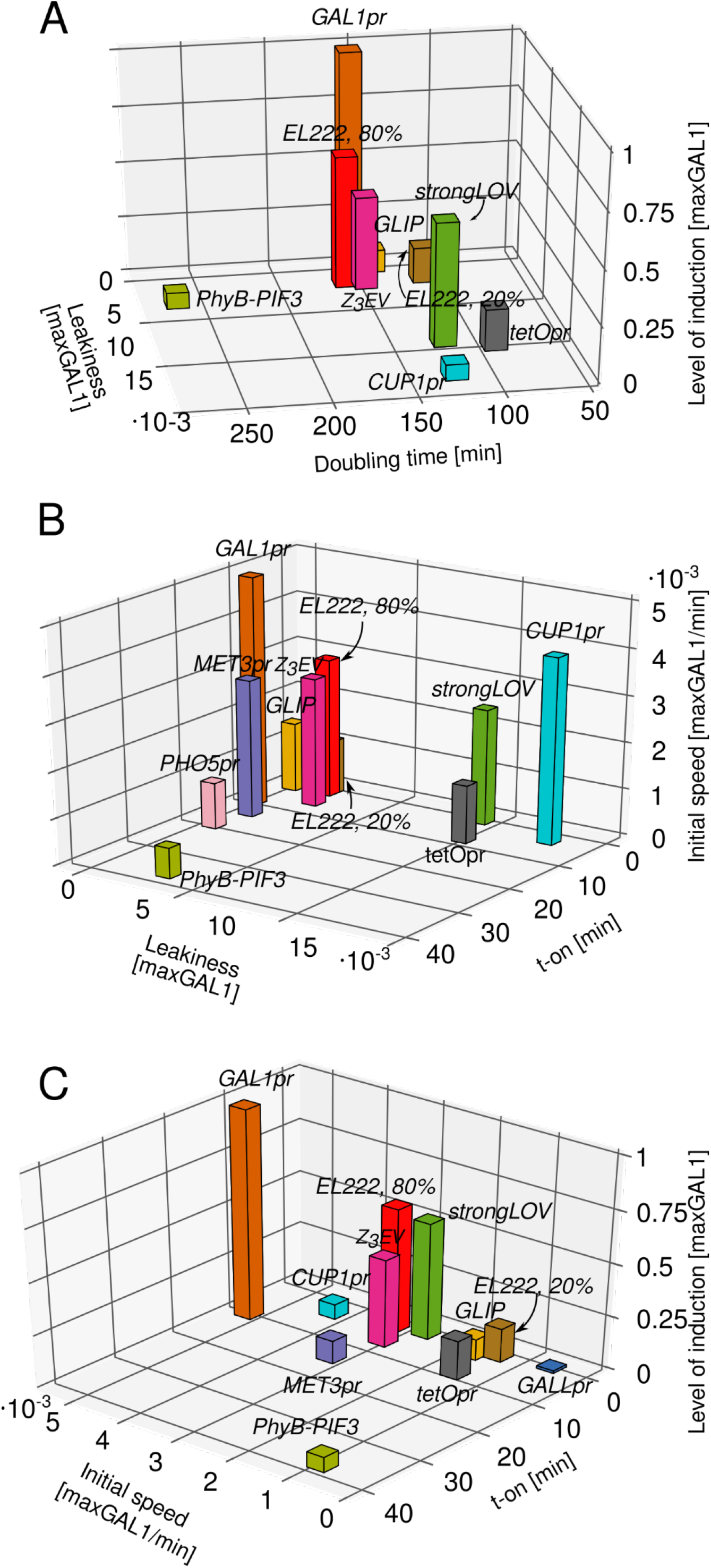
Multidimensional benchmarking of inducible systems illustrates performance trade-offs. The underlying data is the same as in Figs. 5, 6 and 7. Levels of induction shown in panels A and C are the steady-state levels of induction, except for *PHO5pr*, for which we show the level of activation at t = 3.5 h. *EL222* refers to the WT-El222 transcription factor induction of *LIP* under 20% or 80% light intensity, *strongLOV* refers to Glu84Asp El222 induction of *LIP* under 20% light, while *GLIP* is induced by El222 under 80% light. Numbers of analyzed cells are given in the Supplementary Table 14.

### Experimentally tuning the time between Start and mitosis

One of the goals of synthetic biology is to engineer complex artificial cellular behaviors. This often requires multiple inducible systems to be controlled simultaneously with high temporal precision. A scenario where such precision is necessary is in controlling inherently dynamic systems such as the cell cycle. Here, we control the lag between cell cycle Start and mitosis by independently inducing the expression of Start and M-phase cyclins in succession.

Cyclins are regulatory proteins, which, together with the cyclin-dependent kinase Cdk1, control the processes required for cell cycle initiation, progression, and exit.^110^ G1 cyclin (*CLN3*) and G1/S cyclins (*CLN1,2*) trigger entry into the cell cycle, while M phase cyclins (*CLB1, CLB2*) are needed for mitosis.^110^

In order to control entry into the cell cycle, we used a *MET3pr-CLN2* construct, which controls cell cycle Start in a strain in which all other Start cyclins have been deleted (*cln1-3Δ*).^44^ To tune the expression of the major mitotic cyclin *CLB2*, whose rate of expression is known to be limiting for the speed of mitosis^111,112^, we put an undegradable version of this cyclin (*CLB2kd*)^113^ under the control of El222-*LIP*. We chose El222-*LIP* among other tested systems because of its short response time (t-on), monotonicity, and relative strength. In addition, El222-*LIP* induction can be modulated by varying the light intensity^6,29^. *LIP-CLB2kd* is solely responsible for mitotic entry in a strain in which both mitotic cyclins were deleted (*clb1,2Δ*). This strain is kept viable by a *GALL-CLB2* construct in galactose medium prior to the measurements.^112^ Cells lacking all G1 and G1/S cyclins are arrested in G1 phase, while cells lacking *CLB1* and *CLB2* are arrested prior to M phase.

Before inducing *LIP-CLB2kd*, we ran cells through a sequence of media switches designed to deplete the Clb2 protein expressed from *GALL-CLB2*. We call these steps the Clb-depletion protocol^114^ (Fig. 15 A): After growing cells in G-Met (synthetic complete medium containing galactose and no methionine) medium, where the *MET3pr-CLN2* and *GALL-CLB2* constructs kept cells viable, we synchronized the population by switching the medium to G+Met (in which cells arrest in G1) for 2 h. Then, the medium was switched back to G-Met for 50 min, and cells restarted the cell cycle. After this, *MET3pr-CLN2* was turned off to prevent a second cycle, and after 20 min, *GALL-CLB2* was turned off by switching to medium that contains glucose instead of galactose, roughly at the end of mitosis to coincide with the time of activation of the Clb inhibitors Cdh1 and Sic1. After Clb depletion, we released cells from the G1 arrest by switching the medium from +Met to –Met and began the main experiment by turning on the light source, which activated the *LIP-CLB2kd* construct.

**Figure 15.**
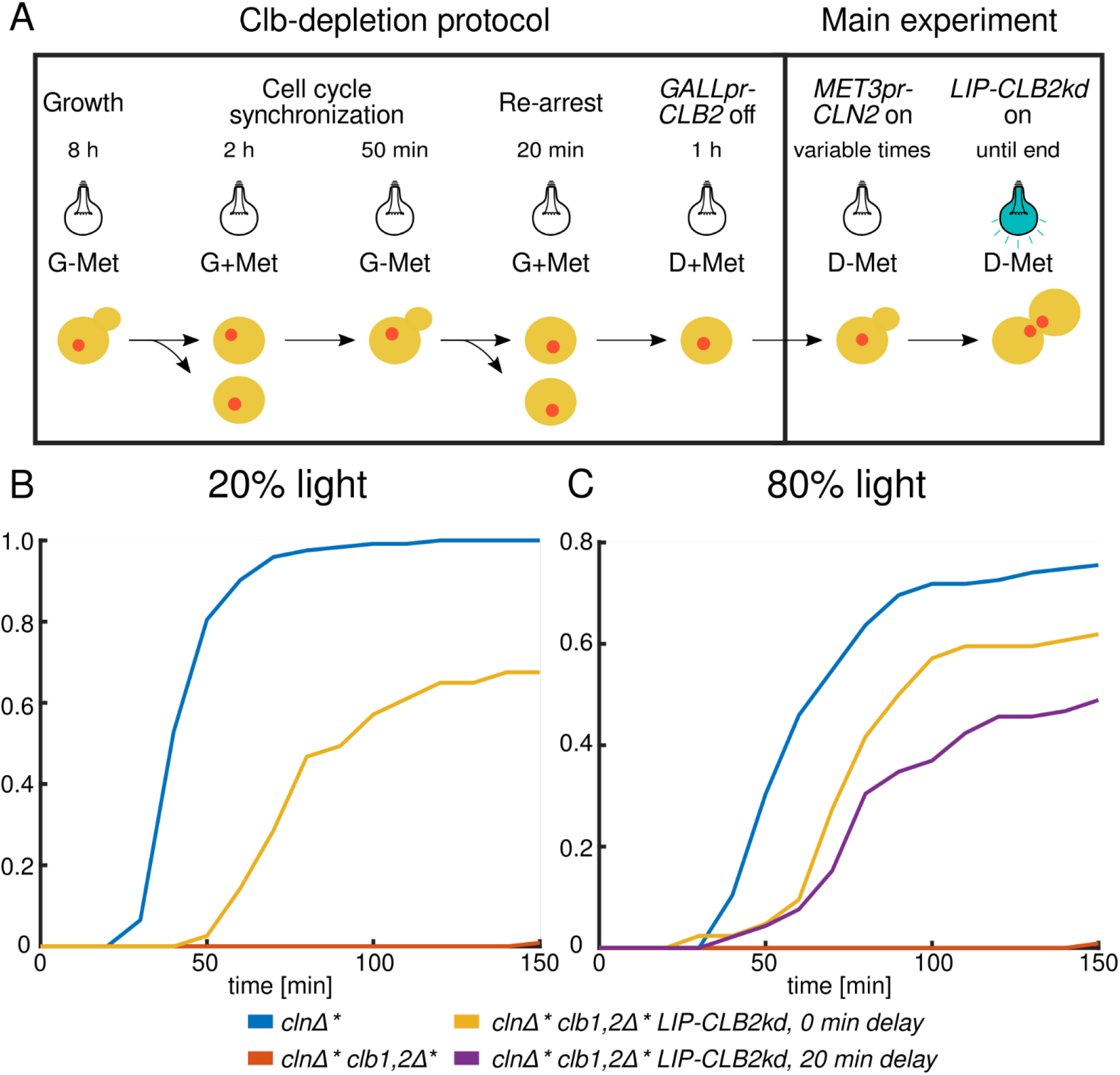
Independent triggering of cell cycle Start and mitosis to simulate wild-type timing. A: Illustration of the protocol. B: Budding-to-anaphase duration with 20% diascopic light intensity. C: Budding-to-anaphase duration with 80% light intensity. *clnΔ\** denotes *clnΔ MET3pr-CLN2pr* while *clb1,2Δ*\* denotes *clb1,2Δ GALL-CLB2*. The same experiment with control *clnΔ* clb1,2Δ\** cells (without the *LIP-CLB2kd* construct) is shown in panels B and C. Number of scored cells shown in Supplementary Table 18.

We varied *LIP-CLB2kd* expression by changing the light intensity that cells were exposed to and by changing the delay between the –Met pulse, which triggered entry into the cell cycle, and the light pulse, which triggered mitotic entry. Given that the presence of Clb2 around Start is known to block budding^115^, we started the induction of *LIP-CLB2kd* either after or at the same time as *MET-CLN2*. To monitor the dynamics of the cell cycle, we included the fluorescently labeled *HTB2-mCherry*^116^ construct in our strains, which marked the position of the nucleus throughout the cell cycle. For cell cycle timing, we measured the time from bud appearance to the separation of the fluorescently labeled nuclei in anaphase.

First, we applied 20% of the maximal light intensity to induce *LIP-CLB2kd* expression (Fig. 15 B) simultaneously with *MET3pr-CLN2* activation. Around 60% of cells with the *LIP-CLB2kd* construct that budded successfully finished mitosis. The effect was due to timely expression from the *LIP-CLB2kd* construct since residual Clb2 from the *GALL-CLB2* construct was not enough to drive cells through mitosis; this was verified by detecting almost no nuclear divisions in cells without the *LIP-CLB2kd* construct (Fig. 15 B and C). The Clb-depletion protocol had indeed removed Clb proteins effectively. However, their speed was slower than cells with wild-type *CLB1,2* (difference of the mean: 39.7 min).

In order to observe the effects of stronger *LIP-CLB2kd* induction, we applied light with 80% of the maximal intensity (Fig. 15 C) simultaneously with *MET3pr-CLN2* activation. This decreased the difference in time from bud emergence to nuclear separation compared to wild-type *CLB1,2* (difference of the mean: 16.8 min) with the proportion of *clnΔ* clb1,2Δ* LIP-CLB2kd* cells that finish mitosis similar to the experiment with 20% light intensity. Also, we could modulate the dynamics of mitosis progression by delaying the *LIP-CLB2kd* pulse relative to the *MET3pr-CLN2* pulse by +20 min. However, the proportion of wild-type *CLB1,2* cells that finished mitosis in the presence of 80% light was reduced, from around 100% in the presence of 20% light to around 75% in the presence of 80% light. This suggests that the higher intensity of light was toxic for cycling cells. Thus, different underlying effects may cause cells with the *LIP-CLB2kd* construct to not finish mitosis with 20% or 80% light: inappropriate rate or timing of the *CLB2kd* pulse in the former case and light toxicity in the latter.

## Discussion

Quantitative characterizations of inducible systems are needed to guide experimental designs. Here, we systematically and comprehensively benchmarked the characteristics of inducible systems in budding yeast. For some inducible systems, the level of activity is known to depend on the level of the inducer. Given that the input-output relationships for most of the tunable systems investigated here are known to be highly sigmoidal^40,75,117^, we focused on the characterization of the systems’ dynamic properties, not steady-state dose-response relationships.

We showed that the maximal levels of induction of these systems span a >50 fold range, suggesting that the library described here is diverse enough to guide different choices of inducible systems, at least, with respect to induction strength. With kinetic and steady-state parameters taken together, none of the tested systems performed optimally, emphasizing the need for the multidimensional characterization and the need for the development of novel tools for the precise dynamic control of cellular processes.

Although the naturally occurring yeast promoters can impose pleiotropic effects, our analysis of fundamental leakiness shows that, in cases where there are molecular mechanisms that actively inhibit their transcription (such as for *GAL1pr* and *MET3pr*), these promoters can exhibit substantially lower leakiness than other systems. This also validates the strategy for reducing leakiness of synthetic promoters by borrowing the regulatory sequences that keep the naturally occurring promoters off, as in the case for *GLIP*. However, to achieve orthogonality of leakiness to metabolism, more elaborate constructs are needed, such as the synthetic systems that repress the transcription of their own activators, as in the newly developed self-repressible Tet-Off system^118^.

A benchmark has the benefit of making the characteristics of a comprehensive set of inducible systems that different subgroups of researchers may or may not know about, in principle, known to all. In addition, the quantitative nature of our benchmark, using an intuitive unit of activity (maxGAL1), enables more precise experiments. So, even some of the better known shortcomings of inducible transcriptional systems, e.g., the carbon source-dependent decrease of growth rates (*GAL* systems) and the high leakiness of the tetracycline-inducible system, can be accounted for precisely now. For example, the tetracycline-inducible system in its ‘off’ state can be used as a constitutive promoter that is roughly as strong as *GALL* in the ‘on’ state. Furthermore, at a qualitative level, many features of the systems we analyzed were unpublished, for example, the two-step activation of *PHO5pr*, small dynamic range of *CUP1pr*, long time delay after deactivation of *tetO* and Z_3_EV, non-monotonic activation of *MET3pr* and *GALL*, high stochasticity of PhyB-PIF3 and *PHO5pr*, and high relative leakiness of PhyB-PIF3 and *CUP1pr*.

The analysis of some of the inducible systems also adds to the description of their mechanisms. We worked out the different levels and sources of the *GAL1*-based promoters’ leakiness. For example, we demonstrated that *GLIP* as a synthetic *GAL1*-based system is affected by the carbon source and requires glucose to keep it tightly off. Furthermore, by inducing *MET3pr* in a strain lacking Met17, an enzyme in the methionine biosynthetic pathway, we showed that the internal production of methionine contributes to the decline in *MET3pr* activity in the absence of external methionine. Through a comparison of the transcriptional Gal4-based PhyB-PIF3 system with the PhyB-PIF3 system for subcellular localization, we found that the large noise levels come from the Gal4 functionality, not the interaction of PhyB and PIF3.

We introduced strongLOV, a mutant El222 transcription factor that requires less light for the same level of activity and thus could reduce phototoxicity. As the El222 optogenetic system is extensively used in organisms other than budding yeast such as mammalian cell lines^119^, bacteria^120^, zebrafish^58^, and plants^121^, the new mutation described here ought to be useful for light control experiments in different fields of biology, as well as contribute to further understanding of LOV-domain proteins photochemistry.

The comparatively little explored *ARG3* promoter showed an interesting OR gate behavior as well as the opposite activation with respect to methionine compared to *MET3pr*. Although dynamic control using *ARG3pr* may be impeded by its small dynamic range and high leakiness, its level of expression in the ON state is comparable to *MET3pr*, which is useful in scenarios where this is the physiological level of expression.

Lastly, we showed that with two fast-acting inducible systems, we could simulate the succession of cell cycle Start and mitosis with nearly wild-type timing.

## Methods

### Plasmid library construction

All plasmids were constructed and propagated using *E. Coli* DH5α. DNA digestion and ligation were performed using restriction endonucleases and T4 DNA ligase from New England Biolabs (USA). The *promoter-yEVenus-PEST* library was constructed by cloning different promoter sequences upstream of the *yEVenus* ORF using *PacI* and *BamHI* restriction enzymes. All PCRs were performed with Phusion Polymerase (New England Biolabs, USA). All constructs were verified by Sanger sequencing (Microsynth AG, Switzerland). Summary and details of the construction of plasmids used in the study are given in Supplementary Table 1.

### Strain construction

Wild-type haploid *W303* budding yeast strains (*MATa ade2-1 leu2-3 ura3-1 trp1-1 his3-11,15 can1-100*) were transformed with plasmids with the inducible *promoter-yEVenus-PEST* constructs by digesting the plasmids with *StuI* endonuclease inside the *URA3* gene. Transformations were performed using the standard lithium acetate method^122^ and transformed strains were selected using -Uracil dropout plates. For systems involving synthetic transcription factors (light-, doxycycline-, and estradiol-inducible systems), constructs encoding transcription factors were transformed in a strain of the opposite mating type from the strain containing the *promoter-yEVenus-PEST* construct and the transcription factor plasmids were integrated into the *HIS3* locus. The two strains were then crossed and the resulting progeny that contained both transcription factor and *promoter-yEVenus-PEST* constructs were selected and used in further experiments. Plasmid integration and construct activity were verified by fluorescence microscopy after the appropriate induction of the constructs. Strains that showed fluorescence were screened for single-copy integrations using polymerase chain reaction (PCR) with primer sets that allowed one or several copies of the construct in the genome to be distinguished (Supplementary Note 4). Some researchers used the Gal4-based PhyB-PIF3 system in the *gal4Δ gal80Δ* background^54,55^. However, the system is shown to work well also in the absence of these two deletions^123^, and in our experiments we opted for the simpler version with the endogenous copies of *GAL4* and *GAL80* present. To remove the *PEST* sequence from strains that had the *promoter-yEVenus-PEST-ADH1t* construct, we created a *KanMX*-marked plasmid (pVG97) that, when cut with the *AfeI* restriction enzyme and used to transform strains with the *promoter-yEVenus-PEST::URA3* construct results in genomic *promoter-yEVenus-ADH1t*. *PEST* removal was confirmed by the absence of the functional *URA3* copy and by PCR in all constructed strains. Summary and details of the strain construction used in this study are given in Supplementary Table 2.

### Media and growth conditions

Cells were grown in CellASIC ONIX microfluidic plates for haploid yeast cells in media controlled by the ONIX2 microfluidics system (Merck, Germany). Details regarding the composition of the media used for different promoter induction experiments are given in Supplementary Note 1.

For experiments with light-induced *CLB2kd*, cells were first grown in G-M medium from a single cell to a colony for 8-12 h. After that, to ensure that no left-over Clb2 would affect the cell cycle in which the *LIP-CLB2kd* construct was induced, the Clb-depletion protocol^114^ was applied as described in the main text.

### Microscopy

Images were recorded using a Nikon Ti2-E microscope equipped with a 60x objective and a Hamamatsu Orca-Flash 4.0 camera. The microscope was operated using NIS-Elements software and the objective’s axial position was controlled by the Nikon Perfect Focus System. To reduce photobleaching of the fluorescent protein, images were taken every 10 min with 100 ms exposure time.

### Image analysis

Image analysis was performed using YeaZ, a Python-based tool for yeast cell segmentation^34^. Briefly, we first determined the boundaries of cells in phase-contrast images. The levels of fluorescence for each cell were then calculated as an average of the pixel intensities in the yellow fluorescence channel for pixels that were within the cell boundaries. For further analyses, we subtracted the autofluorescence of unlabeled wild-type cells from the fluorescence values of *promoter-yEVenus-PEST* carrying cells.

### Data analysis and modeling of gene expression

To extract parameters for the systems’ kinetic properties, we compared the single-cell expression data with a minimal model presented in Fig. 5 A. After solving the equations of the model, we obtained:

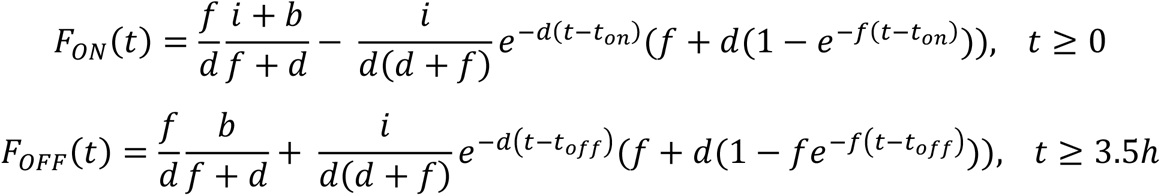

To simplify the fitting procedure, we further reduced the complexity of the two functions *F_ON_* (*t*) and *F_OFF_* (*t*) by expanding them in Taylor series and keeping only the first two terms:

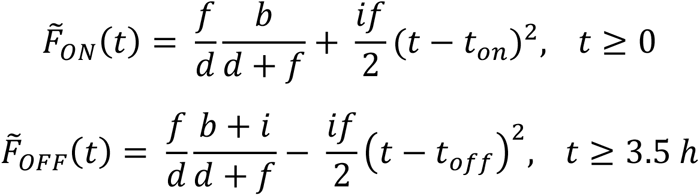

Based on 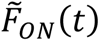 and 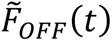, we could extract the induction parameters unambiguously. First, we extracted the term describing basal activity of the inducible transcriptional system, 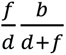, using the fluorescence values during the time prior to the induction (from t = -60 min to t = 0 min) for most of the systems, or at timepoint t = 0 h for the optogenetic systems *LIP*, *GLIP,* and PhyB-PIF3. Next, we fitted the part of the curve around the start of the induction period; this allowed us to extract the initial speed of the induction *i* and the delay of the transcriptional induction *t-on*. To unambiguously extract *i* and *t-on* from the second term of the Taylor expansion, we used a fixed value for the yEVenus maturation time *f* that we measured in an independent experiment (Supplementary Note 2). For most inducible systems, we fitted the timepoints from t = -50 min to t = 50 min. Exceptions were *GALL*, *CUP1pr*, which start showing non-monotonic activation soon after the initial rise and for which we used timepoints from t = -50 min to t = 30 min. For PhyB-PIF3, which turned on very slowly, we used timepoints from t = 0 to t = 60 min. For *LIP* and *GLIP,* we fitted the expression values from t = 0 min to t = 50 min. To extract *t-off*, we fitted the fluorescence values after removal of the inducer. For this, we used timepoints from t = 210 min to 270 min. Next, we extracted the degradation-and-dilution rate *d* from the part of the plots in Fig. 2 that correspond to the decay of the fluorescent protein by fitting to an exponential decay function. For this, we used the timepoints starting from an hour after the circuit was switched off, which is roughly four maturation half-times, ln(2)/*f*, so that the exponential term in *f* became negligible. That is, we used timepoints from t = 270 min to t = 390 min. Finally, to extract the basal activity parameter *b* from the fitted 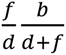 term of the Taylor expansion of the turn-on dynamics, we used the previously extracted parameter *d*. We note that since *d* was close to zero for the two systems that do not turn off well (Z_3_EV and PhyB-PIF3), the extracted parameter *b* might not represent the systems’ leakiness well. However, we show their fundamental leakiness in maxGAL1 units in Fig. 5 E and Fig. 6 A. The model fits were obtained by minimizing the sum of the squared residuals using the fminsearch function in Matlab 2019a. The matlab code to carry out these fits is made available as detailed in Code Availability below. For examples of fits of single-cell time courses, see Supplementary Fig. 5.

For making violin plots, we used a bandwidth of 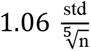, as suggested in ref.^124^ (std - the standard deviation, n - the number of elements in the set).

## Author contributions

VG performed microscopy experiments. VG and AS analyzed the microscopy data. VG performed modelling and analyzed the results. VG created genetic constructs and strains. VG and SJR wrote the manuscript. SJR supervised the work.

## Acknowledgment

We thank Dr. Enrico Tenaglia, and Roxane Dervey for help with the experiments and media preparation; Prof. Sebastian Maerkl, Dr. Evan Olson, and Shiyu Cheng for help with preparation of media for *PHO5pr* induction; Prof. Mustafa Khammash and Dr. Dirk Benzinger for providing plasmids and technical advice on the optogenetics measurements; Prof. Anton Khmelinskii for providing the strains with fluorescently labeled Arg3; Prof. José L. Avalos and Dr. Evan Zhao for providing *EL222* plasmids; Prof. Attila Becskei for providing the plasmid with *tetO* and *rtTA*; Prof. David Botstein for providing plasmids with the *Z_3_EV* system; and Prof. Peter Quail for providing plasmids with *PhyB-PIF3*.

## Code availability

The code generated for this study can be found at https://github.com/lpbsscientist/promoter-benchmark-model.

## Data availability

Plasmids generated in the study are deposited with Addgene (www.addgene.org). Strains generated in the study are deposited with National BioResource Project – Yeast database (https://yeast.nig.ac.jp/yeast/). The data generated in the study (coarse-grained parameters and single-cell data) are available at https://promoter-benchmark.epfl.ch/.

## Supplementary Information

**Supplementary Figure 1.**
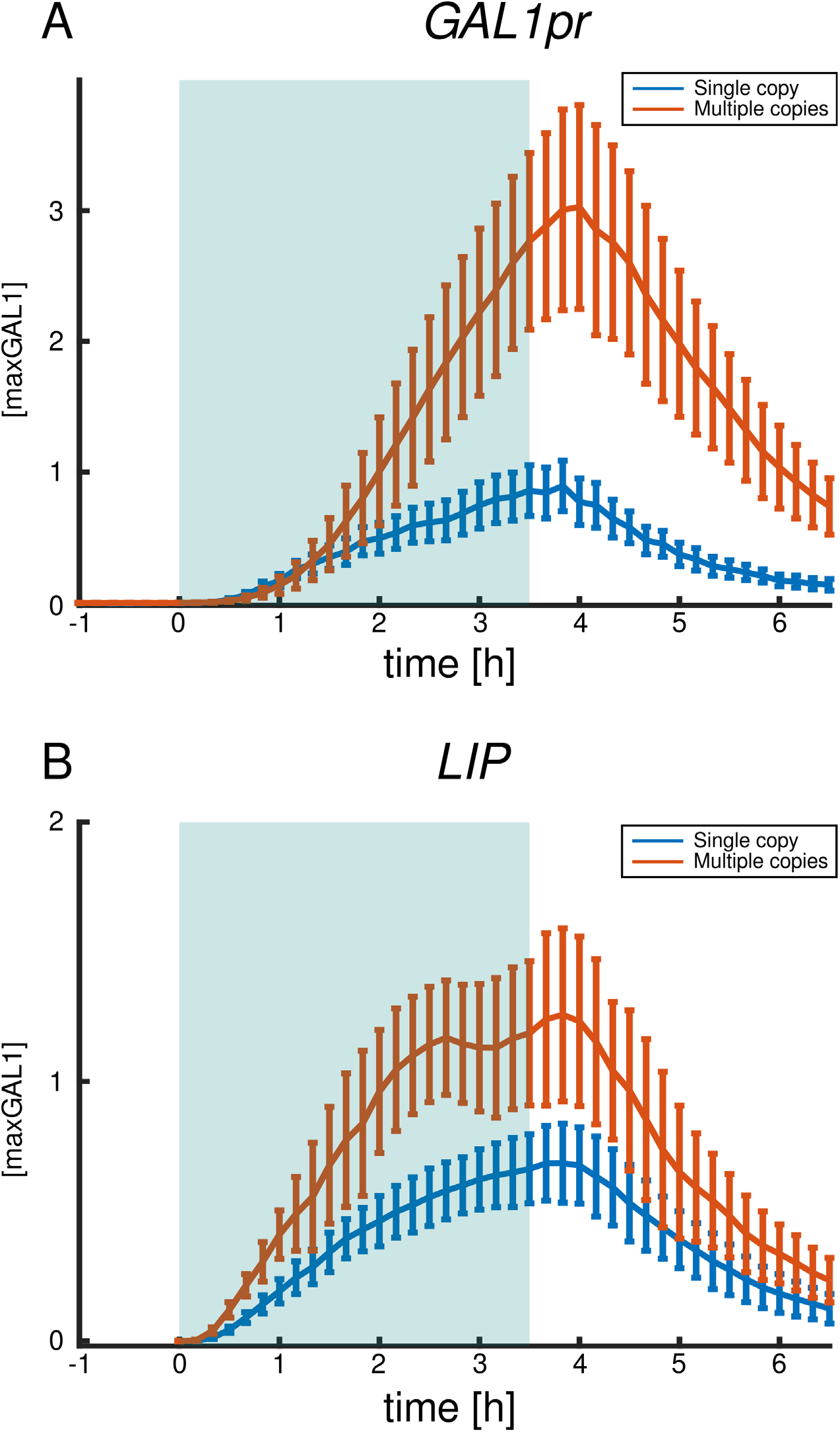
The number of copies of the reporter construct influences the observed activity level, rendering data from different sources with unknown reporter copy numbers not comparable. A: *GAL1pr-yEVenus* construct integrated as single (blue) or multiple copies (red) at the *URA3* locus. B: Similarly for El222-*LIP*. A, B: Time courses of the inducible system activity during dynamic perturbation. The blue background represents the induction period. Vertical error bars indicate the standard deviation around each timepoint.

**Supplementary Figure 2.**
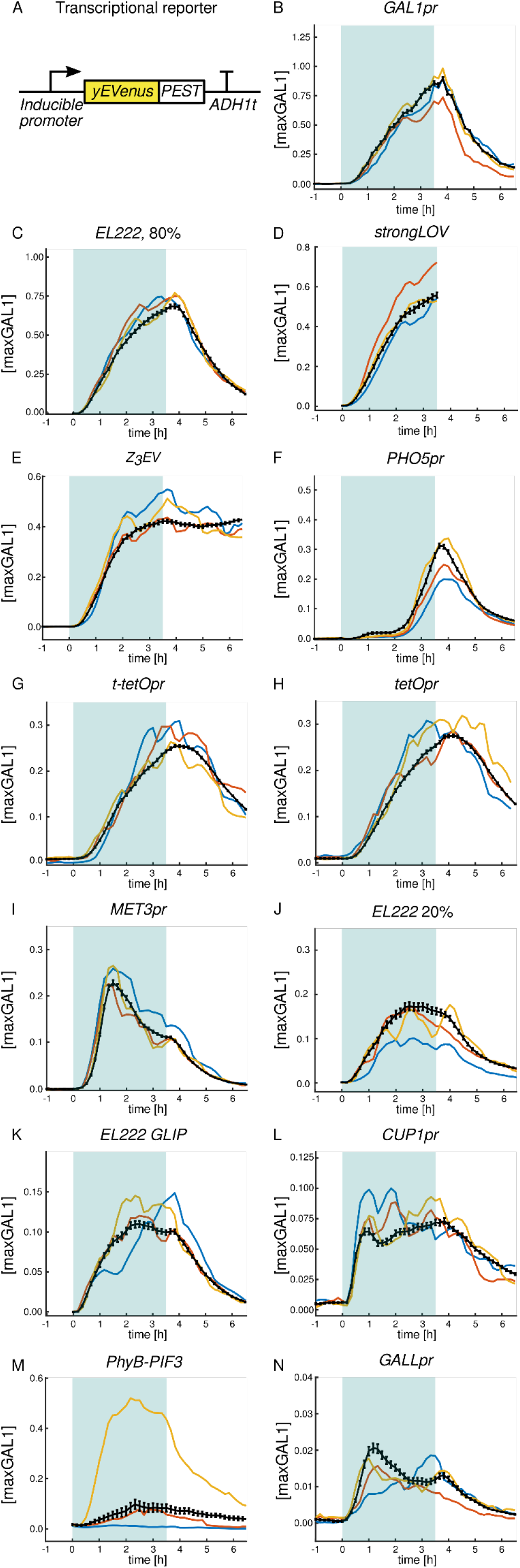
On and off dynamics of inducible systems with standard error of the mean (SEM) shown (instead of the standard deviation as in Fig. 2). The small SEMs show that the mean activity has been determined with high confidence based on the number of cells analyzed. A: The reporter for transcriptional activity consists of an inducible promoter and the fast-folding yellow fluorescent protein *yEVenus* gene fused to a constitutive degron (*PEST*) and the *ADH1* terminator. B-J: Time courses of activation and deactivation for different inducible systems sorted in descending order by peak strength. Induction starts at t = 0 h and finishes at t = 3.5 h. The blue background represents the induction period. Promoter activity is given in maxGAL1 units. Black lines show the average of the mean cellular expression and standard error of the mean. Colored lines show different representative single-cell time courses. For the light-inducible systems, fluorescence was not measured prior to induction in order to avoid possible activation by the light source used for fluorescent protein excitation. *EL222 20%, EL222 80%, strongLOV,* and *GLIP* are defined in the caption of Fig. 2. Due to the high sensitivity of strongLOV to the light used for fluorescent protein activation, microscopy measurement of the off dynamics was not possible; for turn-off experiments in bulk culture see Fig. 13.

**Supplementary Figure 3.**
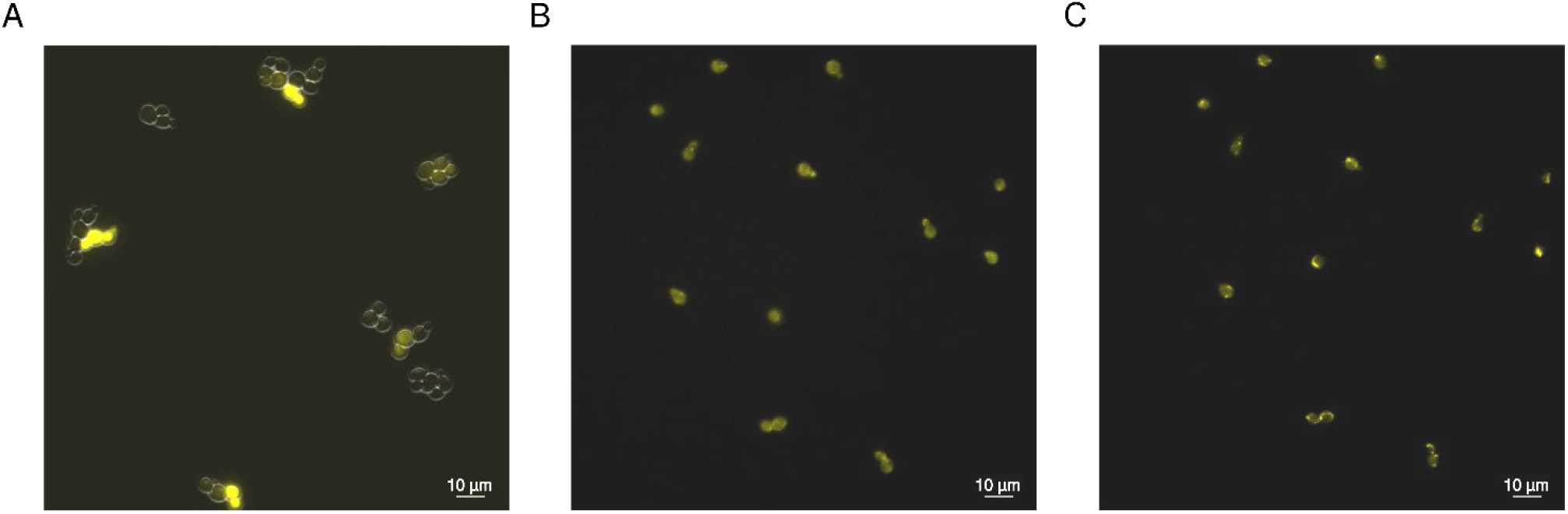
The PhyB-PIF3 system for transcriptional control of *GAL1pr* shows higher cell-to-cell variability in response to red light compared to the PhyB-PIF3 system used for subcellular relocalization. A: The PhyB-PIF3 system for transcriptional control. B, C: Using the same experimental setup, we measured the responsiveness of the PhyB-PIF3 system used for inducing mitochondrial localization of a yellow fluorescent protein. B: After incubation with PCB, cells with *PhyB-mCherry-Tom7* and *Bem1-mCitrine-PIF3*^125^ constructs were illuminated with far-red light for 30 s (diodes with 740 nm emission peak), and a snapshot of the off state was taken 30 s later. These cells show the off state of the system, where Bem1-mCitrine is allowed to assume its diffuse localization. C: Cells with the PhyB-PIF3-based mitochondrial tethering construct responded uniformly to red light by changing the location of Bem1-mCitrine-PIF3. Induction was performed by illuminating the cells with red light (650 nm emission peak) for 30 s and imaging after 1 min from the start of the induction to allow for localization (same cells shown as in panel B with the same normalization of the pixel intensity). Given the high reliability of the system shown in panels B and C, these experiments suggest that the stochasticity of the transcriptional PhyB-PIF3 system does not come from noise in the PhyB-PIF3 interaction.

**Supplementary Figure 4.**
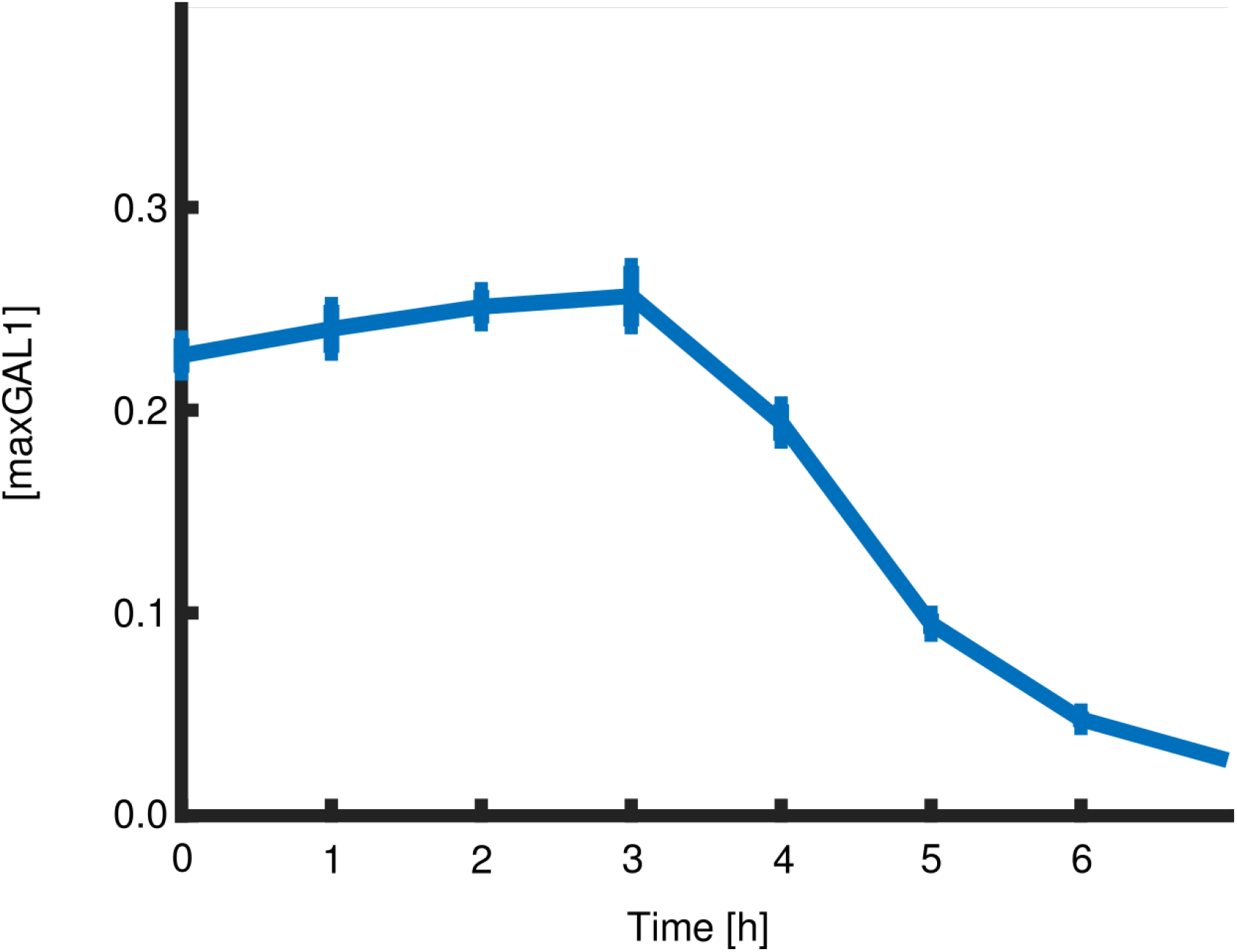
Expression under the control of Z_3_EV remains on for hours after estradiol is depleted from the media. The experiment was performed in liquid culture by transferring cells carrying *yEVenus-PEST* expressed by the Z_3_EV system induced with 0.5 µM estradiol to non-inducing media at t = 0. Vertical bars indicate the standard error of the mean (SEM). To make sure that no residual estradiol was carried to the off-state medium, we washed the cells 3 times by centrifugation and resuspension. The concentration of the inducer that we used before these washes was half of what is used in most of the experiments in the publication^19^ where the system was first presented.

**Supplementary Figure 5.**
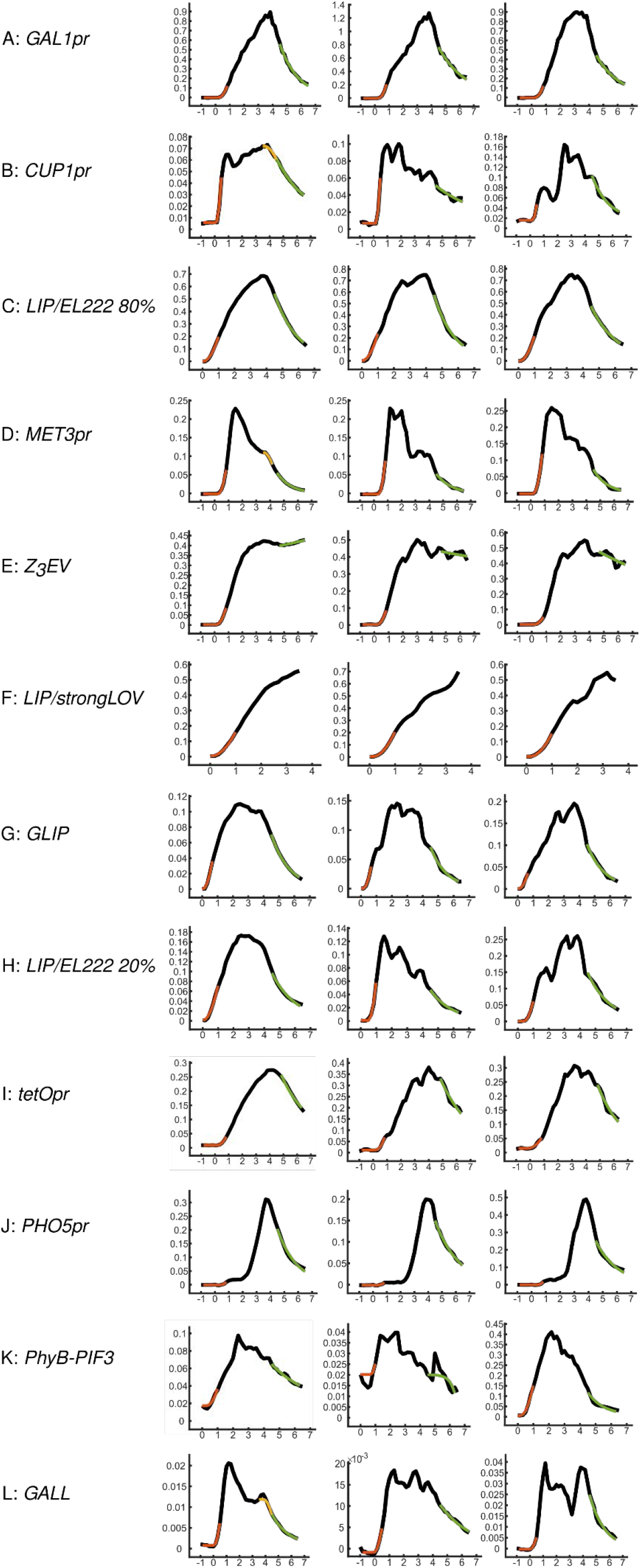
Single-cell fits. A-L: The first column shows the fits to the curve averaged across the population of the cells while the next two columns show examples of fits to single-cell time courses. The black curve represents the measured data; the orange curve denotes the fit to the initial part of the dynamics from which *b*, *tau-on,* and *i* are extracted; the green curve represents the fit to the part of the time course from which the degradation-and-dilution rates (*d*) are extracted. In the cases where the turn-off delay was estimated from the average time-course data (*CUP1pr, MET3pr, PHO5pr*, and *GALL*), the yellow curve shows the fit from which *t-off* is extracted. The y-axis on all plots is in maxGAL1 units, while the x-axis is in hours.

**Supplementary Figure 6.**
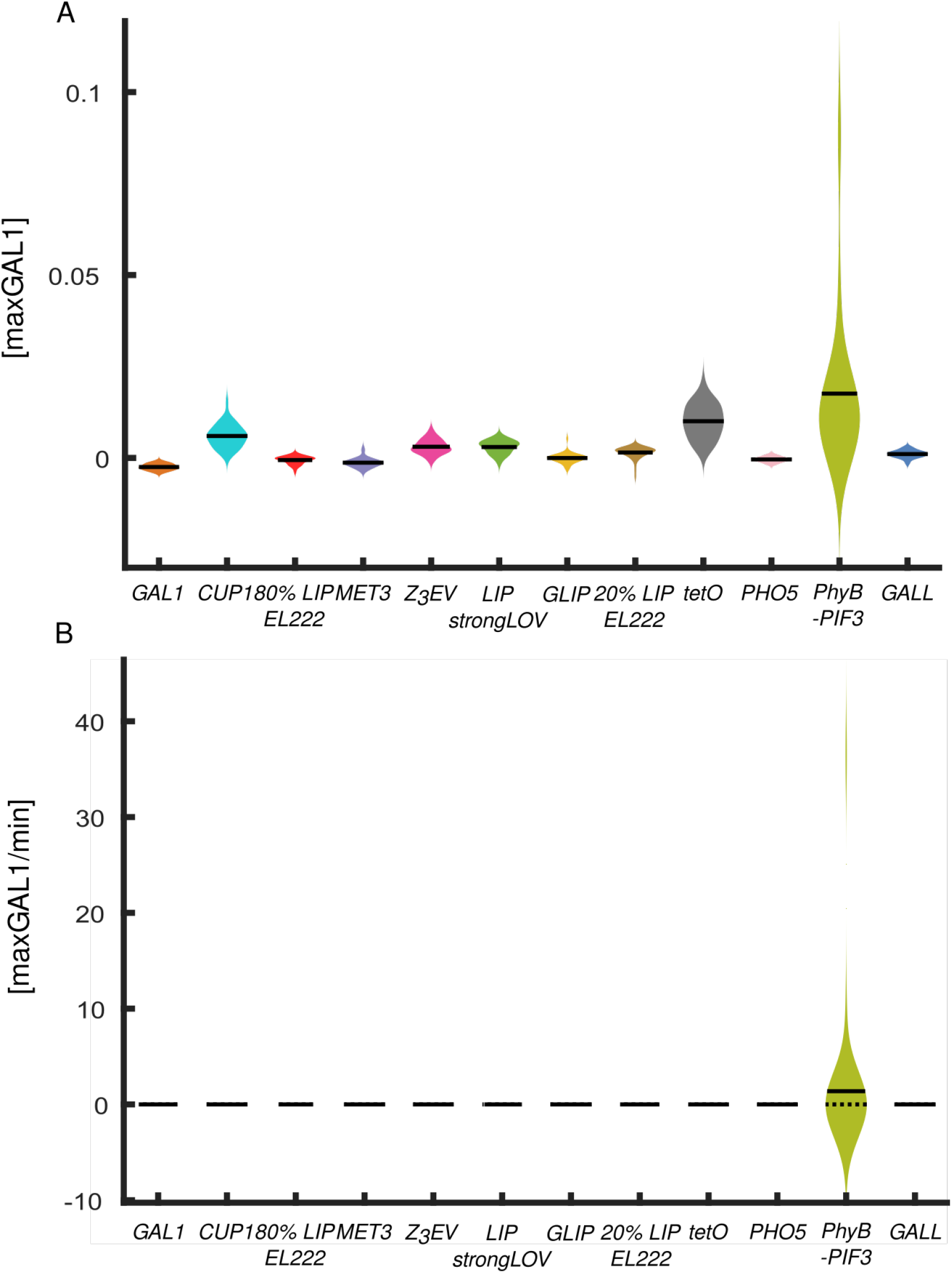
Single-cell characteristics of the inducible systems, shown with the whole population of *PhyB-PIF3* cells, including the outlier cells that were excluded in Fig. 5. A: Basal fluorescence (compare with panel E in Fig. 5). B: Basal level parameter (b) (compare with panel F in Fig. 5). Few (n = 3) out of 33 cells with the PhyB-PIF3 system have leakiness much higher than the mean of the population. This results in the higher estimated bandwidth used for plotting violin plots and obstructs the comparison between the systems, hence we excluded them from the main plot in Fig. 5 but show them here.

**Supplementary Figure 7.**
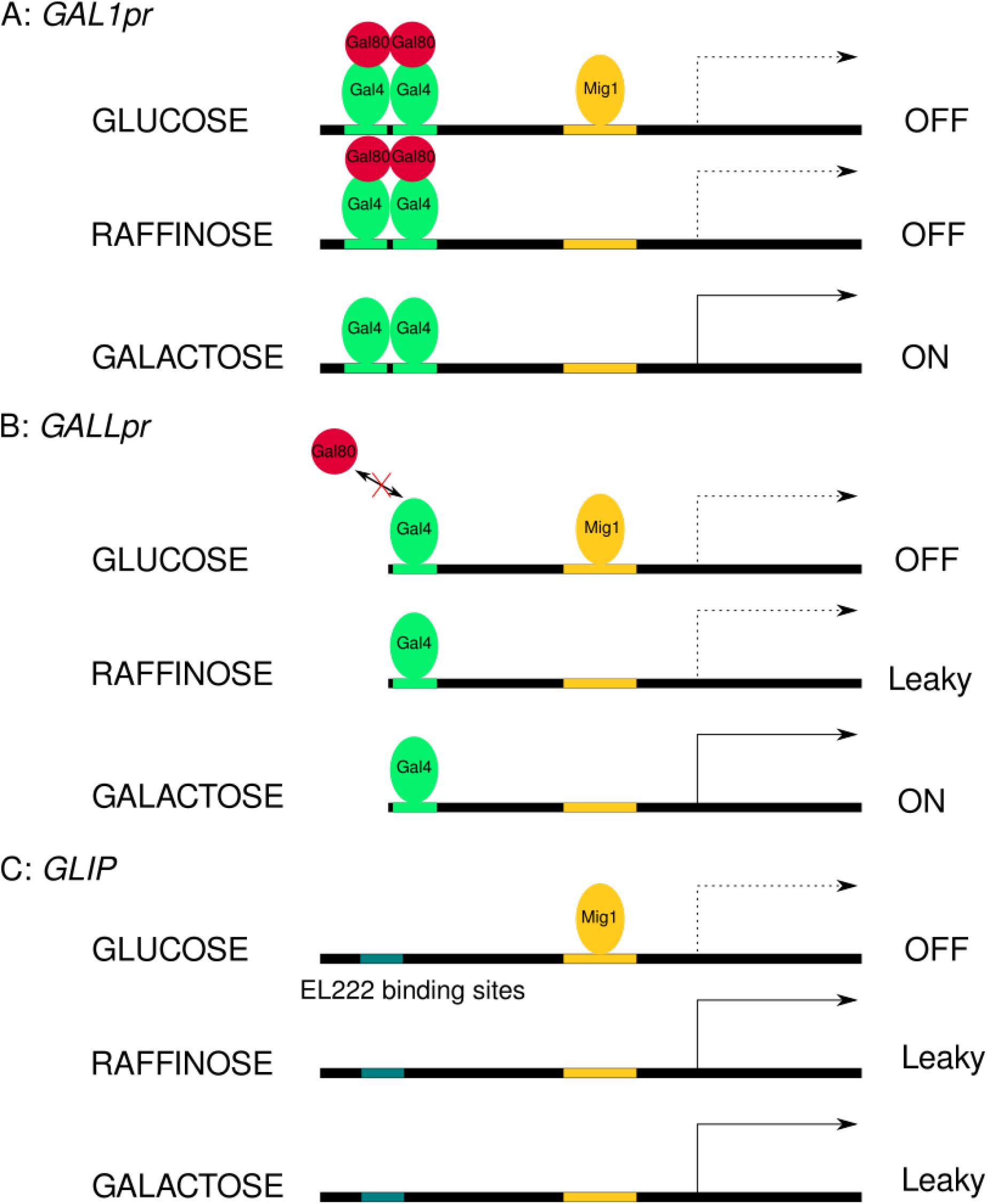
A simple molecular model based on past findings explains the leakiness of *GAL1pr*, *GALL,* and *GLIP* in different carbon sources. A: In glucose, *GAL1pr* is repressed by Mig1 and the Gal80 homodimer ^38^. In raffinose, *GAL1pr* is only repressed by the Gal80 homodimer^80^. Repression by Gal80 alone is strong enough to suppress leakiness below the detection limit of our transcriptional reporter without the *PEST* sequence. B: Once the repression of *GALL* by Mig1 is relieved in *raffinose*, it exhibits substantially higher leakiness compared to *GAL1pr*, presumably due to the less efficient binding of the Gal80 homodimer to the Gal4 monomer, as demonstrated previously^82^. C: The *GAL1pr*-based light-inducible promoter (*GLIP*) inherited its Mig1 binding sites from *GAL1pr*, which is reflected in low GLIP leakiness in glucose media. Inactivation of Mig1 in raffinose or galactose leads to the same level of leaky transcription presumably due to basal activity of the system.

**Supplementary Figure 8.**
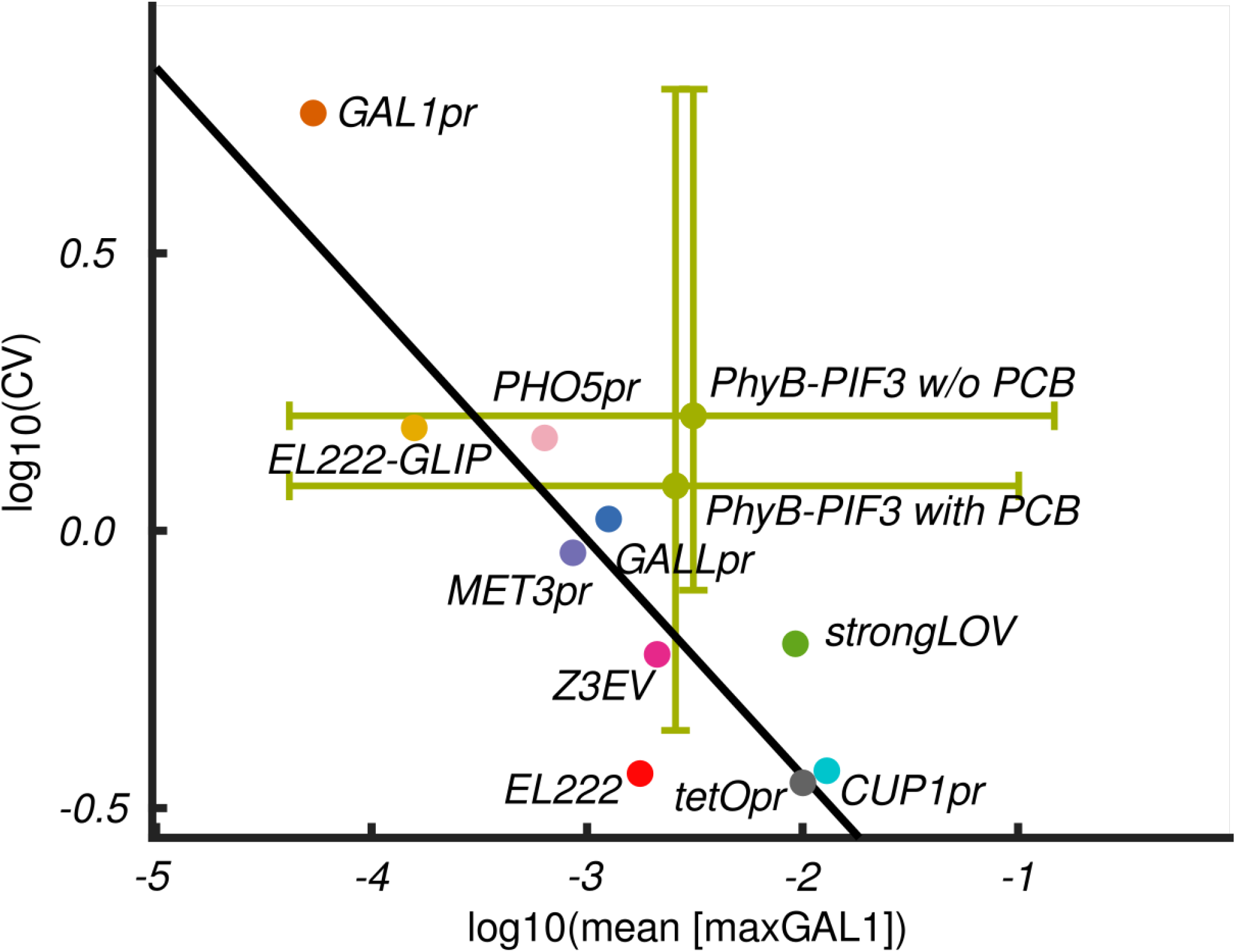
Log of noise (CV) versus log of mean (expressed in maxGAL1 units) of the leakiness of the different inducible systems in their ‘off’ states. In this figure, we show in particular that the high noise in the PhyB-PIF3 system is not due to variable or noisy PCB import or consumption. Noise is calculated as the coefficient of variation for the population. The bars around the PhyB-PIF3 values show 90% confidence intervals in both directions, reflecting very high variability in the off state. The linear fit is based on the data for all inducible systems except *PHO5pr* and PhyB-PIF3.

**Supplementary Figure 9.**
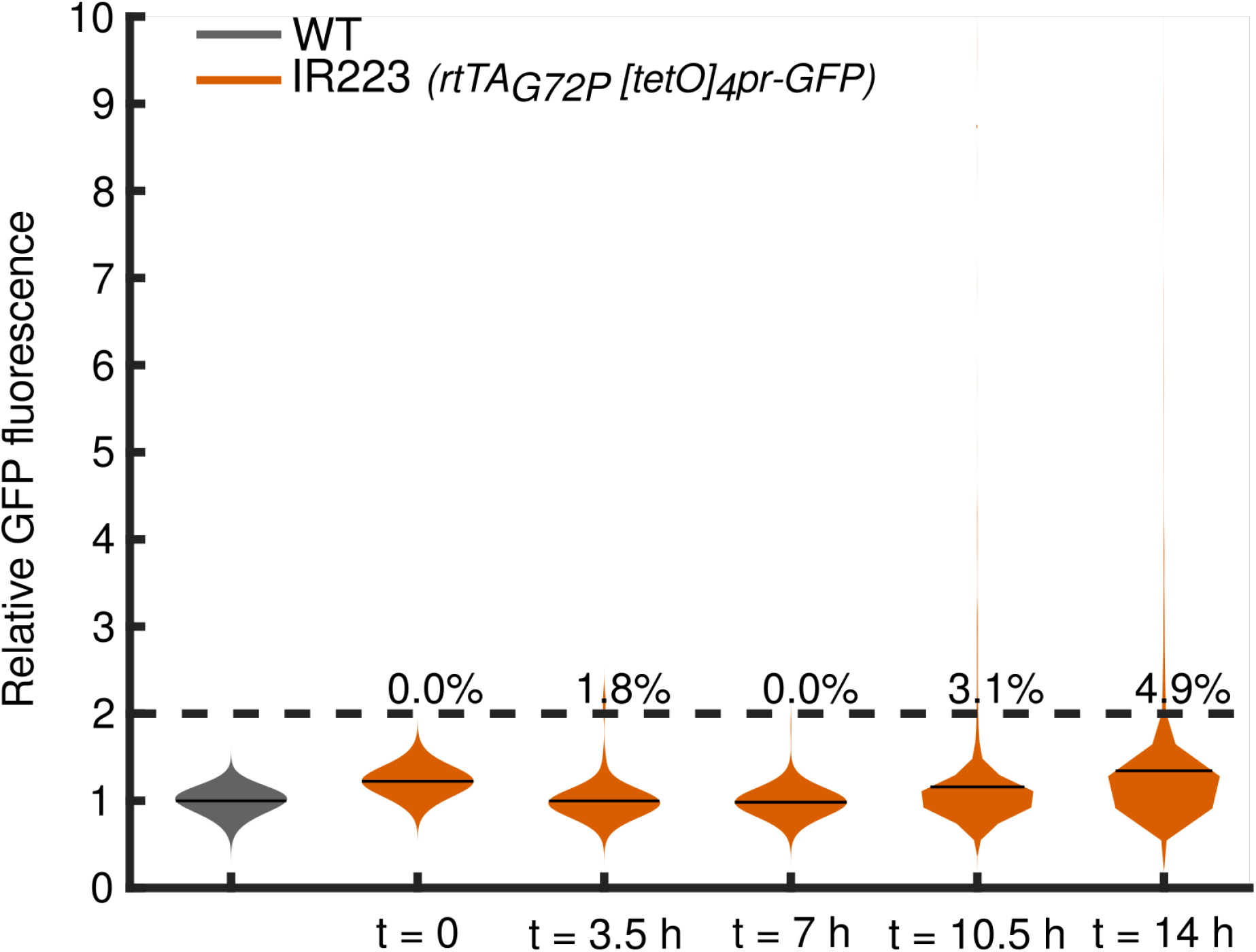
The IR223 strain containing the least leaky Tet-On system in ref. ^83^ was tested for induction for 14 hrs. We used the concentration of doxycycline tested in ref. ^83^ (100 mg/L doxycycline, a concentration 22.5x higher than the one used for the induction of the non-mutated Tet-On system). Values of GFP fluorescence are scaled relative to WT autofluorescence. Percentages indicate the fraction of cells that have fluorescence levels above 200% of WT autofluorescence (dashed line).

**Supplementary Figure 10.**
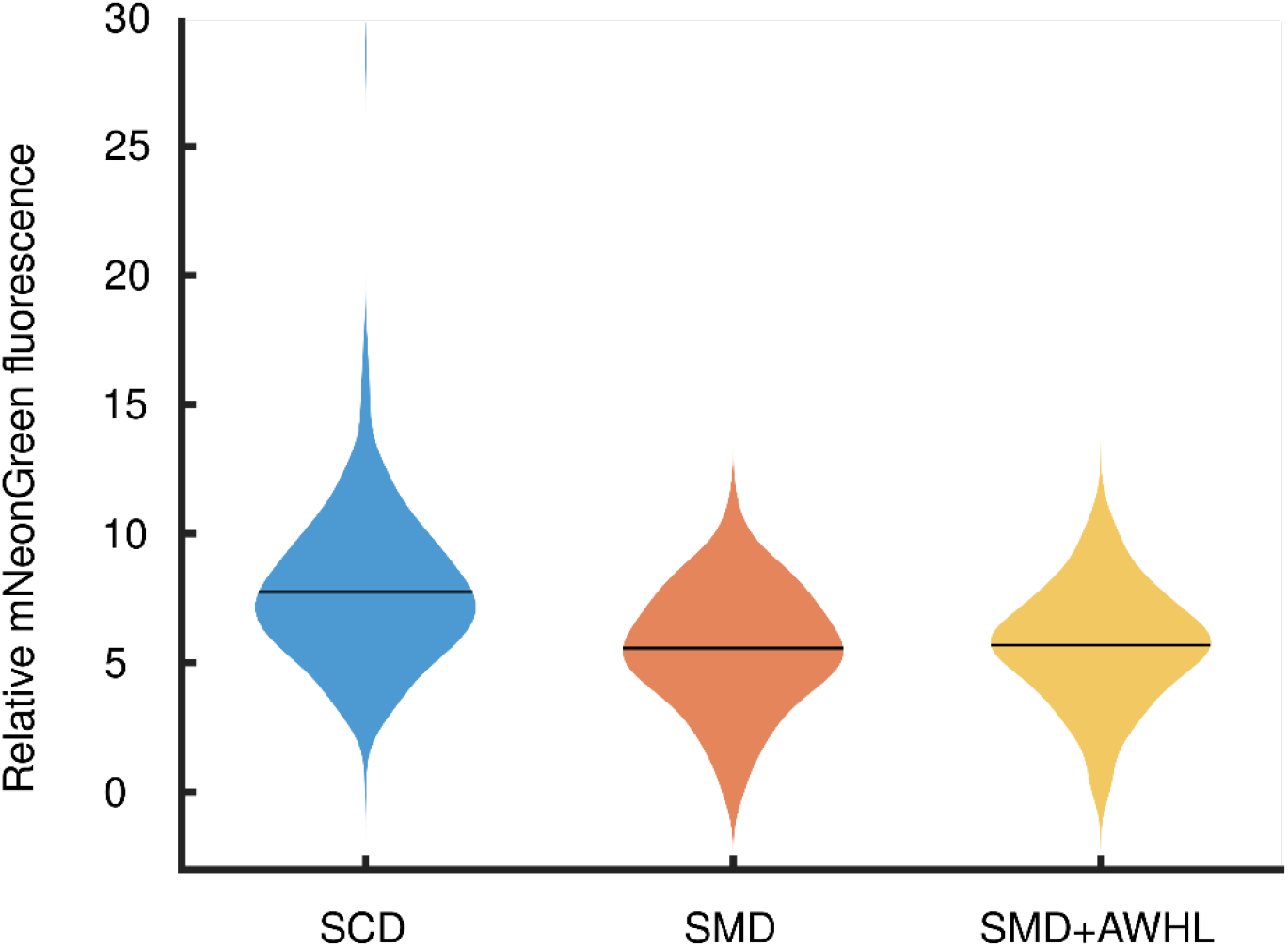
Expression of *ARG3pr-ARG3-mNeonGreen* measured in synthetic complete and in synthetic minimal media. SCM - Synthetic complete media; SMM – Synthetic minimal media; SMM+AWHL – Synthetic minimal media with adenine, tryptophan, histidine and leucine, for which the tested strain was auxotrophic. Fluorescence values are relative to wild-type autofluorescence in the green channel. Numbers of analyzed cells are given in Supplementary Table 17. Black horizontal bars indicate the mean.

**Supplementary Figure 11.**
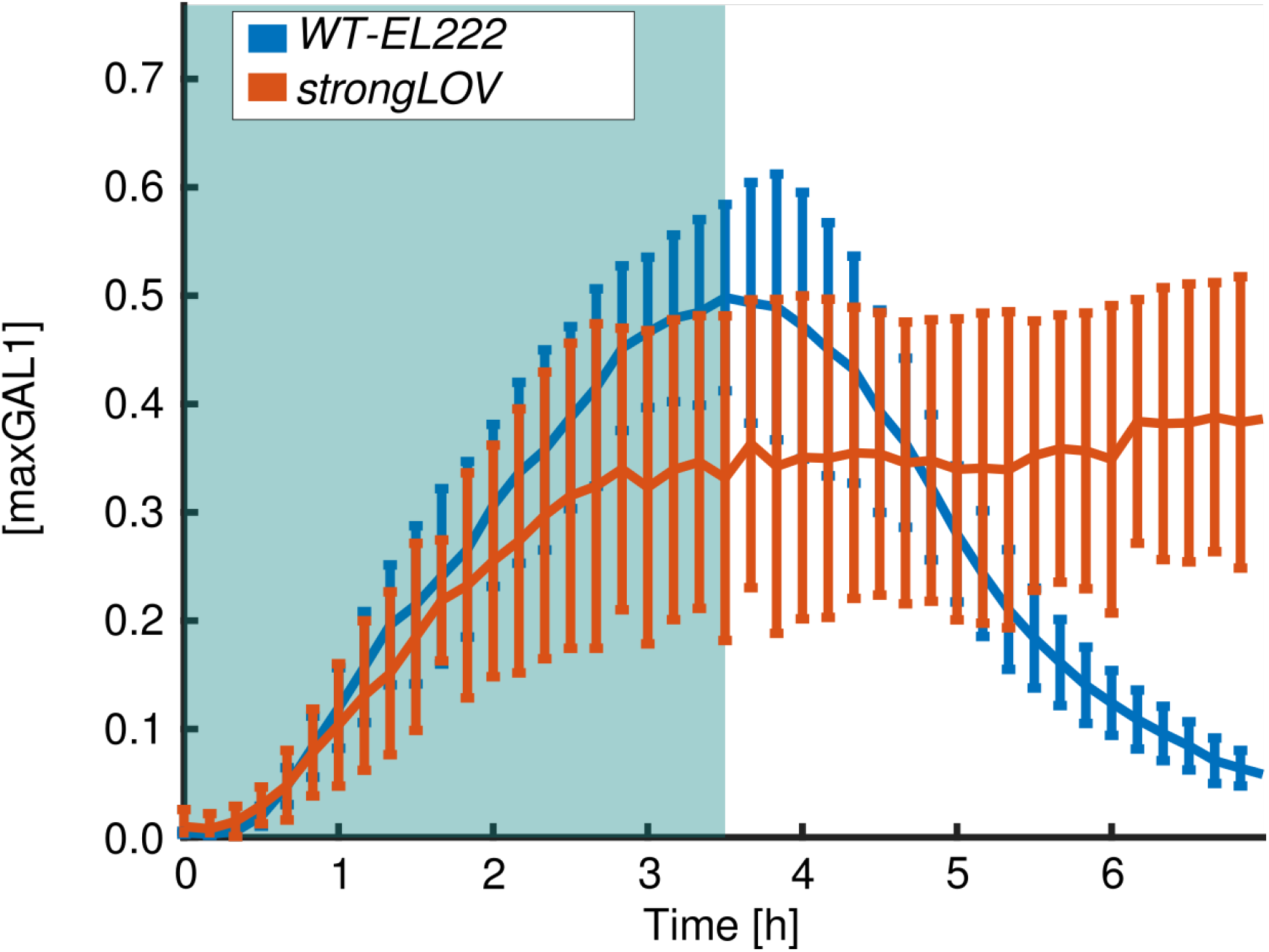
Comparison of wild-type El222 and strongLOV (El222 Glu84Asp) under induction with 80% of the maximal light intensity. Unlike for WT El222, we did not observe the decline in strongLOV after turning off the diascopic illumination used for induction at t = 3.5 h. This could either be due to the long-lasting active state of strongLOV or due to high sensitivity of strongLOV to the light used for exciting the *LIP-yEVenus-PEST* reporter, which partly overlaps with the El222 activation spectrum. To differentiate these possibilities, we performed experiments in liquid culture shown in Fig. 13 B in which we turned off the blue light source for inducing strongLOV-*LIP* and monitored the dynamics of the fluorescent reporter by sampling the population of cells, which were kept in the dark, at various times. We found that when kept in absolute darkness, the strongLOV system switched off with the same kinetics as El222 (Fig. 13 B). Blue background denotes the presence of light. Vertical bars denote the standard deviation.

**Supplementary Table 1:** Plasmids used in the study

| Plasmid | Backbone | Insert | Restriction enzymes used for cloning | Bacterial selection marker | Source |
| --- | --- | --- | --- | --- | --- |
| pVG9 | pCL10 | *pC120 – yEVENus – CLN2 PEST – ADH1t* | *PacI* and *BamHI* | AmpR | *This study* |
| pVG10 | pCL10 | *GALL – yEVENus – CLN2 PEST – ADH1t* | *PacI* and *BamHI* | AmpR | *This study* |
| pVG11 | pCL10 | *[GAL1pr-pC120-GAL1pr] – yEVENus – CLN2 PEST – ADH1t* | *PacI* and *BamHI* | AmpR | *This study* |
| pVG45 | pCL10 | *CUP1pr – yEVENus – CLN2 PEST – ADH1t* | *PacI* and *BamHI* | AmpR | *This study* |
| pVG46 | pCL10 | *PHO5pr – yEVENus – CLN2 PEST – ADH1t* | *BsWI* and *BamHI* | AmpR | *This study* |
| pVG47 | pCL10 | *tetOpr – yEVENus – CLN2 PEST – ADH1t* | *PacI* and *BamHI* | AmpR | *This study* |
| pVG48 | EZL105<sup>6</sup> | *PGK1pr – rtTA – CYC1t* | *NheI* and *XhoI* | AmpR | *This study* |
| pVG49 | pCL10 | *GAL1pr – yEVENus – CLN2 PEST – ADH1t* | *PacI* and *BamHI* | AmpR | *This study* |
| pVG50 | pCL10 | *ADH1t – tetOpr – yEVENus – CLN2 PEST – ADH1t* | *PacI* and *BamHI* | AmpR | *This study* |
| pVG88 | pCL10 | *ARG3pr – yEVENus – CLN2 PEST – ADH1t* | *PacI* and *BamHI* | AmpR | *This study* |
| EZL105 | - | *PGK1pr – EL222 – CYC1t* | - | AmpR | <sup>6</sup> |
| pVG52 | pVG35 (unpublished) | *pC120 – CLB2kd – yEVENus – ADH1t* | *XbaI* and *SapI* | AmpR | *This study* |
| pVG97 | pVG94 (unpublished) | *yEVENus – STOP – URA3 5'UTR* | *KpnI* | AmpR | *This study* |
| pVG106 | pVG48 | *PGK1pr – Z3EV – CYC1t* | *NheI* and *XhoI* | AmpR | *This study* |
| pVG107 | pCL10 | *Z3EVpr – yEVENus – CLN2 PEST – ADH1t* | *BamHI* and *PacI* | AmpR | *This study* |
| pVG108 | pRD14 (unpublished) | *ADH1pr – PhyB – GAL4BD – ADH1t* and *ADH1pr – PIF3 – GAL4AD – ADH1t* | *DraIII* and *AflIII* | AmpR | *This study* |
| pVG109 | pCL10 | *MET3pr – yEVENus – CLN2 PEST – ADH1t* for integration in chromosome I | *KpnI* | AmpR | *This study* |
| pVG122 | EZL105<sup>6</sup> | *PGK1pr – EL222 (Glu84Asp) – CYC1t* | *EcoRI* and *XhoI* | AmpR | *This study* |
| pCL10 | - | *MET3pr – yEVENus – CLN2 PEST – ADH1t* | - | AmpR | *LPBS* |

**Supplementary Table 2:**
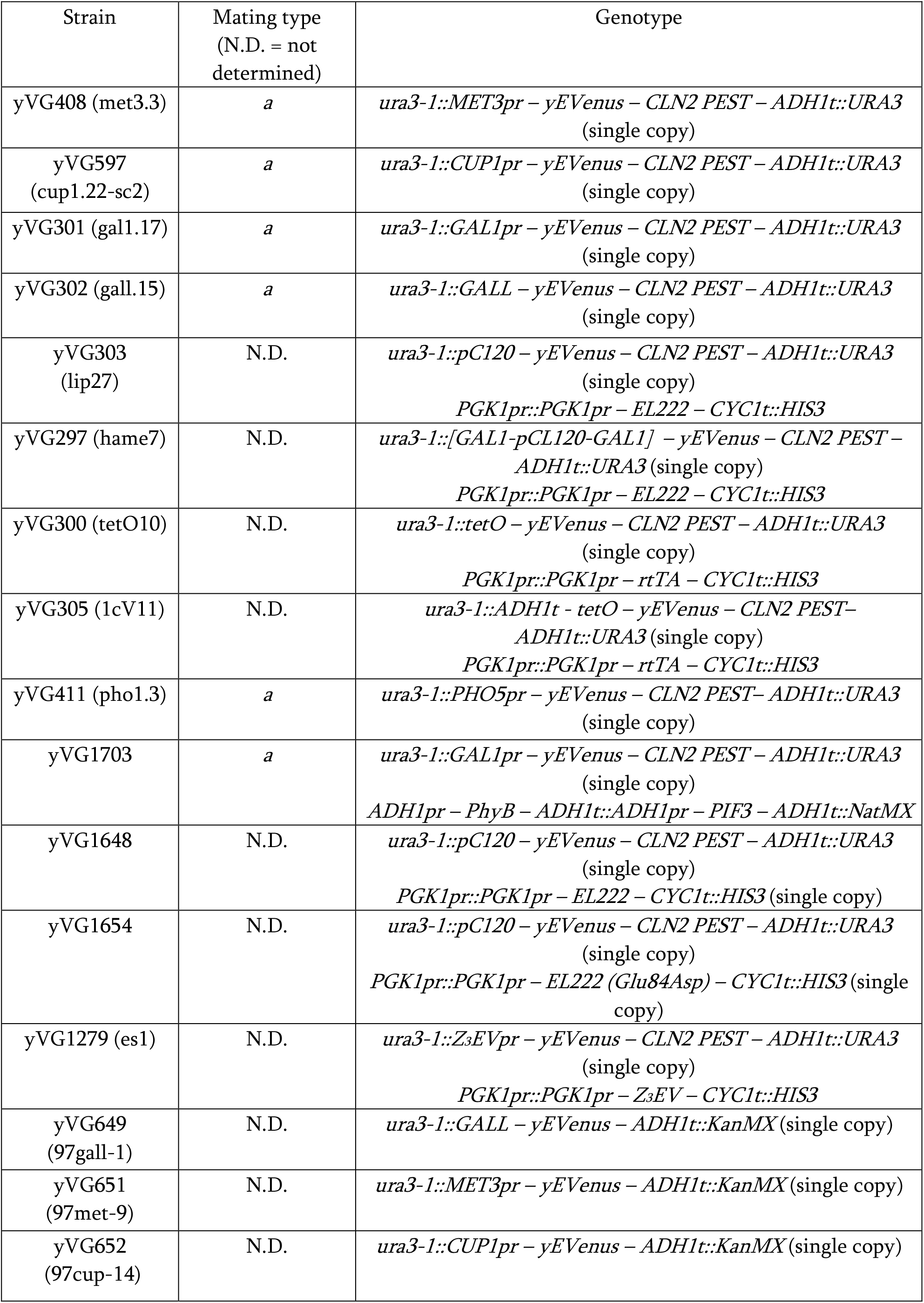
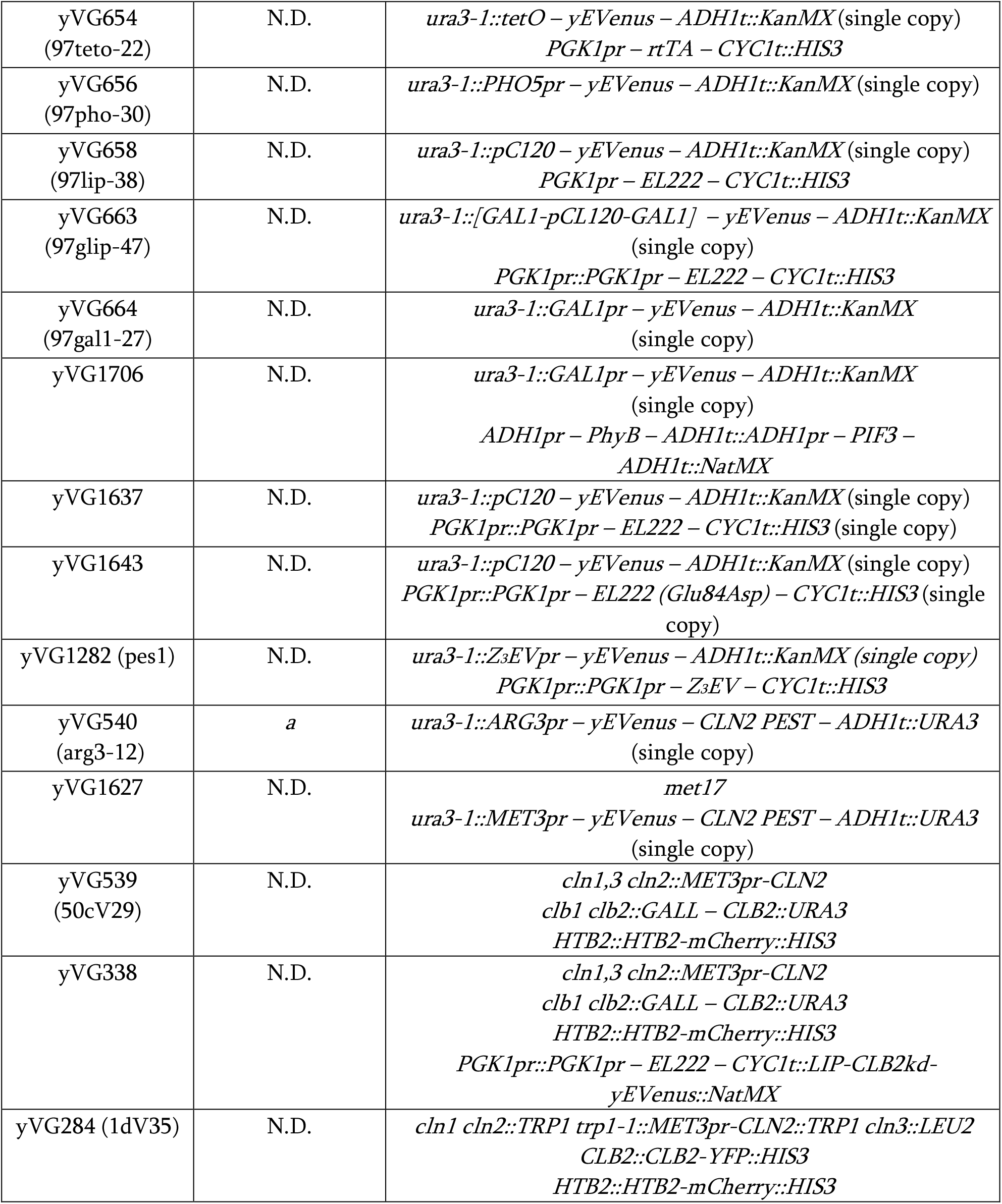
Yeast strains used in the study

Supplementary Tables 3-12. p-values for significance of the differences between all pairs of measurements described in the main text. Red background denotes p < 0.05. In cases where the t-score or z-score, based on which the p values were calculated, was bigger than 50, we approximated the Student distribution by a standard *N*(0,1) distribution. The number of degrees of freedom in these cases was always bigger than 30.

**Supplementary Table 3:** p-values calculated by one-tailed t-test for initial slope (*i*) data, Figure 5 B.

|  | GAL1 | CUP1 | LIP_80p | MET3 | Z3EV | stron... | GLIP | LIP_20p | tetO | PHO5 | PhyB | GALL |
| --- | --- | --- | --- | --- | --- | --- | --- | --- | --- | --- | --- | --- |
| GAL1 |  |  |  |  |  |  |  |  |  |  |  |  |
| CUP1 | 5.56e-02 |  |  |  |  |  |  |  |  |  |  |  |
| LIP_80p | 4.63e-05 | 2.34e-03 |  |  |  |  |  |  |  |  |  |  |
| MET3 | 4.28e-05 | 2.73e-03 | 8.78e-01 |  |  |  |  |  |  |  |  |  |
| Z3EV | 1.19e-05 | 3.35e-04 | 2.22e-01 | 3.75e-01 |  |  |  |  |  |  |  |  |
| strongLOV | 1.97e-06 | 2.34e-05 | 9.67e-03 | 3.23e-02 | 1.34e-01 |  |  |  |  |  |  |  |
| GLIP | 6.38e-09 | 1.96e-10 | 4.58e-14 | 3.58e-11 | 2.22e-11 | 1.23e-06 |  |  |  |  |  |  |
| LIP_20p | 9.13e-10 | 6.01e-12 | 1.44e-16 | 1.05e-13 | 5.60e-14 | 6.81e-09 | 5.87e-02 |  |  |  |  |  |
| tetO | 1.53e-09 | 1.36e-11 | 5.01e-16 | 3.18e-13 | 1.71e-13 | 1.99e-08 | 1.26e-01 | 6.76e-01 |  |  |  |  |
| PHO5 | 2.50e-10 | 3.01e-11 | 1.16e-09 | 3.94e-09 | 2.13e-08 | 2.14e-06 | 9.61e-02 | 7.08e-01 | 5.02e-01 |  |  |  |
| PhyB | 1.60e-11 | 3.32e-13 | 6.11e-12 | 1.66e-11 | 1.02e-10 | 1.20e-08 | 1.78e-03 | 5.21e-02 | 2.55e-02 | 1.83e-01 |  |  |
| GALL | 7.46e-11 | 5.12e-15 | 2.03e-24 | 2.74e-22 | 1.43e-23 | 1.34e-13 | 3.94e-13 | 1.44e-07 | 7.12e-09 | 3.07e-03 | 2.24e-01 |  |

**Supplementary Table 4:**
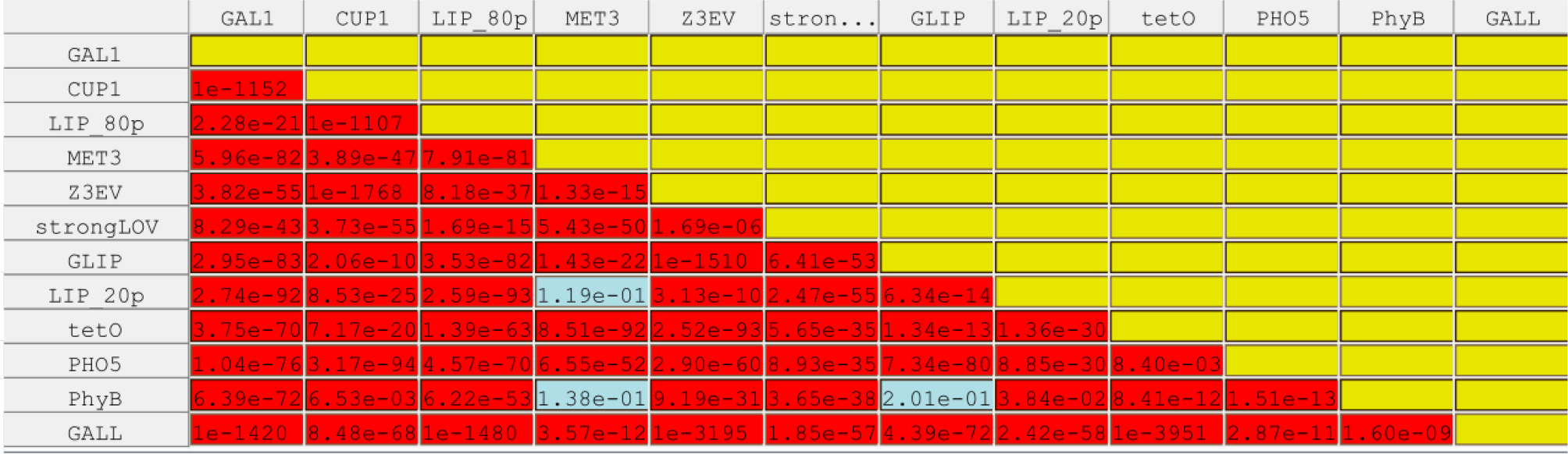
p-values calculated by one-tailed t-test for pairs of maximum fluorescence data, Figure 5 C.

|  | GAL1 | CUP1 | LIP_80p | MET3 | Z3EV | stron... | GLIP | LIP_20p | tetO | PHO5 | PhyB | GALL |
| --- | --- | --- | --- | --- | --- | --- | --- | --- | --- | --- | --- | --- |
| GAL1 |  |  |  |  |  |  |  |  |  |  |  |  |
| CUP1 | 1e-1152 |  |  |  |  |  |  |  |  |  |  |  |
| LIP_80p | 2.28e-21 | 1e-1107 |  |  |  |  |  |  |  |  |  |  |
| MET3 | 5.96e-82 | 3.89e-47 | 7.91e-81 |  |  |  |  |  |  |  |  |  |
| Z3EV | 3.82e-55 | 1e-1768 | 8.18e-37 | 1.33e-15 |  |  |  |  |  |  |  |  |
| strongLOV | 8.29e-43 | 3.73e-55 | 1.69e-15 | 5.43e-50 | 1.69e-06 |  |  |  |  |  |  |  |
| GLIP | 2.95e-83 | 2.06e-10 | 3.53e-82 | 1.43e-22 | 1e-1510 | 6.41e-53 |  |  |  |  |  |  |
| LIP_20p | 2.74e-92 | 8.53e-25 | 2.59e-93 | 1.19e-01 | 3.13e-10 | 2.47e-55 | 6.34e-14 |  |  |  |  |  |
| tetO | 3.75e-70 | 7.17e-20 | 1.39e-63 | 8.51e-92 | 2.52e-93 | 5.65e-35 | 1.34e-13 | 1.36e-30 |  |  |  |  |
| PHO5 | 1.04e-76 | 3.17e-94 | 4.57e-70 | 6.55e-52 | 2.90e-60 | 8.93e-35 | 7.34e-80 | 8.85e-30 | 8.40e-03 |  |  |  |
| PhyB | 6.39e-72 | 6.53e-03 | 6.22e-53 | 1.38e-01 | 9.19e-31 | 3.65e-38 | 2.01e-01 | 3.84e-02 | 8.41e-12 | 1.51e-13 |  |  |
| GALL | 1e-1420 | 8.48e-68 | 1e-1480 | 3.57e-12 | 1e-3195 | 1.85e-57 | 4.39e-72 | 2.42e-58 | 1e-3951 | 2.87e-11 | 1.60e-08 |  |

**Supplementary Table 5:**
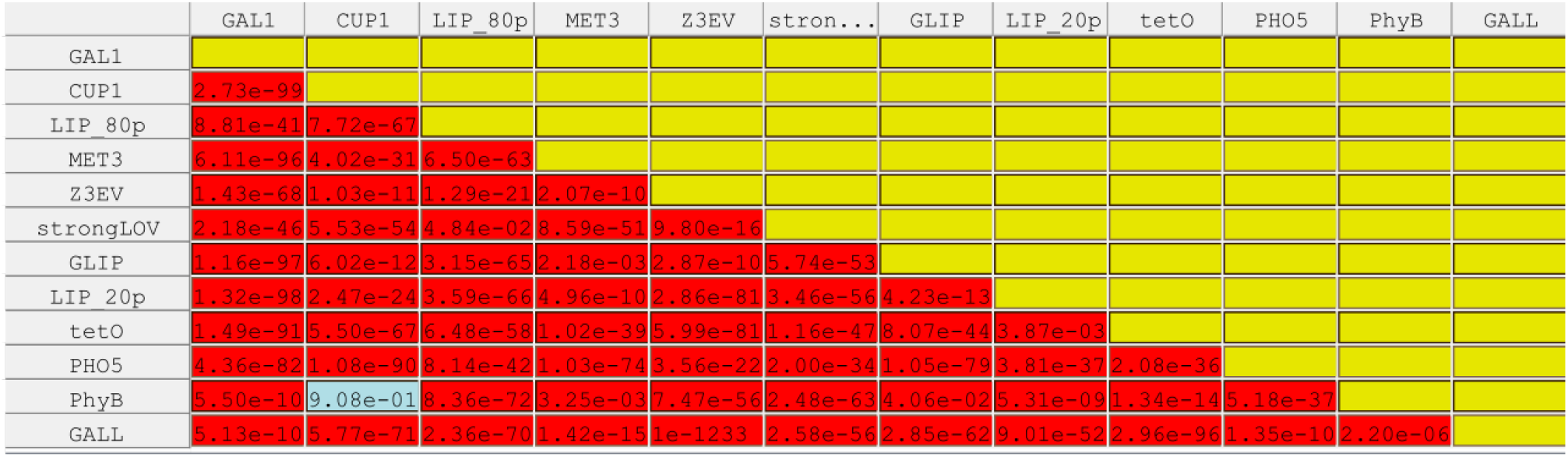
p-values calculated by one-tailed t-test for pairs of steady-state fluorescence data, Figure 5 D.

**Supplementary Table 6:**
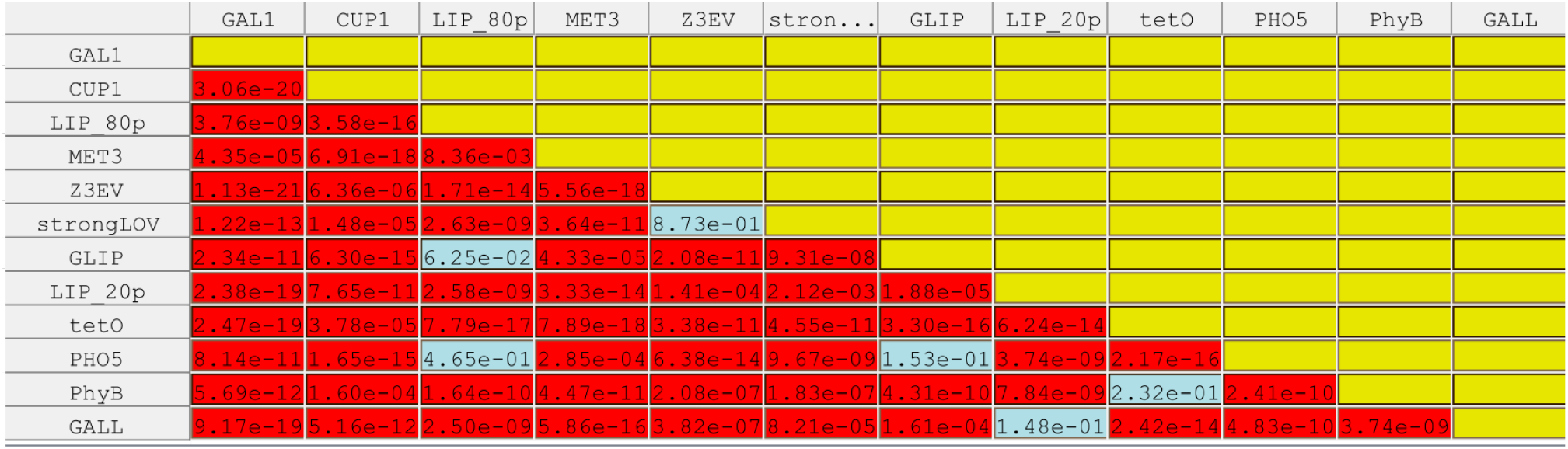
p-values calculated by one-tailed t-test for pairs of basal fluorescence data, Figure 5 E.

**Supplementary Table 7:** p-values calculated by one-tailed t-test for pairs of basal fluorescence parameter (*b*), Figure 5 F.

|  | GAL1 | CUP1 | LIP_80p | MET3 | Z3EV | stron... | GLIP | LIP_20p | tetO | PHO5 | PhyB | GALL |
| --- | --- | --- | --- | --- | --- | --- | --- | --- | --- | --- | --- | --- |
| GAL1 |  |  |  |  |  |  |  |  |  |  |  |  |
| CUP1 | 4.35e-14 |  |  |  |  |  |  |  |  |  |  |  |
| LIP_80p | 7.23e-09 | 7.73e-10 |  |  |  |  |  |  |  |  |  |  |
| MET3 | 1.16e-01 | 8.20e-13 | 1.42e-03 |  |  |  |  |  |  |  |  |  |
| Z3EV | 1.28e-12 | 2.06e-08 | 5.83e-03 | 5.61e-07 |  |  |  |  |  |  |  |  |
| strongLOV | 6.00e-10 | 2.72e-02 | 3.97e-06 | 6.46e-09 | 7.80e-05 |  |  |  |  |  |  |  |
| GLIP | 1.29e-09 | 1.12e-07 | 4.89e-02 | 2.23e-05 | 7.00e-01 | 3.62e-04 |  |  |  |  |  |  |
| LIP_20p | 2.20e-20 | 1.18e-04 | 1.73e-09 | 6.70e-13 | 1.05e-07 | 1.09e-01 | 7.20e-04 |  |  |  |  |  |
| tetO | 5.32e-16 | 6.68e-03 | 7.12e-13 | 1.58e-15 | 1.47e-11 | 5.75e-06 | 1.88e-11 | 9.38e-09 |  |  |  |  |
| PHO5 | 2.65e-11 | 3.06e-09 | 3.24e-01 | 7.71e-05 | 4.41e-02 | 1.40e-05 | 1.71e-01 | 8.45e-09 | 2.51e-12 |  |  |  |
| PhyB | 9.02e-04 | 8.52e-01 | 1.51e-02 | 2.46e-03 | 3.39e-02 | 4.39e-01 | 4.21e-02 | 1.93e-01 | 1.26e-01 | 2.13e-02 |  |  |
| GALL | 7.24e-19 | 6.02e-05 | 5.81e-08 | 1.03e-11 | 5.02e-06 | 6.61e-02 | 3.42e-03 | 6.54e-01 | 4.76e-09 | 3.50e-07 | 1.66e-01 |  |

**Supplementary Table 8:** p-values calculated by one-tailed t-test for pairs of degradation rates (*d*) data, Figure 5 G.

|  | GAL1 | CUP1 | LIP_80p | MET3 | Z3EV | GLIP | LIP_20p | tetO | PHO5 | PhyB | GALL |
| --- | --- | --- | --- | --- | --- | --- | --- | --- | --- | --- | --- |
| GAL1 |  |  |  |  |  |  |  |  |  |  |  |
| CUP1 | 1.79e-55 |  |  |  |  |  |  |  |  |  |  |
| LIP_80p | 2.23e-01 | 1.31e-50 |  |  |  |  |  |  |  |  |  |
| MET3 | 9.20e-51 | 5.69e-13 | 5.89e-55 |  |  |  |  |  |  |  |  |
| Z3EV | 1.12e-12 | 7.42e-76 | 4.58e-11 | 1e-2238 |  |  |  |  |  |  |  |
| GLIP | 4.08e-07 | 2.73e-67 | 1.45e-09 | 4.72e-23 | 4.68e-12 |  |  |  |  |  |  |
| LIP_20p | 5.37e-09 | 1.03e-33 | 2.43e-06 | 1.16e-84 | 4.29e-14 | 3.56e-22 |  |  |  |  |  |
| tetO | 1.90e-61 | 4.11e-01 | 2.98e-56 | 3.20e-18 | 1.41e-12 | 1.37e-67 | 1.90e-45 |  |  |  |  |
| PHO5 | 7.35e-01 | 1.56e-55 | 1.46e-01 | 3.97e-47 | 2.65e-15 | 8.08e-06 | 9.88e-09 | 1.57e-67 |  |  |  |
| PhyB | 2.07e-85 | 1.39e-43 | 9.94e-83 | 1e-1323 | 1.94e-03 | 1.83e-96 | 3.09e-82 | 1.99e-45 | 3.49e-97 |  |  |
| GALL | 4.56e-02 | 6.00e-04 | 9.35e-02 | 1.81e-10 | 4.28e-18 | 5.07e-04 | 7.14e-01 | 9.86e-04 | 3.73e-02 | 1.20e-15 |  |

**Supplementary Table 9:** p-values calculated by one-tailed t-test for pairs of t-on data from Figure 5 H.

|  | GAL1 | CUP1 | LIP_80p | MET3 | Z3EV | stron... | GLIP | LIP_20p | tetO | PHO5 | PhyB | GALL |
| --- | --- | --- | --- | --- | --- | --- | --- | --- | --- | --- | --- | --- |
| GAL1 |  |  |  |  |  |  |  |  |  |  |  |  |
| CUP1 | 1e-3924 |  |  |  |  |  |  |  |  |  |  |  |
| LIP_80p | 1e-5525 | 1.73e-57 |  |  |  |  |  |  |  |  |  |  |
| MET3 | 1.71e-64 | 1e-4345 | 1e-5703 |  |  |  |  |  |  |  |  |  |
| Z3EV | 2.84e-92 | 6.79e-81 | 1e-1194 | 1e-1466 |  |  |  |  |  |  |  |  |
| strongLOV | 1e-2928 | 4.28e-36 | 8.48e-01 | 1e-3710 | 3.79e-97 |  |  |  |  |  |  |  |
| GLIP | 1e-1874 | 6.77e-28 | 4.24e-01 | 1e-2612 | 1.27e-77 | 4.00e-01 |  |  |  |  |  |  |
| LIP_20p | 2.69e-76 | 2.97e-23 | 2.99e-04 | 1e-1450 | 1.47e-55 | 4.28e-04 | 5.74e-03 |  |  |  |  |  |
| tetO | 5.38e-02 | 8.82e-60 | 3.74e-70 | 3.71e-17 | 2.25e-37 | 1.42e-75 | 8.52e-81 | 7.13e-85 |  |  |  |  |
| PHO5 | 3.26e-60 | 1e-1563 | 1e-1896 | 1.64e-48 | 4.90e-77 | 1e-1773 | 1e-1663 | 1e-1430 | 2.64e-66 |  |  |  |
| PhyB | 4.21e-31 | 2.57e-49 | 3.61e-53 | 1.99e-22 | 6.69e-42 | 1.07e-53 | 1.08e-54 | 3.00e-58 | 2.47e-30 | 3.08e-01 |  |  |
| GALL | 1e-4801 | 1.66e-89 | 2.20e-71 | 1e-5555 | 1e-2026 | 6.13e-73 | 2.42e-62 | 6.12e-34 | 1e-1341 | 1e-2564 | 1.52e-62 |  |

**Supplementary Table 10:** p-values calculated by one-tailed t-test for pairs of t-off data, Figure 5 I.

|  | GAL1 | CUP1 | MET3 | Z3EV | tetO | GALL |
| --- | --- | --- | --- | --- | --- | --- |
| GAL1 |  |  |  |  |  |  |
| CUP1 | 7.68e-26 |  |  |  |  |  |
| MET3 | 8.93e-22 | 1.69e-18 |  |  |  |  |
| Z3EV | 1.07e-03 | 8.21e-04 | 1.00e-03 |  |  |  |
| tetO | 8.41e-27 | 1.26e-12 | 9.15e-24 | 5.70e-04 |  |  |
| GALL | 1e-1872 | 1.47e-25 | 1e-1535 | 6.31e-04 | 2.04e-02 |  |

**Supplementary Table 11:** p-values calculated by one-tailed t-test for pairs of leakiness measurements data from Figure 6 A.

|  | WT | GAL1 | CUP | EL222 | MET | Z3EV | stron... | GLIP | TETO | PHO | PhyB_PIF | GALL |
| --- | --- | --- | --- | --- | --- | --- | --- | --- | --- | --- | --- | --- |
| WT |  |  |  |  |  |  |  |  |  |  |  |  |
| GAL1 | 9.12e-07 |  |  |  |  |  |  |  |  |  |  |  |
| CUP | 1e-2689 | 1e-2667 |  |  |  |  |  |  |  |  |  |  |
| EL222 | 1e-2223 | 1e-2124 | 1e-1973 |  |  |  |  |  |  |  |  |  |
| MET | 2.12e-75 | 5.45e-69 | 1e-2232 | 1.48e-73 |  |  |  |  |  |  |  |  |
| Z3EV | 4.15e-60 | 1.95e-58 | 1e-1458 | 1.73e-04 | 3.28e-30 |  |  |  |  |  |  |  |
| strongLOV | 1.03e-10 | 4.90e-10 | 5.64e-25 | 9.93e-82 | 5.92e-94 | 1.86e-76 |  |  |  |  |  |  |
| GLIP | 1.04e-54 | 2.55e-34 | 1e-2628 | 1e-1998 | 8.29e-57 | 4.15e-55 | 1.04e-10 |  |  |  |  |  |
| TETO | 1e-1774 | 1e-1756 | 8.87e-36 | 1e-1183 | 1e-1405 | 4.68e-20 | 2.36e-02 | 1e-1722 |  |  |  |  |
| PHO | 1.94e-24 | 2.67e-21 | 1e-2183 | 6.11e-55 | 9.69e-04 | 9.06e-36 | 3.25e-97 | 1.24e-15 | 1e-1383 |  |  |  |
| PhyB_PIF | 6.95e-12 | 1.40e-11 | 2.53e-54 | 1.49e-03 | 2.79e-07 | 2.03e-02 | 2.45e-27 | 5.40e-11 | 1.55e-35 | 2.40e-08 |  |  |
| GALL | 2.62e-13 | 3.98e-12 | 1e-2057 | 4.63e-24 | 4.50e-12 | 3.58e-16 | 1.14e-88 | 1.18e-11 | 1e-1266 | 9.95e-18 | 1.85e-05 |  |

**Supplementary Table 12:** p-values calculated by one-tailed t-test for pairs of the leakiness data from Figure 6 B.

|  | WT_D | GAL1_D | GAL1_R | GALL... | GALL... | GLIP_D | GLIP_R | GLIP_RG | PhyB_R | PhyB... | PhyB... |
| --- | --- | --- | --- | --- | --- | --- | --- | --- | --- | --- | --- |
| WT_D |  |  |  |  |  |  |  |  |  |  |  |
| GAL1_D | 5.88e-05 |  |  |  |  |  |  |  |  |  |  |
| GAL1_R | 9.12e-07 | 3.81e-01 |  |  |  |  |  |  |  |  |  |
| GALL_in_D | 1.11e-16 | 3.12e-05 | 1.08e-03 |  |  |  |  |  |  |  |  |
| GALL_in_R | 2.62e-13 | 1.72e-12 | 3.98e-12 | 1.21e-12 |  |  |  |  |  |  |  |
| GLIP_D | 1.04e-54 | 1.07e-36 | 2.55e-34 | 1.26e-23 | 1.18e-11 |  |  |  |  |  |  |
| GLIP_R | 1.61e-18 | 7.89e-18 | 1.78e-18 | 2.83e-18 | 5.76e-13 | 1.82e-17 |  |  |  |  |  |
| GLIP_RG | 1e-2233 | 1e-2203 | 1e-2198 | 1e-2181 | 1e-1145 | 1e-2134 | 3.26e-02 |  |  |  |  |
| PhyB_R | 1.37e-06 | 6.33e-06 | 8.63e-06 | 2.37e-05 | 1.75e-06 | 2.28e-04 | 1.02e-10 | 1.79e-10 |  |  |  |
| PhyB_R_PCB | 6.95e-12 | 1.24e-11 | 1.40e-11 | 2.10e-11 | 1.85e-05 | 5.40e-11 | 5.42e-09 | 1.92e-11 | 4.11e-08 |  |  |
| PhyB_D_PCB | 5.75e-18 | 1.58e-17 | 1.95e-17 | 3.93e-17 | 1.57e-06 | 2.05e-16 | 9.74e-23 | 3.38e-28 | 9.92e-11 | 2.72e-01 |  |

**Supplementary Table 13:**
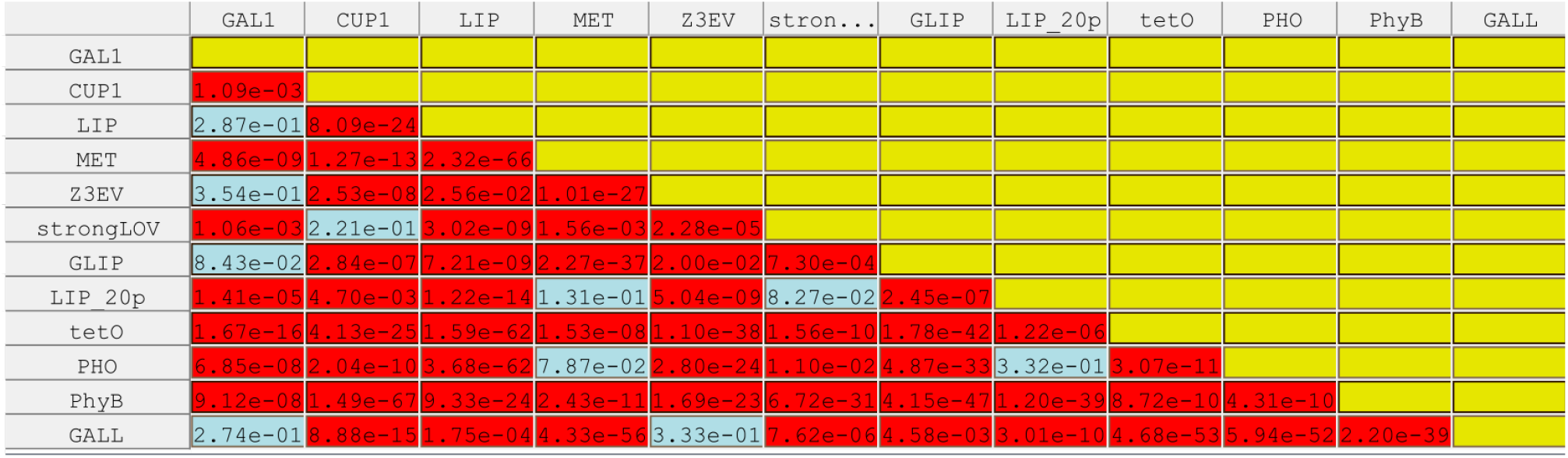
p-values calculated by one-tailed z-test for pairs of area doubling time data from Figure 7.

Supplementary Table 14-16: Numbers of cells used in the experiments

**Supplementary Table 14:** Number of cells used in experiments shown in Figures 2, 5, 6, 7, 8, 9 and 13.

| Inducible system | Number of cells<br>at time t = 0 h | Number of cells<br>at time t = 3.5 h | Number of cells<br>present at the<br>time of shut-off | Relevant figures |
| --- | --- | --- | --- | --- |
| *GAL1pr* | 30 | 129 |  | 2, 5, 6, 7, 8, 9 |
| *LIP* | 35 | 110 |  | 2, 5, 6, 7, 8, 9 |
| *PHO5pr* | 46 | 232 |  | 2, 5, 6, 7, 8, 9 |
| *t-tetOpr* | 92 | 534 |  | 2, 5, 6, 7, 8, 9 |
| *tetOpr* | 66 | 366 |  | 2, 5, 6, 7, 8, 9 |
| *MET3pr* | 84 | 306 |  | 2, 5, 6, 7, 8, 9 |
| *GLIP* | 37 | 135 |  | 2, 5, 6, 7, 8, 9 |
| *CUP1pr* | 62 | 229 |  | 2, 5, 6, 7, 8, 9 |
| *GALL* | 73 | 177 |  | 2, 5, 6, 7, 8, 9 |
| Z<sub>3</sub>EV | 66 | 190 |  | 2, 5, 6, 7, 8, 9 |
| PhyB-PIF3 | 37 | 81 |  | 2, 5, 6, 7, 8, 9 |
| wt-El222-*LIP*<br>20% induction | 70 | 300 |  | 13 |
| strongLOV-*LIP*<br>20% induction | 22 | 78 |  | 13 |
| *GAL1pr* shut-off |  |  | 185 | 5 |
| *LIP* shut-off |  |  | 113 | 5 |
| *tetOpr* shut-off |  |  | 103 | 5 |

**Supplementary Table 15:** Number of cells in experiments shown in Figure 3.

| Construct | Number<br>of cells<br>at t = 0 | Number of<br>cells at<br>time t = 3.5<br>h | Relevant figures |
| --- | --- | --- | --- |
| *MET3pr-yEVenus*<br>integrated at chrV in<br>*MET17-WT* strain | 24 | 73 | 3A, B |
| *MET3pr-yEVenus*<br>integrated at chrI in<br>*MET17-WT* strain | 22 | 60 | 3A |
| *MET3pr-yEVenus*<br>integrated at chrV in<br>*met17Δ* strain | 27 | 39 | 3B |

**Supplementary Table 16:**
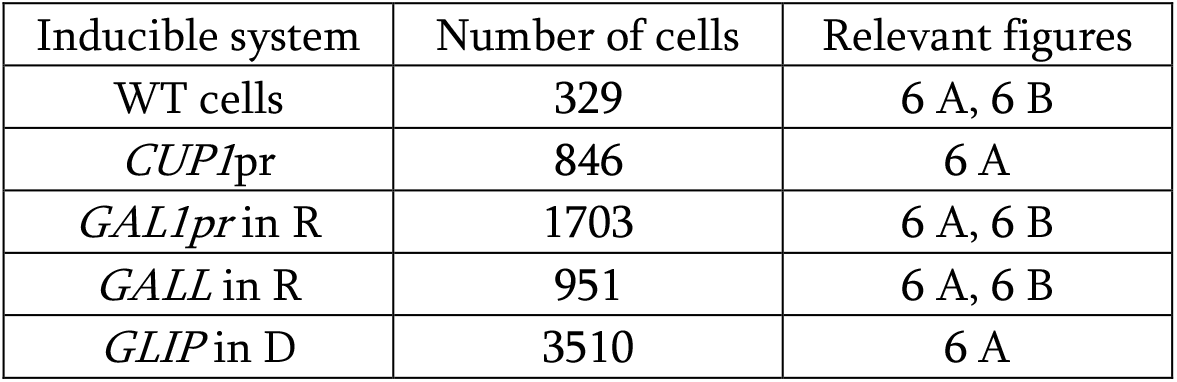
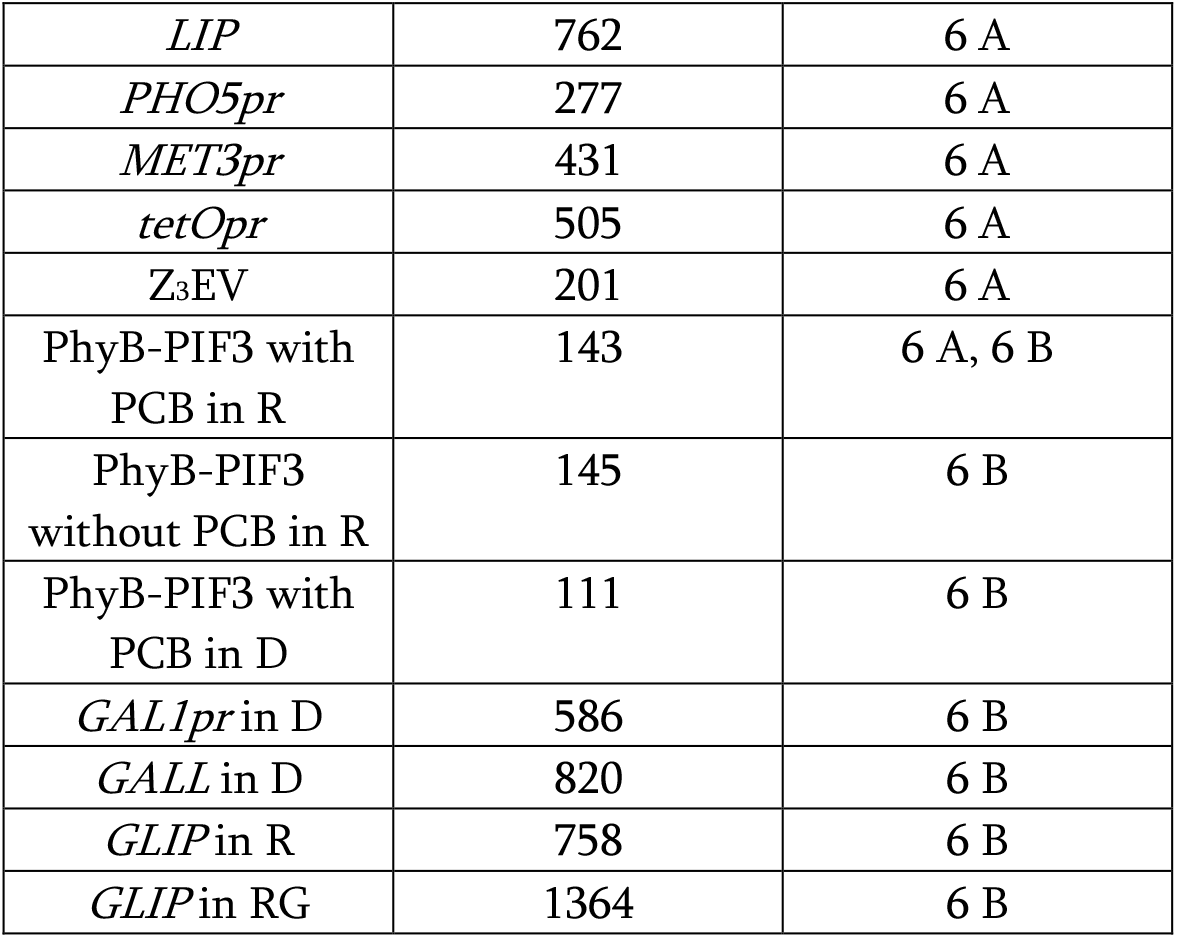
Number of cells used in experiments shown in Figure 6.

**Supplementary Table 17:** Number of cells used in experiments shown in Figure 10.

| Genetic construct/condition | Number of cells | Relevant figures |
| --- | --- | --- |
| *ARG3pr-yEVENUS* in SCD+Met | 321 | 10 |
| *ARG3pr-yEVENUS* in SMM | 69 | 10 |
| *ARG3pr-yEVENUS* in SMM +<br>essential nutrients | 120 | 10 |
| *ARG3pr-ARG3-mNeonGreen* in<br>SCD+Met | 560 | Supplementary Figure 10 |
| *ARG3pr-ARG3-mNeonGreen* in<br>SMM | 179 | Supplementary Figure 10 |
| *ARG3pr-ARG3-mNeonGreen* in<br>SMM + essential nutrients | 677 | Supplementary Figure 10 |

**Supplementary Table 18:** Number of cells used in experiments shown in Figure 15.

| Strain | Experiment | Number of scored<br>cells | Relevant figures |
| --- | --- | --- | --- |
| clnΔ* | 20% light, 0 min delay | 124 | 15 B |
| clnΔ* clb1,2Δ* LIP-CLB2kd | 20% light, 0 min delay | 47 | 15 B |
| clnΔ* | 80% light, 0 min delay | 137 | 15 C |
| clnΔ* clb1,2Δ* LIP-CLB2kd | 80% light, 0 min delay | 86 | 15 C |
| clnΔ* clb1,2Δ* LIP-CLB2kd | 80% light, 20 min delay | 99 | 15 C |
| clnΔ* clb1,2Δ* | N/A | 114 | 15 B, 15 C |

### Supplementary Note 1: Environments used for system induction and deactivation

#### Media used for induction experiments

Standard synthetic complete media without methionine (SC-Met)^126^ was used as the basis for other media, with modifications specific for each system detailed below. We used 2% w/v glucose (D), 3% w/v raffinose (R), or 3% w/v glucose (G).

#### Media for MET3pr induction experiments

Non-inducing: SCD+10x Met (1x Met = 0.02g/mL). Inducing condition: SCD-Met.

#### Media for CUP1pr induction experiments

Non-inducing condition: To make SCD-Met-Cu^2+^, we used yeast nitrogen base without copper (Formedium, UK). Inducing condition: SCD-Met-Cu^2+^ with CuSO_4_ added (0.3 mM).

#### Media for PHO5pr induction experiments

Non-inducing condition: SCD-Met. Inducing condition: To make SCD-Met-Pi (Pi – inorganic phosphate) we used yeast nitrogen base without ammonium-sulfate, without phosphates and without sodium-chloride (MP Biomedicals 4027-812).

#### Media for GAL1pr and GALL induction experiments

Non-inducing condition: SCR-Met. Inducing condition: SCRG-Met (1x raffinose and 1x galactose).

#### Media for tetOpr and t-tetOpr induction experiments

Non-inducing condition: SCD-Met. Inducing condition: SCD-Met with doxycycline added (10 µM).

#### Media for ARG3pr induction experiments

Non-inducing condition: SDC-Met+10xArg (1x Arg = 0.02 g/L of L-arginine monohydrochloride). Inducing condition: SCD-Met-Arg.

For experiments with *ARG3pr*, we also used synthetic minimal media (SMM), containing yeast nitrogen base without all amino acids and without ammonium sulfate, sodium hydroxide, succinic acid, and glucose; as well as SMM+AWHL – Synthetic minimal media with adenine, tryptophan, histidine and leucine, for which the strain we used was auxotrophic.

#### Media for Z3EVpr induction experiments

Non-inducing condition: SCD-Met. Inducing condition: SCD-Met+0.5 µM β-estradiol (diluted from 100x ethanol solution; kept in glass container).

#### Light conditions for the El222-LIP, El222-GLIP, and strongLOV-LIP system

We used the diascopic LED light source of the Nikon Ti2-E microscope for induction. To tune the strength of the inducer, we scaled the level of the input white light to 20%, 40%, or 80% of the maximal intensity, depending on the experiment presented in the manuscript. At maximal strength, the diascopic light produces a beam of white light with 19.60 mW power distributed over a planar circle area with diameter 8.5 mm (average light intensity of 443.67 W/m^2^), as measured by an optical power meter (Thorlabs, US) equipped with an ND2 filter and a S120C sensor (Thorlabs, US) set to a wavelength of 447 nm.

#### Light conditions for the PhyB-PIF3 system

Unless otherwise stated, cells in which PhyB-PIF3 was induced were incubated with PCB for at least 2 h (final concentration of 31.25 µM, diluted from 100x DMSO stock) in the SCR media in darkness. Manipulations during the pre-induction period were performed under green light which does not cause the degradation of PCB. Non-inducing condition: 16 far-red LEDs with a radiation power of 200 mW each and 750 nm emission peak (Roithner LaserTechnik, Austria) assembled on a breadboard and placed above the cell microfluidic chamber at a ≈ 5 cm distance. Inducing condition: 16 red LEDs with a luminosity of 2500 mcd each and 648 nm emission peak (Mouser Electronics, US) assembled on a breadboard and placed above the cell microfluidic chamber at a ≈ 5 cm distance.

### Supplementary Note 2: Measuring yEVenus maturation rate

To limit the space of model parameters to fit to experimental data, we measured the yEVenus protein maturation rate, *f*, directly under our experimental conditions. We blocked protein translation using cycloheximide in cells in which the fluorescent protein is expressed briefly, as in ref.^43^. Specifically, after growing cells in non-inducing media, we turned on *MET3pr-yEVenus* (without the *PEST* sequence) for 30 min, and then turned it off while at the same administering cycloheximide (at a final concentration of 20 ug/mL) which blocks protein synthesis. For an accurate estimation of the maturation rate, we used frequent imaging (every 3 min) but of only a small number of large colonies to avoid photobleaching.

In this experiment, fluorescence levels increase upon brief promoter induction and remain stable after fluorescent protein maturation (Supplementary Fig. 12 A). Since cells stop growing due to the translational block and fluorescent proteins have a half-life of several hours in these cells^127^, the maturation of the already translated fluorescent protein is the only process affecting fluorescence levels. With respect to the model presented in Fig. 5 A, this means that *d* is approximately zero, which leaves *f* as the only parameter influencing the fluorescence. We thus estimated *f*, the maturation rate of yEVenus, by fitting the observed single cell fluorescence levels to:

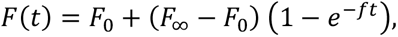

where *F*(*t*) is the level of fluorescence over time, *F_o_* is the level in the beginning of the experiment and *F*_∞_ is the final level. Since there is a lag in the induction of *MET3pr-yEVenus* with respect to the media change, to estimate the maturation rate accurately, we used the timepoints during which the fluorescence level averaged across the population was rising from 5% of (*F*_∞_ − *F*_0_) to 95% of (*F*_∞_ − *F*_0_). After fitting the expression levels on single-cell data (N = 34) to the linearized equation for maturation dynamics 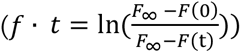, we obtained the mean maturation half-life, 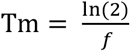, of (16.74 ± 0.69) min (mean ± s.e.m.) (Supplementary Fig. 12 B), a value which is in agreement with previous measurements of Venus maturation dynamics *in vivo*^30,32,43^.

**Supplementary Figure 12.**
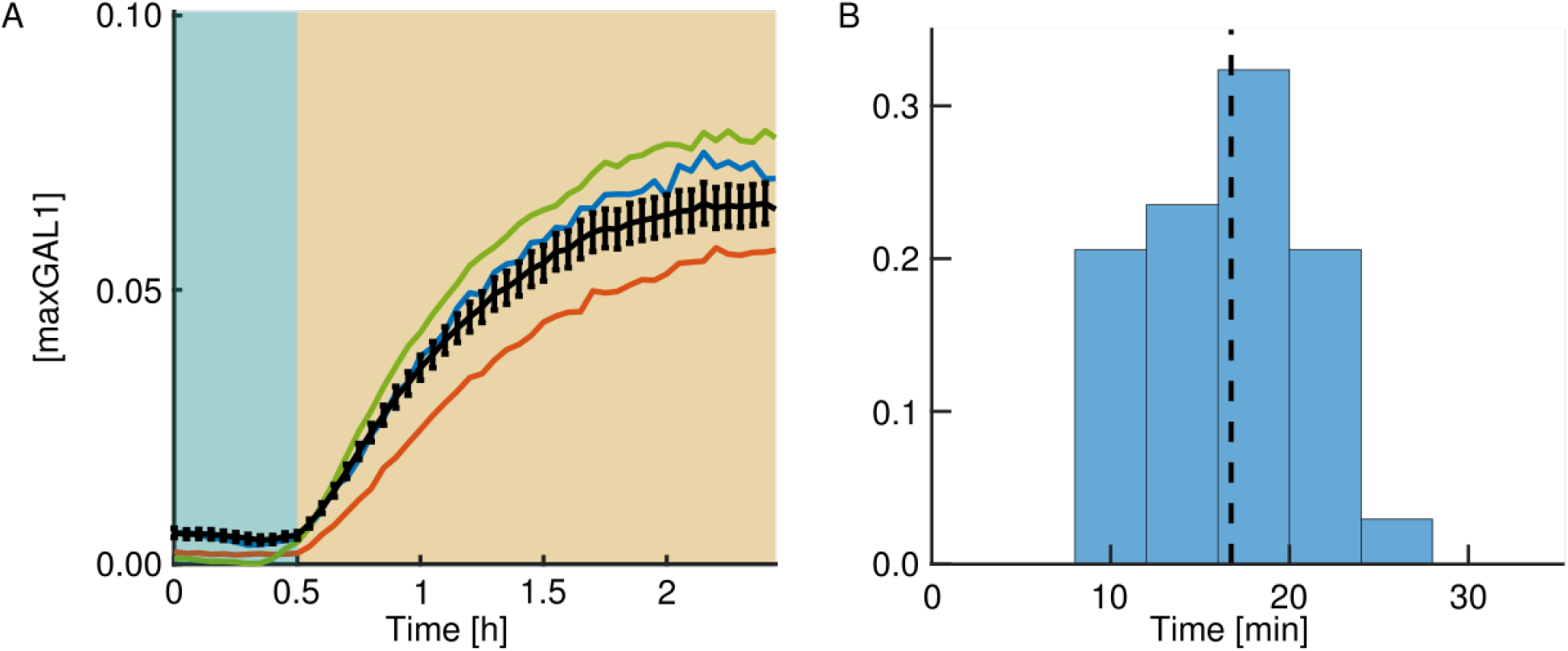
Estimation of yEVenus maturation rate in budding yeast using a translational block A: Cells with the *MET3pr-yEVenus* construct were grown in methionine-rich medium (t < 0 h), then exposed to a brief pulse of no-methionine medium from t = 0 h to t = 0.5 h, which induces the circuit (blue background). After this, cycloheximide was added (yellow background). Black line denotes the average fluorescence level over time (standard errors of the mean (SEM) shown). Colored lines show representative single-cell time courses. B: Histogram of estimated yEVenus maturation half-lives from the single-cell data. Dashed line shows the mean maturation half-life, Tm = 16.74 min.

### Supplementary Note 3: Measuring half-life of yEVenus-PEST fusion protein

The fit in Fig. 8 suggests that active degradation mediated by the PEST degron and dilution due to growth in glucose media contribute about equally to the degradation-and-dilution parameter *d* with half-lives of around 90 min each. However, previous work on the PEST degron also used the last 178 amino acids from the Cln2 protein’s C-terminus and showed that the half-life of yEGfp3 fused to PEST is between 20 and 30 min.^61^ This value was determined by observing fluorescence decay after a cycloheximide block in a *S150-2B* budding yeast strain grown in YPD medium, and was validated by western blot quantification. To verify the degradation rate we obtained from the model fit under our experimental conditions, we measured the half-life of *yEVenus-PEST* in our *W303* cells directly.

We performed a time-course experiment in which we monitored the decay of fluorescence after a cycloheximide translational block. Cells with the *MET3pr-yEVenus-PEST* construct were initially grown in conditions that induce the circuit. Then, we either added methionine to shut off *yEVenus-PEST* expression in control cells or methionine and cycloheximide to additionally shut off translation. By fitting a linear regression to the log of the fluorescence values at timepoints after which cycloheximide takes effect as judged by the abrupt decline of growth, we estimated the growth rate of the cells and the decay rate of yEVenus-PEST for both experimental conditions (Supplementary Fig. 13). The extracted growth doubling time of cells without cycloheximide is T*_g1/2_* = 89.71 min (95% confidence interval: 88.89 min – 90.56 min) while the degradation-and-dilution half-life of yEVenus-PEST is T*_d1/2_* = 43.62 min (95% confidence interval: 43.62 min – 45.59 min). As expected, cells effectively stop growing in cycloheximide, and we measured the growth doubling time to be T*_g1/2_* = 24.83 h (95% confidence interval: 22.98 h – 27.00 h), and is reflected in the larger protein degradation-and-dilution half-life of T*_d1/2_* = 86.10 min (95% confidence interval: 84.40 min – 87.86 min). The differences between the degradation rates and the growth rates give the half-lives for the component of the decay which is due to active degradation mediated by the PEST degron. These values are T*_PEST_* = 91.37 min in the case where cycloheximide is present, and T*_PEST_* = 88.62 min for cells in rich media with methionine.

These values are in agreement with the results in Fig. 8, suggesting that PEST indeed destabilized the yEVenus-PEST in our *W303* cells less compared to yEGfp3-PEST in the *S150-2B* background in previous work^61^ Moreover, the overall degradation-and-dilution half-life of yEVenus-PEST expressed from *MET3pr* in the *W303* genetic background and in synthetic complete media with methionine studied elsewhere was around 39 min^43^ which is in agreement with 43 min we observe. Thus, the differences in the PEST degradation rate can be due to differences in media or, more plausibly, genetic backgrounds of the strains, or specifically differences in the *PEST* sequence encoding the last 178 amino acids (exact *PEST* sequence in S150-2B budding yeast strain was not available from the research article^61^ nor from yeastgenome.org).

**Supplementary Figure 13.**
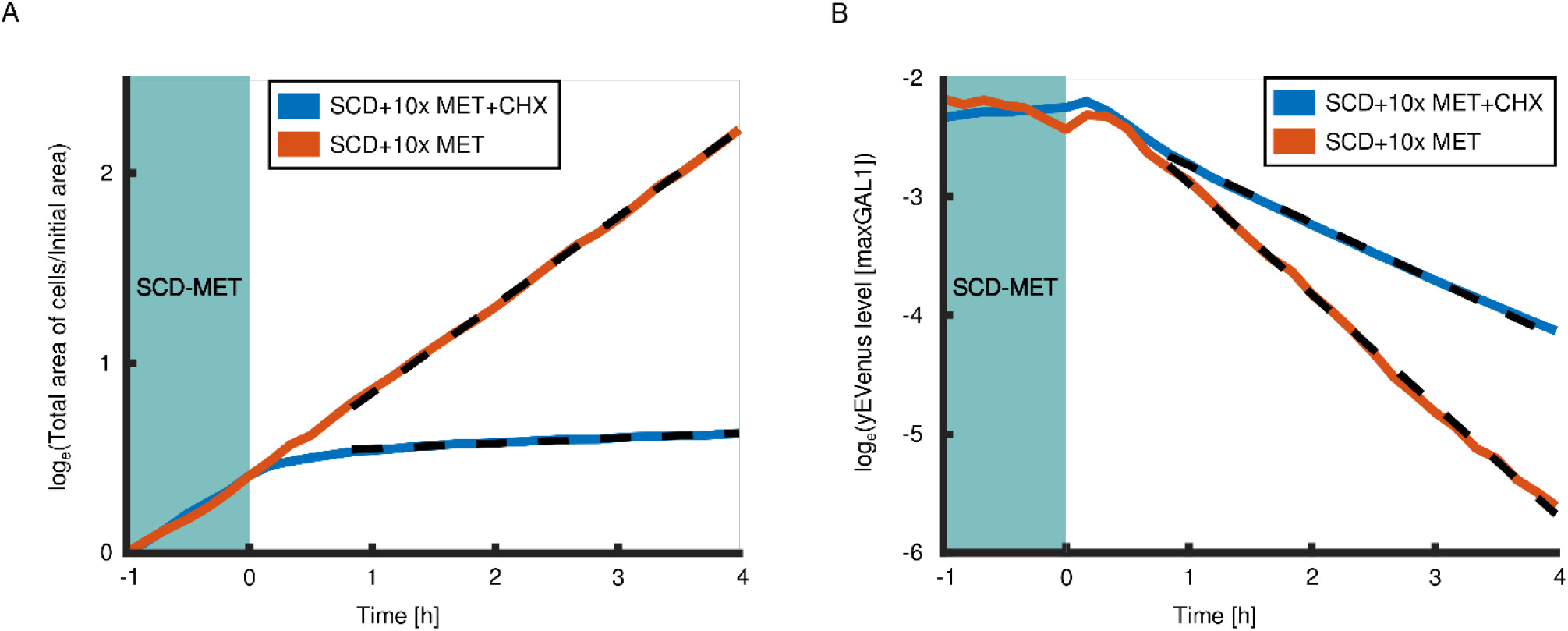
Measuring the half-life of yEVenus-PEST using a translational block. We exposed cells expressing *MET3pr-yEVenus-PEST* to cycloheximide (CHX, c = 20 µg/mL) and monitored yellow fluorescence (orange curve, panel A) and growth (orange curve, panel B). At the same time, we administered only methionine (MET) at 10x concentration, which turns off the genetic circuit, to another group of cells (blue curve in panels A and B). Prior to exposure to different media (t < 0 h), both groups of cells were grown in synthetic complete media lacking methionine (SCD-MET, blue background in panels A and B). Due to faster growth and dilution, yellow fluorescence averaged over cell area decayed faster in cells without cycloheximide. To extract the growth and decay rates, we fit linear functions to the log of the fluorescence values (dashed lines close to orange and blue curves in panels A and B). To subtract the delay with which cycloheximide shuts down translation, we used the timepoints from t = 40 min to t = 4 h for the fit. By finding the differences in the overall degradation rate and the dilution rate, we determined the half-life of yEVenus-PEST for both groups of cells (values in the main text). The number of cells at t = 0 h was 163 for the group of cells treated with cycloheximide and 26 for the group of cells grown in SCD+10xMET, while at t = 4 h these values were 170 and 160, respectively. In panel A, the total area of cells was scaled by the initial area, hence starting at zero after the logarithm was applied.

### Supplementary Note 4: Single-copy integration search procedure

To verify that cells have only one copy of the *promoter-yEVenus-PEST* reporter, we devised a PCR-based procedure that allowed us to distinguish between single and multiple copy insertions. For this, we designed two pairs of primers: p fwd /p rev (p - plasmid) and g fwd /g rev (g - genome) (sequences given in Supplementary Table 19). Both pairs of primers amplify the region containing the *URA3* gene with the difference that the p primers anneal to the plasmid backbone only, while the g primers anneal to the yeast genome only. Since the plasmids were cut inside the plasmid’s *URA3* gene for transformation and insertion, in case of single copy integrations, the p pair of primers should not give a PCR amplicon (Supplementary Figure 14). On the other hand, if the plasmid is integrated in the genome in multiple copies, the p pair of primers will produce an amplicon. With this test, we screened for colonies that showed no PCR product with the p primer pair. To be certain that the lack of amplification was not due to low DNA quality or problems with the PCR reaction, we also performed PCR using the g fwd/p rev and p fwd/g rev pairs of primers, which should show amplification of the DNA regardless of the copy numbers of the reporters. We then only used the strains that showed amplification with g fwd/p rev and p fwd/g rev and no amplification with the p fwd/p rev pair of primers. This also confirmed that the construct is integrated in the *URA3* locus. We repeated this analysis at least twice with PCR reactions performed on independent genomic DNA extractions.

To perform an additional check that strains contained only one copy of the genetic circuit, we designed our *PEST* removal strategy so that strains that do not contain the *PEST* sequence become uracil auxotrophs only in case there is a single copy of *URA3* in the genome. After the transformation with the *KanMX*-marked *PEST*-removal plasmid, strains were dead on plates lacking uracil, confirming again that the *promoter-yEVenus-PEST* construct was present as a single copy.

**Supplementary Figure 14.**
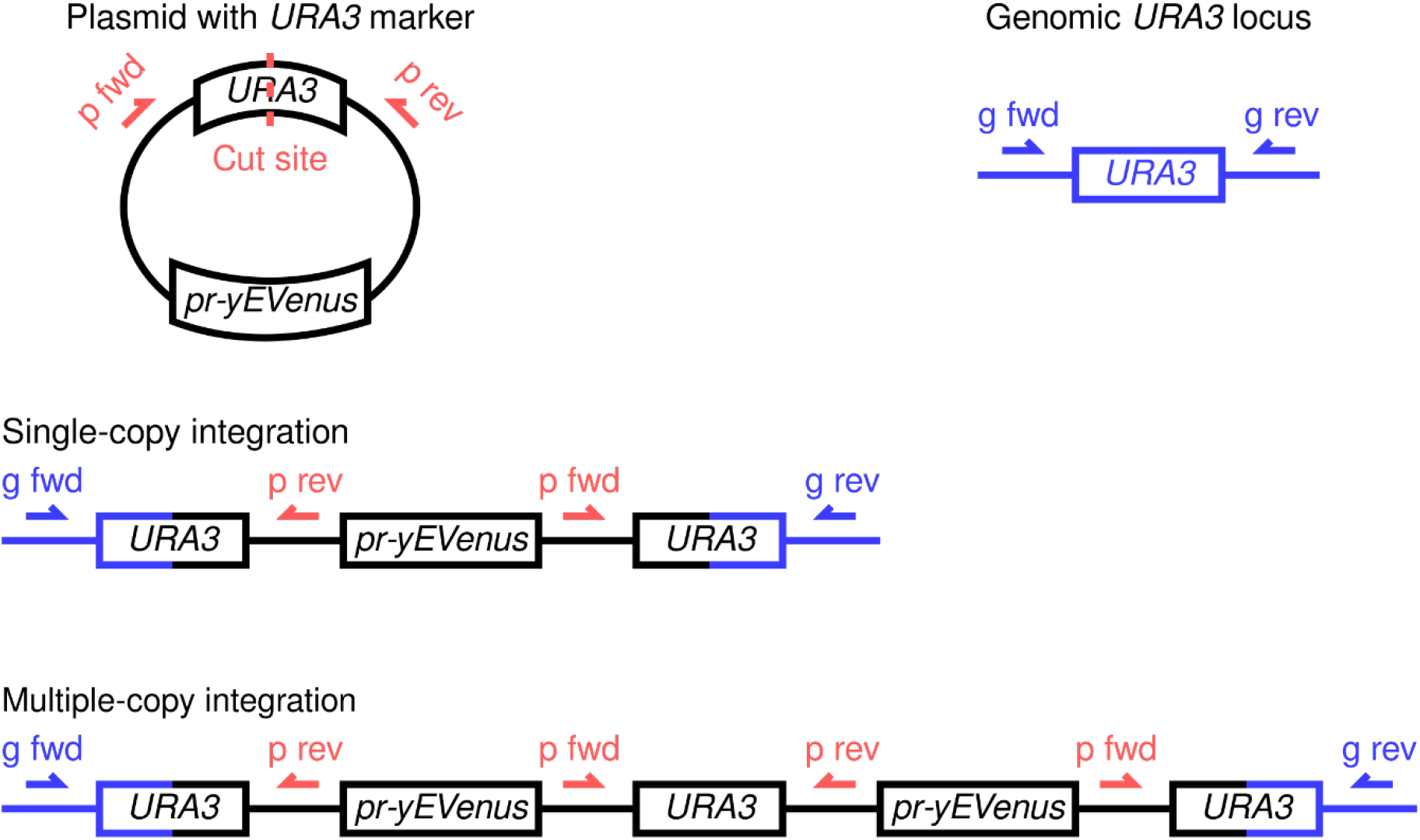
Single and multiple-copy integrations can be distinguished by a PCR-based strategy.

**Supplementary Table 19.**
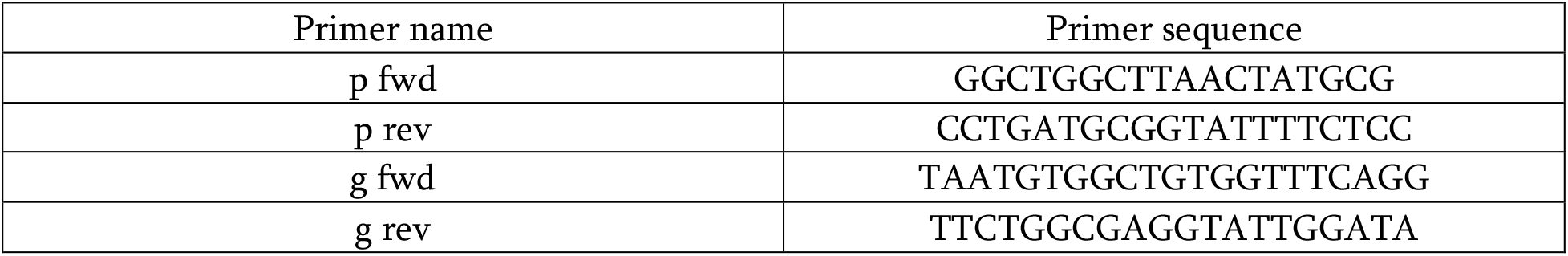
Primer sequences used for checking single-copy integrations

### Supplementary Note 5: DNA sequences of the promoters

Promotors were cloned between BamHI and PacI restriction sites, unless otherwise specified.

pVG9: *LIP* (5 El222 binding sites, also known as pCL120 + minimal promoter)

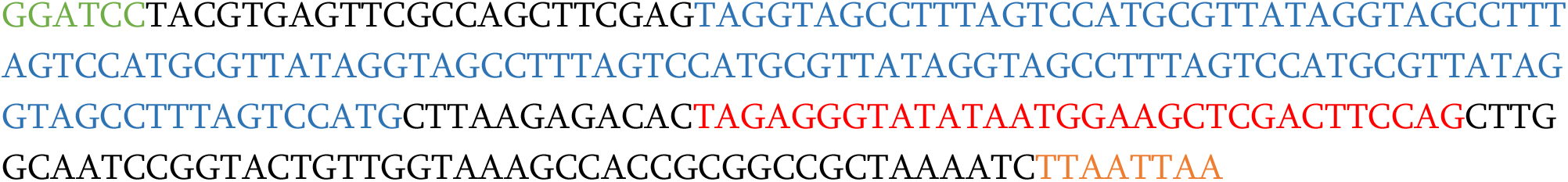

pVG10: *GALL* promoter

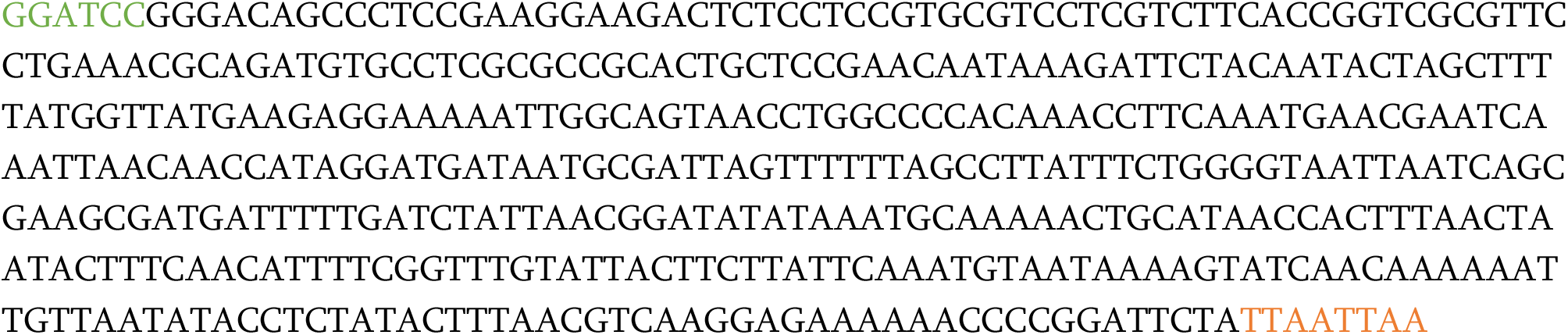

pVG11: *GLIP* (5 El222 binding sites, also known as pC120, surrounded by GAL1 promoter with Mig1 binding sites but without upstream activating sequence)

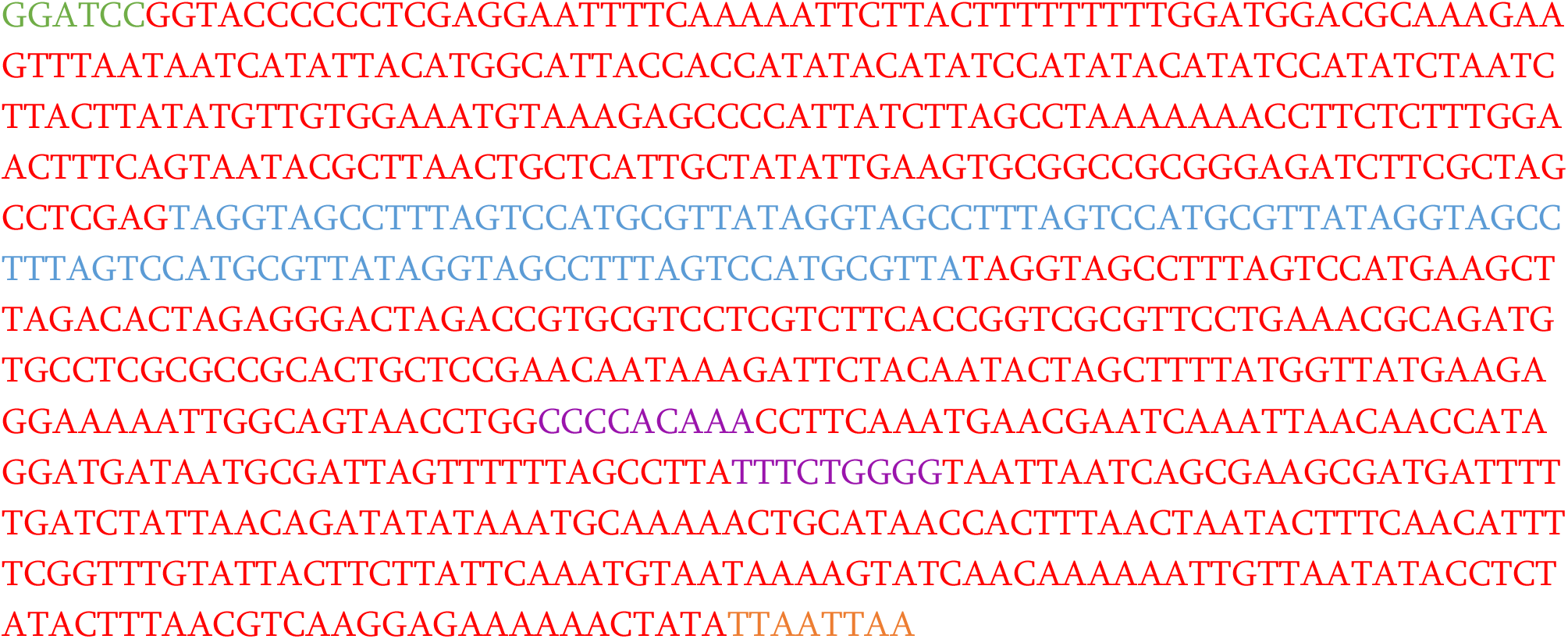

pVG45: *CUP1* promoter

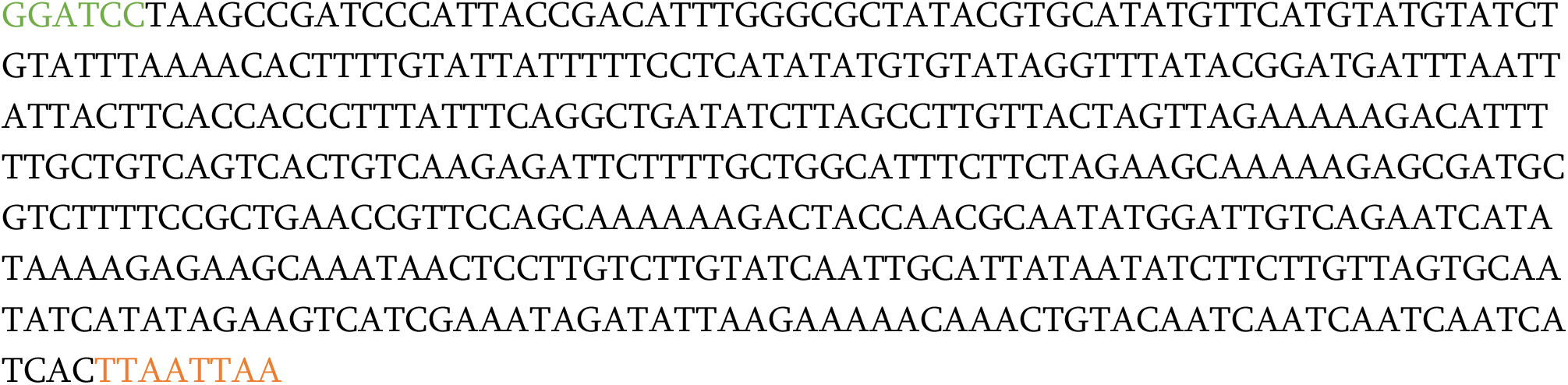

pVG46: *PHO5* promoter (cloned with BsWI and PacI restriction enzymes since there is a BamHI cutsite inside the PHO5 promoter)

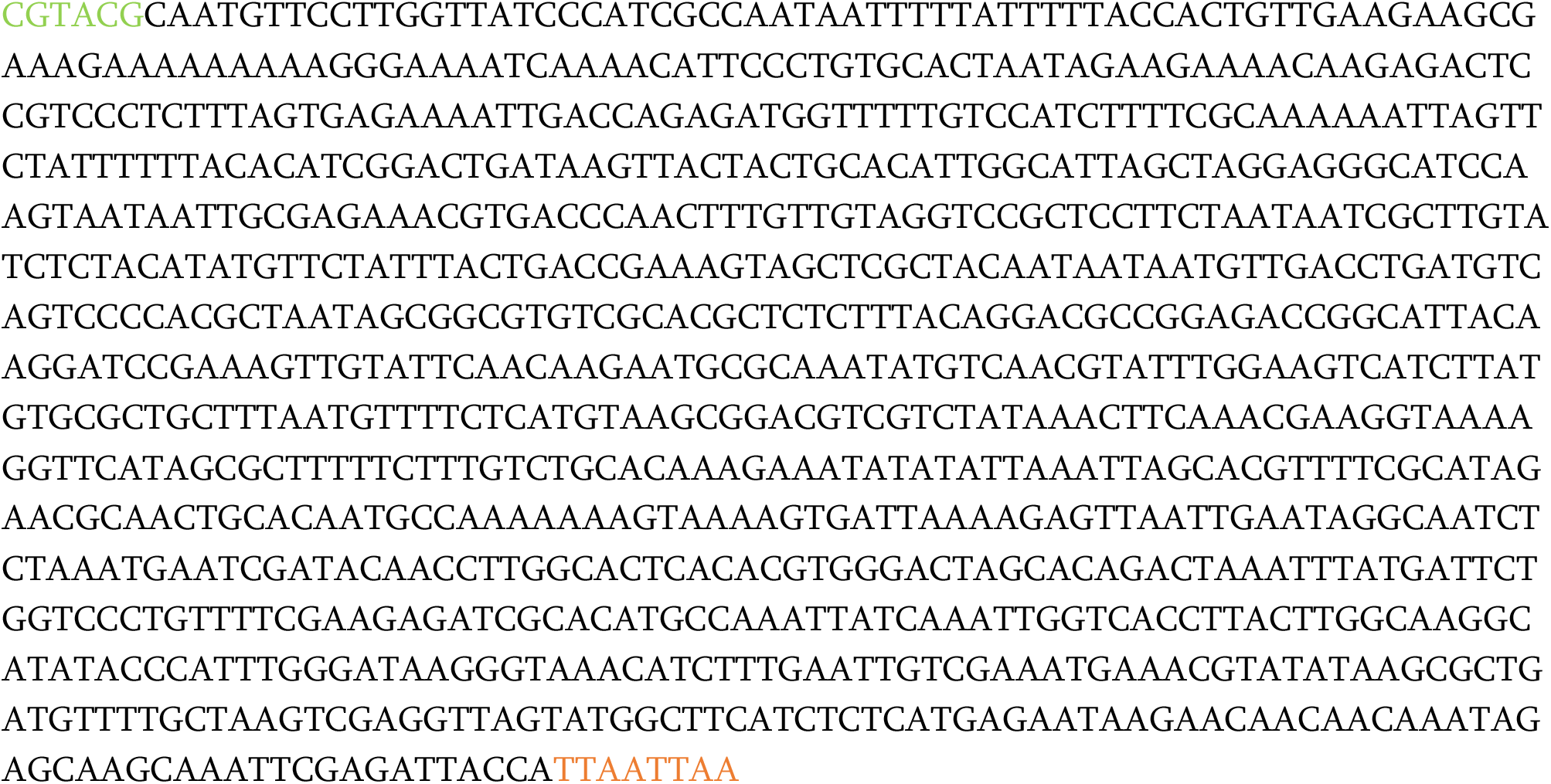

pVG47: *tetOpr* (based on tet operator sequence)

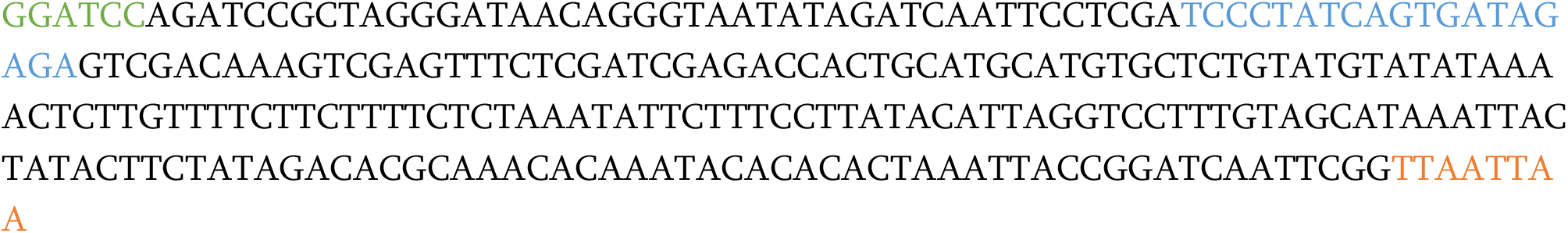

pVG49: *GAL1* promoter

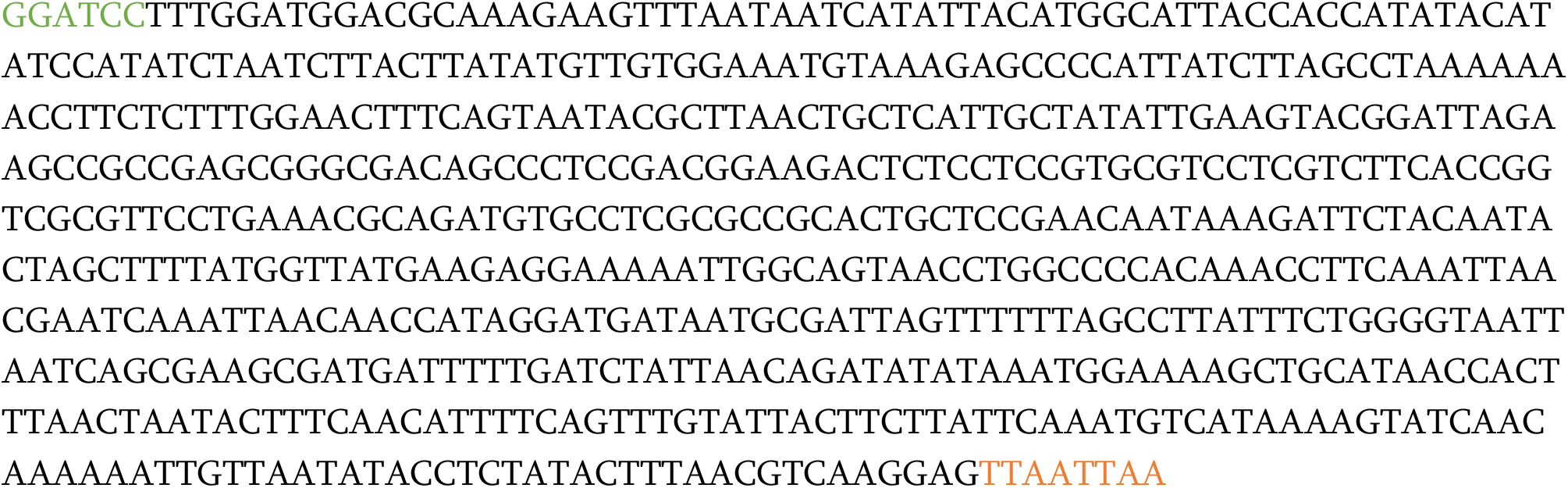

pCL10: *MET3* promoter

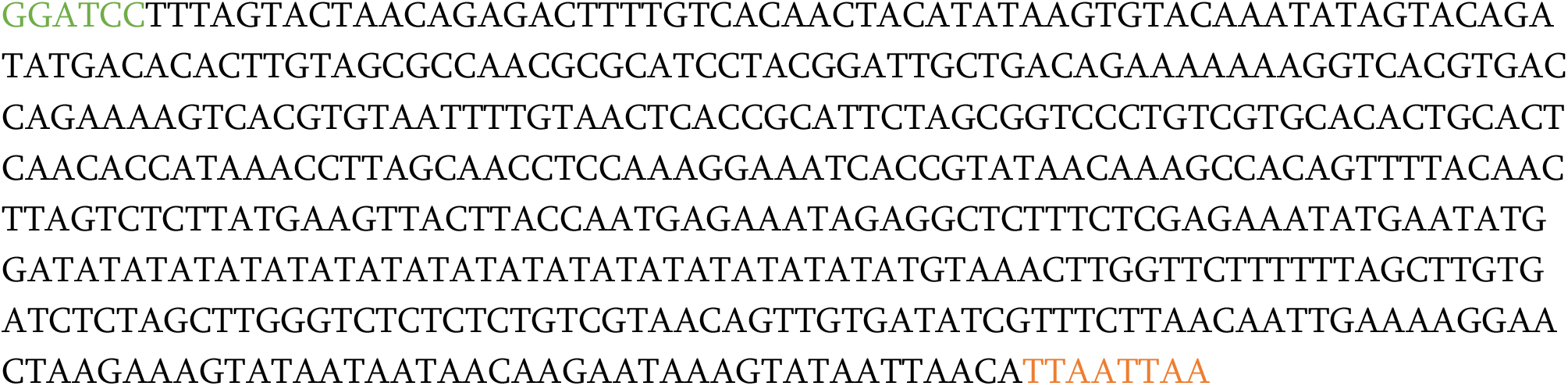

pVG88: *ARG3* promoter

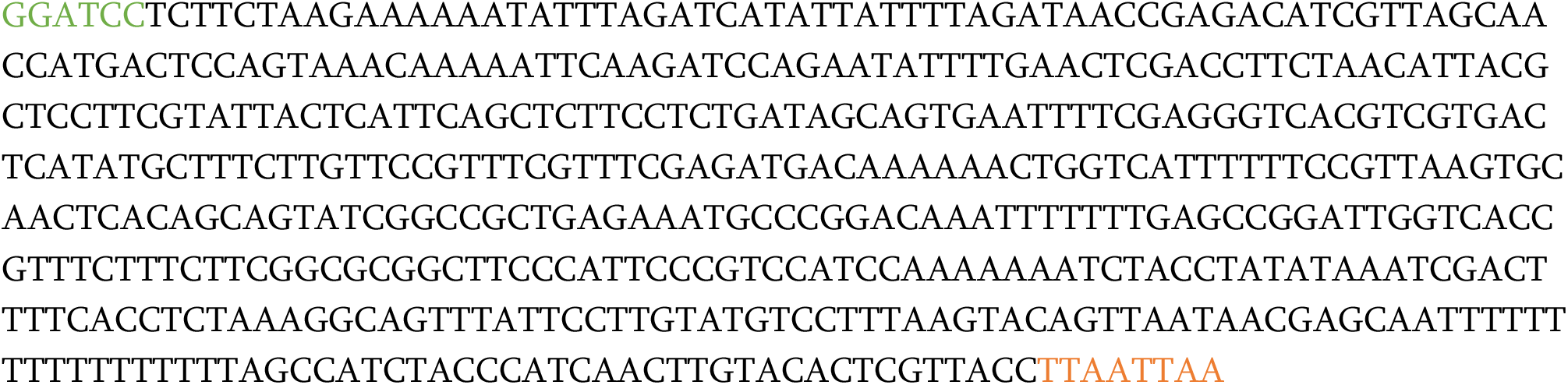

pVG107: Promoter used as a part of the Z_3_EV system

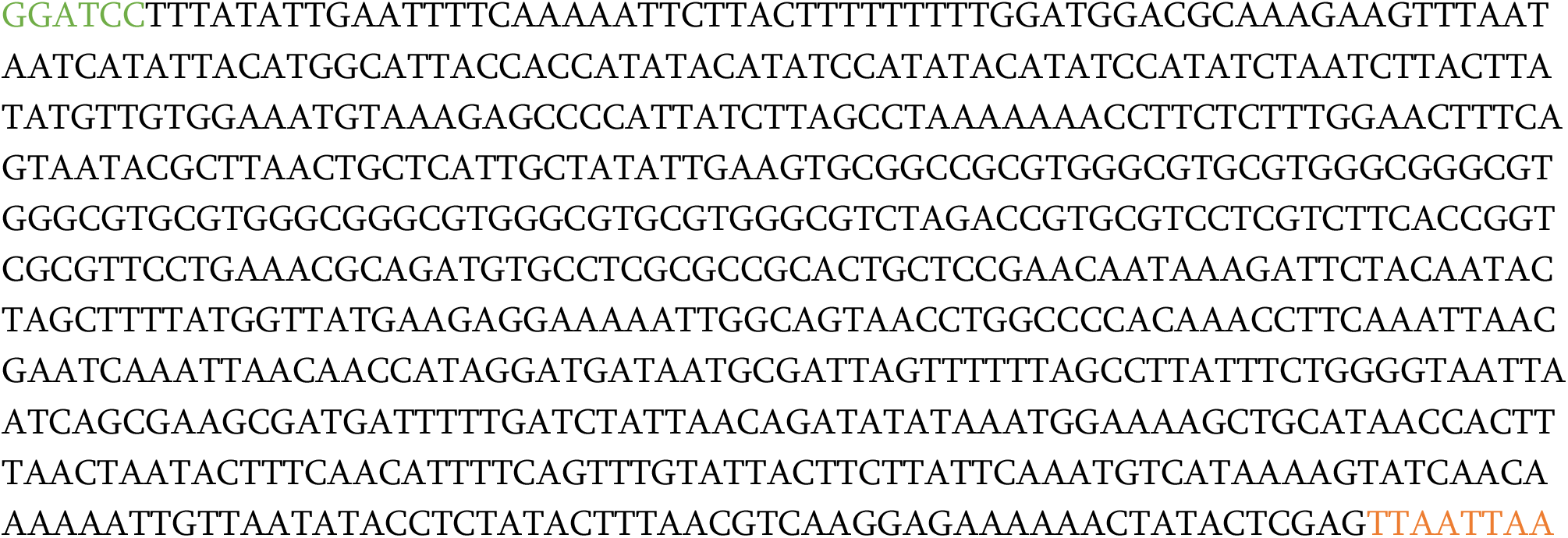

